# Precision Methylome and *in vivo* Methylation Kinetics Characterization of *Klebsiella Pneumoniae*

**DOI:** 10.1101/2020.12.04.409839

**Authors:** Jing Fu, Ju Zhang, Li Yang, Nan Ding, Liya Yue, Xiangli Zhang, Dandan Lu, Xinmiao Jia, Cuidan Li, Chongye Guo, Zhe Yin, Xiaoyuan Jiang, Yongliang Zhao, Fei Chen, Dongsheng Zhou

## Abstract

*Klebsiella pneumoniae* (*K. pneumonia*) is an important pathogen that can cause severe hospital-/community-acquired infections. To panoramically analyze *K. pneumoniae’s* methylation features, we completed the whole genome sequences of 14 *K. pneumoniae* strains covering various serotypes, multilocus-sequence typings (MLSTs), clonal groups (CG), viscosity/virulence and drug-resistances, and further characterized their methylomes using PacBio-SMRT and bisulfite technologies. We identified 15 methylation motifs (13 6mA and two 5mC motifs), among which eight were novel ones. Their corresponding MTases were further validated. Additionally, we analyzed the genomic distribution of G**A**<u>T</u>C and C**C**W<u>G</u>G methylation motifs shared by all strains, and identified differential distributive patterns of some hemi/un-methylated G**A**<u>T</u>C motifs tending to locate in the intergenic regions (IGRs). Specifically, we characterized the *in vivo* methylation kinetics at single-base resolution on a genome-wide scale by simulating the dynamic processes of replication-mediated passive demethylation and MTase-catalyzed re-methylation. The slower methylation-rates of the G**A**<u>T</u>C motifs in the replication origins (*oriC*) and IGRs suggest an epigenetic mechanism implicated in the regulation of replication-initiation and transcription. Our findings illustrate the first comprehensive dynamic methylome map of *K. pneumonia* at single base resolution, and provide an efficient means and important reference for a better understanding of epigenetic regulation in bacteria.

## Introduction

*Klebsiella pneumonia* (*K. pneumoniae*), the important member of the *Enterobacteriaceae*, can cause severe hospital- and community-acquired infections (*e. g*., pneumonia, genitourinary tract infection, and septicemia). There are some typing methods for *K. pneumoniae* strains, such as serotype, multilocus-sequence typings (MLSTs), and clonal group (CG) [1,2]. The related studies indicated that the hypervirulence phenotype usually corresponded to K1/K2/K57 serotypes and CG23-ST23 [1–3], while multidrug-resistance (MDR) phenotype often corresponded to CG258-ST11/ST258 [4].

Several studies on DNA methylation of *K. pneumoniae* strains using molecular biological techniques have identified three DNA methylases (MTases) and corresponding motifs, including two restriction-modification (R-M) systems (M.KpnI: GG<u>T</u>**A**CC; M.KpnBI: CAA**A**N_6_R<u>T</u>CA) and one orphan MTase (Dam: G**A**<u>T</u>C) [5–7]. Further research on Dam indicated the epigenetic mechanism in regulating mismatch repair, virulence and pathogenicity of *K. pneumoniae* strains [8].

Recent rapid progress on high-throughput sequencing techniques, such as Pacific single-molecule real-time (SMRT) sequencing for accurate detection of modified bases (mainly 6mA) on a genome-wide scale and bisulfite sequencing for efficient analysis of genome-wide 5mC [9,10], has greatly facilitated investigations of DNA methylome in bacteria. It is well known that 6mA and 5mC are the two most important types of DNA methylation in prokaryote [9]. To date, many bacterial methylomes have been precisely determined using the above two techniques, including *E. coli*, *Mycoplasma genitalium*, *Bifidobacterium breve*, *Clostridium difficile*, *Campylobacter jejuni*, *Helicobacter pylori* [9–13] and *Mycobacterium tuberculosis* complexes (MTBC) reported by our group [14].

Through precisely and comprehensively analyzing the bacterial methylome, a lot of valuable information has been revealed, including methylation motifs and their corresponding methyltransferases, motif distributions on genomes, and the related epigenetic regulation mechanisms in bacteria [9,15,16]. Most identified MTases and the corresponding motifs belong to R-M system, which primary function is to prevent (cleave) the invading DNA and protect genomic DNAs through methylation-mediated mechanism [9–14]. Distinctively, some orphan MTases (no cognate restriction enzymes) and the corresponding motifs exert multiple epigenetic regulation functions in bacteria [15–18]. Among them, Dam/G**A**<u>T</u>C motif is the most well-known because of its prevalent existence in almost all the *Enterobacteriaceae* bacteria and involvement in the epigenetic regulation of replication, transcription and mismatch repair [15–17, 19–21]. In particular, its regulatory role in replication initiation has been studied in *E. coli*. Its *oriC* region contains five DnaA boxes and 11 G**A**<u>T</u>C motif sites. The replication-mediated passive demethylation causes the hemi-methylated G**A**<u>T</u>C motifs adjacent to the DnaA boxes that are specifically recognized and bound by SeqA, further leading to the competitive occupation of the motif sites between Dam and SeqA [17]. As a result, the re-methylation of the motifs is delayed, which in turn prevents the initiation cascade for chromosome replication induced by *DnaA* protein [17,19]. The re-methylation of the upstream G**A**<u>T</u>C motifs of the third and fifth *DnaA* boxes are the rate-limiting steps for DNA replication initiation in *E.coli* strains [17]. In addition, Dam also participates in the transcriptional regulation of the downstream genes (*e. g., opvAB* in *Salmonella enterica)* [15].

Although several MTases and corresponding motifs have been revealed in *K. pneumoniae* strains, the whole methylome has not been reported so far. Here, we obtained whole-genome sequences of *K. pneumoniae* strains of 14 various types, and then characterized their methylomes using SMRT combined bisulfite sequencing techniques. Fifteen methylation motifs were identified (13 6mA and two 5mC methylation motifs), including eight novel ones corresponding to eight novel MTases termed as *kamA ~ G* (*K. pneumoniae* adenine methyltransferase A ~ *G*) and *kcmA* (*K. pneumoniae* cytosine methyltransferase A). We further analyzed the distribution pattern of the G**A**<u>T</u>C and C**C**W<u>G</u>G methylation motifs shared by all *K. pneumoniae* strains. Importantly, by establishing a mathematical model to simulate the dynamic process of passive demethylation and re-methylation of each motif in the exponential phase, we characterized the genome-wide *in vivo* methylation kinetics at single-base resolution. The motifs at different genomic locations showed different re-methylation rates, and the G**A**<u>T</u>C motifs in *oriC* region and intergenic regions (IGRs) had slower re-methylation rates. Our findings indicate potential roles of epigenetic regulation in the replication initiation and transcription of *K. pneumonia* genome, and provide important reference and insights into our better understanding of *K. pneumonia* epigenomics.

## Results

### General bioinformatic analysis of 14 *K. pneumoniae* strains

We first obtained the whole genome sequences of 14 *K. pneumoniae* strains with various serotypes, MLSTs, CGs, viscosity/virulence and drug-resistances (Table S1-3) by SMRT sequencing followed by correction using Illumina sequencing (**Figure 1**). The genome data have been deposited in National Genomics Data Center (CRA003482) and NCBI (PRJNA477755) for strains of NTUH-K2044, 11492, 11420, 11454, 12208, 11311, 23, 11305, N201205880, 309074, 13190, 283747, 721005, 11021, and 11305. We then constructed a phylogenetic tree using 76 complete genomes of *K. pneumoniae* strains (14 from ours and 62 from online) (Figure S1, Table S4). Our 14 *K. pneumoniae* strains covered many common CGs and MLSTs of *K. pneumoniae* strains in China (*e. g*., CG23-ST23 and CG258-ST11) [22], indicating the well representativeness yet diversity of *K. pneumoniae* strains selected in our study.

**Figure 1.**
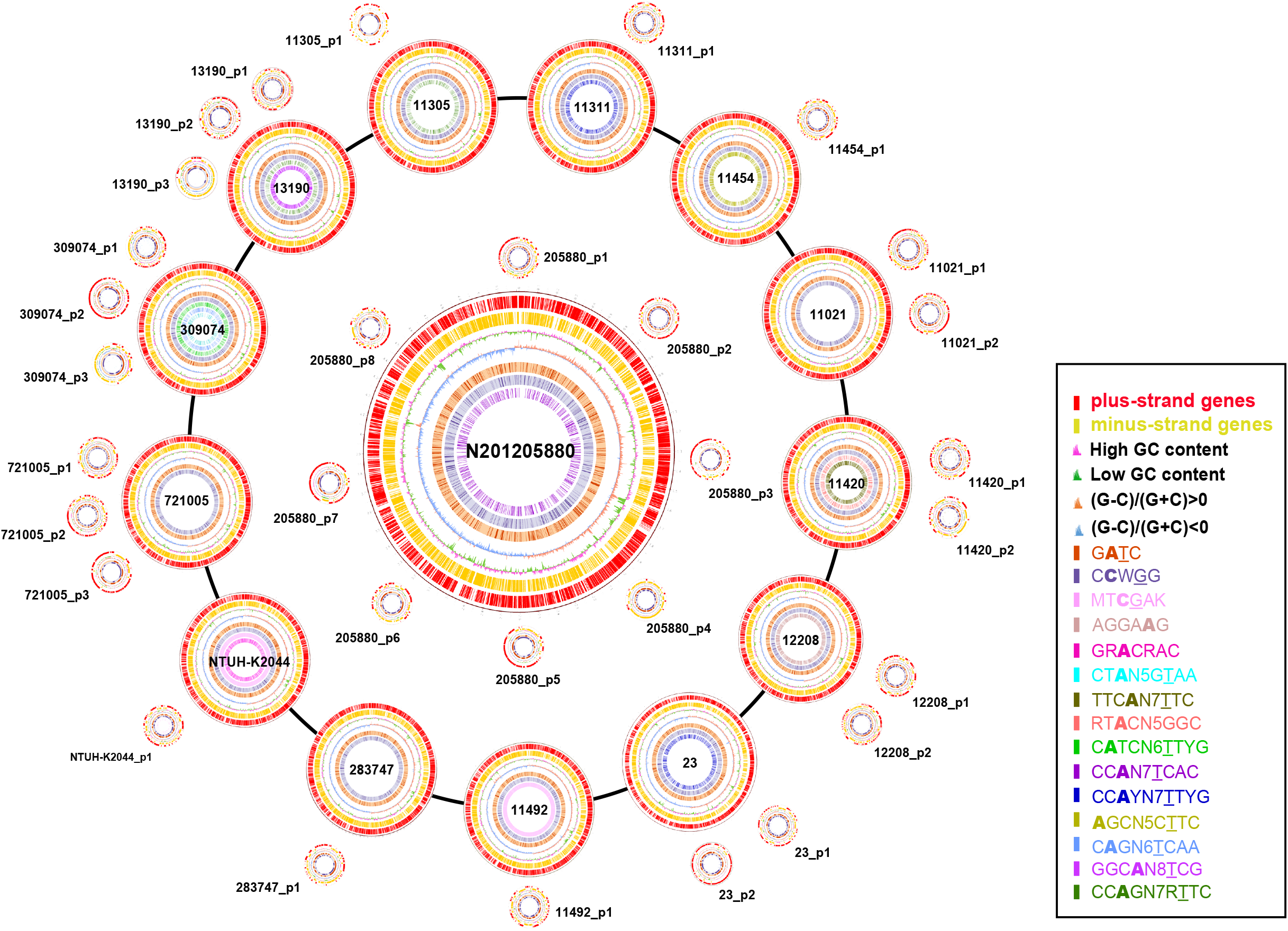
Circos plots displaying general genomic information of 14 *K. pneumoniae* strains. The circles are as follows (from outside to inside): (1) A physical map scaled in megabases (Mb) from the base 1, which is the start of putative replication origin; (2) Coding sequences transcribed in a clockwise direction; (3) Coding sequences transcribed in a counterclockwise direction; (4) G+C content in a window of 2 kb sliding (red and green indicates G+C content that is higher and lower than average, respectively); (5) GC (G-C/G+C) skew in a window of 2 kb sliding (orange and green indicates GC skew above and below zero, respectively); (6) G**A**<u>T</u>C motifs; (7) C**C**W<u>G</u>G motifs. The other circles inside indicate the other 13 methylation motifs.

The bioinformatic analyses provided the general genome information (**Table 1**, Figure 1), including genome size (5.20-5.54 Mb), GC content (57.4-57.9%), predicted protein-coding gene number (4990-5485), gene length (886-922 bp), and the percentage of coding region (88.26-90.23%) [23]. In addition, each *K. pneumoniae* strain contained 1-8 plasmids, with lower GC content (47.27-54.85%), lower percentage of coding region (75.98-88.63%), and shorter average gene length (596-831 bp) (Table 1).

**Table 1.** General genomic information of the 14 *K. pneumoniae* strains.

Additionally, the relatively conserved genomic sequences and structures among the 14 *K. pneumoniae* strains were also indicated: ANI analysis revealed more than 99% identity among the 14 *K. pneumoniae* strains; no extensive translocations, duplications or inversions were found in the *K. pneumoniae* genomes except for strains of 11492 and 11454 each containing a large inverted fragment on the genomes (Figure S2).

### Seven known and eight novel methylation motifs and corresponding methyltransferases in *K. pneumoniae* strains

A total of 13 6mA and two 5mC methylation motifs were identified in all *K. pneumoniae* strains (**Table 2-4**) by SMRT and bisulfite sequencing techniques (Table S2, S5), including seven known and eight novel methylation motifs (**Table 5**, Table S6-7). It is worth noting that G**A**<u>T</u>C and C**C**W<u>G</u>G motifs were shared by all strains. The other motifs were shared by at most two *K. pneumoniae* strains (Table 2-4). Further analyses indicated the relationship between the motifs and stain types (serotype, MLST and CG). The MT**C**<u>G</u>AK motif existed in NTUH-K2044 and 11492 belonging to K1 serotype and ST23-CG23, the common types of hypervirulent *K. pneumoniae* strains. The CC**A**YN_7_<u>T</u>TYG motif was shared by two strains (11311 and 23) of hypervirulent serotype K57 and ST412. The CC**A**GN_7_R<u>T</u>TC motif was present in strains 11305 and 13190 belonging to MDR CG147.

**Table 2.** Three types of modification patterns of GA<u>T</u>C motif from the 14 *K. pneumoniae* strains.

**Table 3.** Modification patterns of the 12 motifs with 6mA among the 14 *K. pneumoniae* strains.

**Table 4.** Modification patterns of the two motifs with 5mC among the 14 *K. pneumoniae* strains.

**Table 5.** 15 methylation motifs and corresponding DNA MTases from the 14 *K. pneumoniae* strains.

Modification analysis indicated that not all motif sites were fully-methylated (methylated on both strands, Table 2-4). A minority of motif sites (< 30%) were detected as being hemi-methylated (methylated on one strand only) or un-methylated within the *K. pneumoniae* genomes. The only exception was the MT**C**<u>G</u>AK motif in the NTUH-K2044 and 11492 genomes showing over half of hemi/un-methylated sites (56.03-56.77%). Further analysis indicated that the un-methylated MT**C**<u>G</u>AK motif prefered to have a guanine (G) in front of the motif (Figure S3, Table S8).

To search for the responsible MTases, we first predicted 22 MTases that might be responsible for the 15 methylation motifs from *K. pneumoniae* genomes in the REBASE database [24]. Among them, seven MTases and their corresponding motifs had been verified in previous studies (Table 5). To determine the MTases responsible for the eight newly detected methylation motifs, we performed the restriction digestion and SMRT/bisulfite sequencing using the plasmids containing the predicted MTase genes in methyltransferase-free *E. coli* ER2796. Crossover validations identified the corresponding eight MTases that could specifically recognize and methylate the respective eight novel motifs (Table 5, Table S6-7, Figure S4-6).

We further analyzed the distribution of 15 MTase genes on the genomes of the *K. pneumoniae* strains. Thirteen genes were located on the chromosomes and two others were located on the plasmids (**Figure 2**). In addition, within the 15 identified MTases, there were 10 Type I MTases, three Type II MTases and two classical orphan MTases (Dam and Dcm). Here *dam* and *dcm* genes were carried by all *K. pneumoniae* strains, and responsible for the methylation of G**A**<u>T</u>C and C**C**W<u>G</u>G motif, respectively. Among the three newly identified Type II MTases, KamC and KamD were predicted to be Type IIG enzymes, which function of endonuclease and methyltransferase were executed by a single gene (Table 5, Table S7).

**Figure 2.**
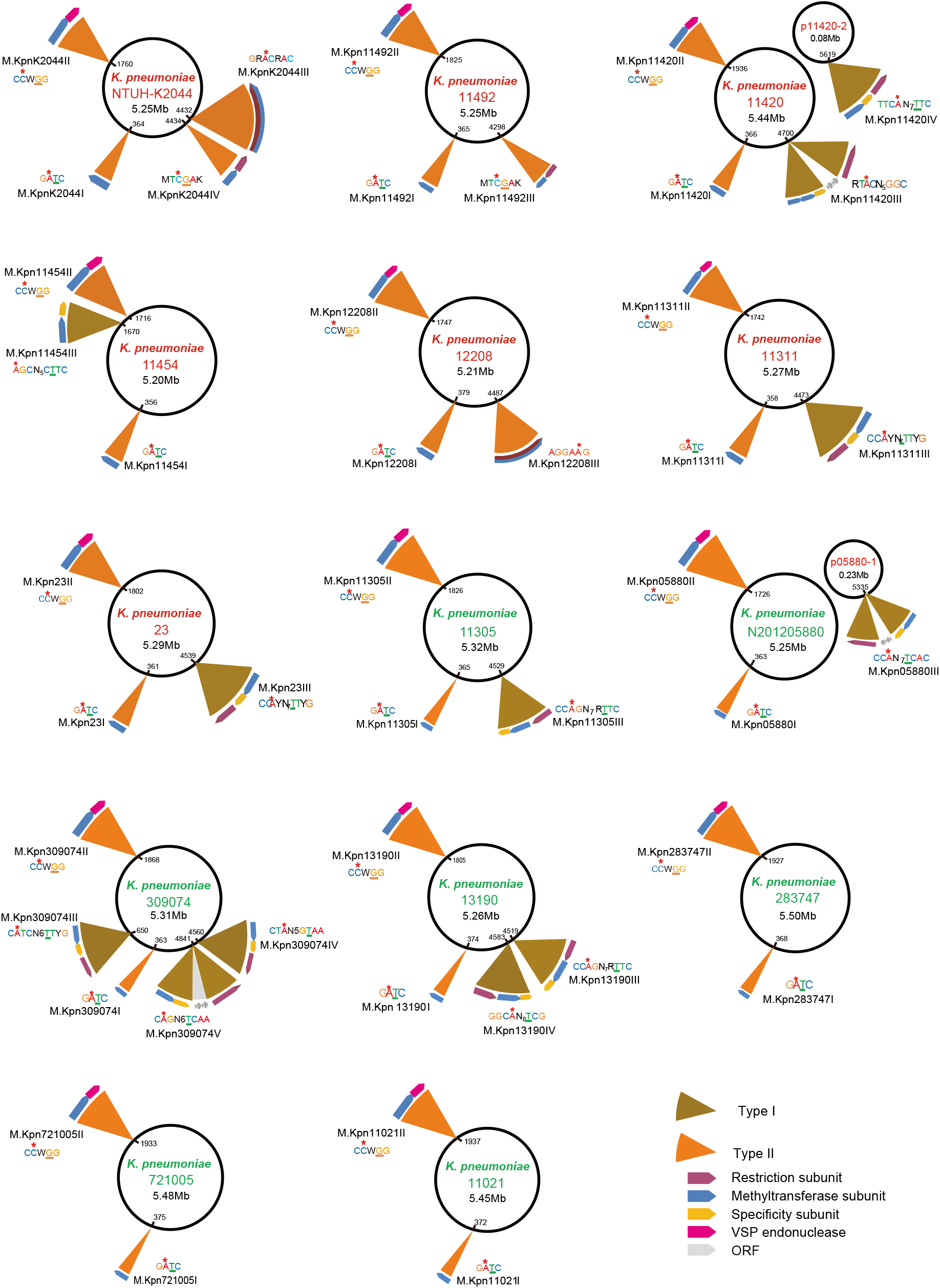
The 15 MTase genes and corresponding methylated sequence motifs for the 14 *K. pneumoniae* strains. Hyper-virulence and low-virulence strains are labeled in red and green letters, respectively. The large and small circles represent chromosome and plasmid genomes, repectively.

### Nonrandom distributions of GA<u>T</u>C and CCW<u>G</u>G motifs on *K. pneumoniae* genomes

Among the 15 methylation motifs, the G**A**<u>T</u>C and C**C**W<u>G</u>G had the most extensive distributions in all of the 14 *K. pneumoniae* strains, each containing ~30,000 G**A**<u>T</u>C and ~20,000 C**C**W<u>G</u>G modified sites (Table 2, 4). The distribution features of the above two motifs on the genomes of *K. pneumoniae* strains were further analyzed, and differential/uneven distribution on the genomes were observed (Figure 1). Both motifs exhibited some high-density and low-density regions on the genomes, where the genes were clustered into different COG functional categories (Figure S7). Importantly, the G**A**<u>T</u>C motif showed the highest distribution density (~34 sites/kb) in the *oriC* region (average density: 5-6 sites/kb) of the 14 *K. pneumoniae* genomes (Figure S8). In contrast, the C**C**W<u>G</u>G motifs didn’t display such enrichment in the *oriC* region.

We then compared the density distributions of G**A**<u>T</u>C and C**C**W<u>G</u>G motifs on the 14 *K. pneumoniae* genomes and the simulated genome with the same base composition (**Figure 3**, Figure S9-10). The result revealed that their distribution densities on the *K. pneumoniae* genomes (1kb consecutive window) were higher than those on the simulated genome, indicating the high-density/nonrandom distribution of these two motifs on the *K. pneumoniae* genome. To explore the underlying causes for their high-density distributions, we investigated the impact of selection pressure on these two motifs by calculating the Ka/Ks ratios (the ratio of nonsynonymous substitutions (Ka) to synonymous substitutions (Ks)) [25] of the corresponding fragments in gene regions (GRs) (more than 90% of the two motifs were located in GRs) (Figure S11). We observed that the amino acid (AA) codons with two motifs (two AA codons for G**A**<u>T</u>C motif, two or three AA codons for C**C**W<u>G</u>G motif) were under strong negative/purifying selection with Ka/Ks ≈ 0.09/0.09 in relative to the ratio of 0.39/0.54 for the scramble sequences in GRs.

**Figure 3.**
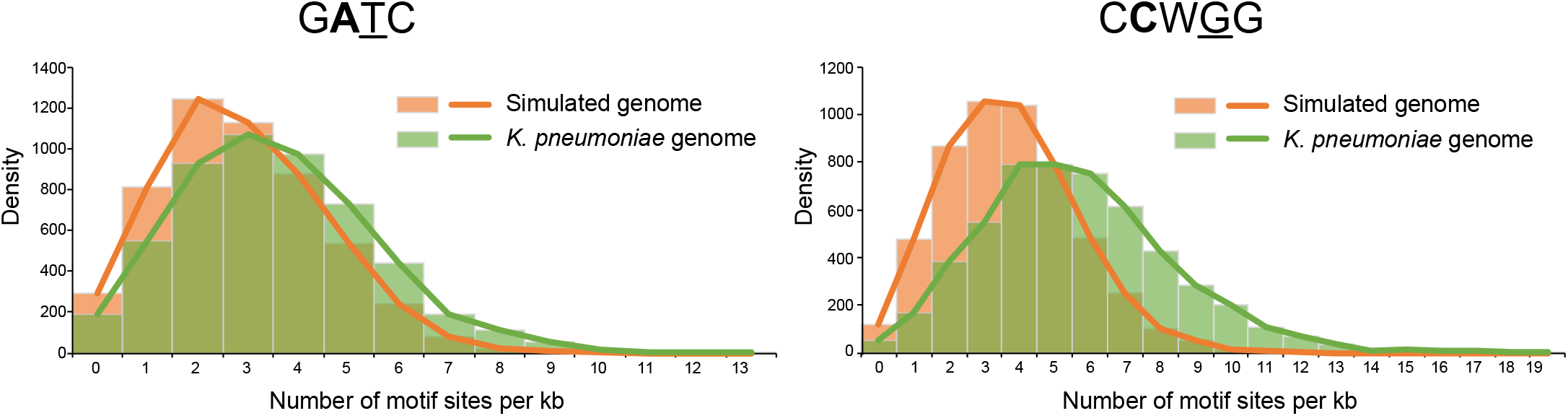
Density distribution of the GA<u>T</u>C/CCW<u>G</u>G motifs on the NTUH-K2044 and random generated genomes. The green histograms show the density distribution of G**A**<u>T</u>C/C**C**W<u>G</u>G on the NTUH-K2044 genome, which also follow Poisson distributions with λ = 5.64/3.62. The orange histograms show the density distribution of G**A**<u>T</u>C/C**C**W<u>G</u>G motifs on the random generated genome, which follow Poisson distribution with λ = 3.73/2.89 (total number of G**A**<u>T</u>C/C**C**W<u>G</u>G motifs × 1000 per genome size).

### Differential distribution patterns of the methylated, hemi-methylated and un-methylated GA<u>T</u>C and CCW<u>G</u>G motifs on *K. pneumoniae* genomes

We identified three methylation patterns (methylated, hemi-methylated and un-methylated) for the G**A**<u>T</u>C and C**C**W<u>G</u>G motifs. Most G**A**<u>T</u>C (84.26-98.13%) and C**C**W<u>G</u>G (77.58-98.84%) sites were detected as methylated state (Table 2, 4), whereas a small percentage of motif sites were hemi-methylated (< 15%) or un-methylated (< 10%) on the *K. pneumoniae* genomes. Further analysis demonstrated that the ratio of hemi-methylated G**A**<u>T</u>C motifs were (~6.48%) much higher than that of un-methylated G**A**<u>T</u>C motifs (~0.38%), while the hemi-methylated C**C**W<u>G</u>G motifs (~3.60%) showed a similar proportion as the un-methylated ones (~2.94%).

We then investigated the distribution ratio of methylated, hemi-methylated and unmethylated G**A**<u>T</u>C and C**C**W<u>G</u>G motifs in GRs and IGRs. The hemi/un-methylated G**A**<u>T</u>C motifs preferred to locate in IGRs, since its ratio (7.14%/34.11%) in IGRs was significantly higher than that of methylated G**A**<u>T</u>C motifs (5.23%) (*P* < 0.01, **Figure 4A**, Table S9). The analysis of the “fraction of methylated reads” (FRAC value) for the motifs in GRs/IGRs also supported the above findings that the hemi/un-methylated G**A**<u>T</u>C motifs were inclined to locate in the IGRs (Figure 4B). In contrast, the hemi/un-methylated C**C**W<u>G</u>G motifs (~7.49%) showed a little lower distribution ratio in IGRs compared with the methylated C**C**W<u>G</u>G motifs (~7.68%, Table S10).

**Figure 4.**
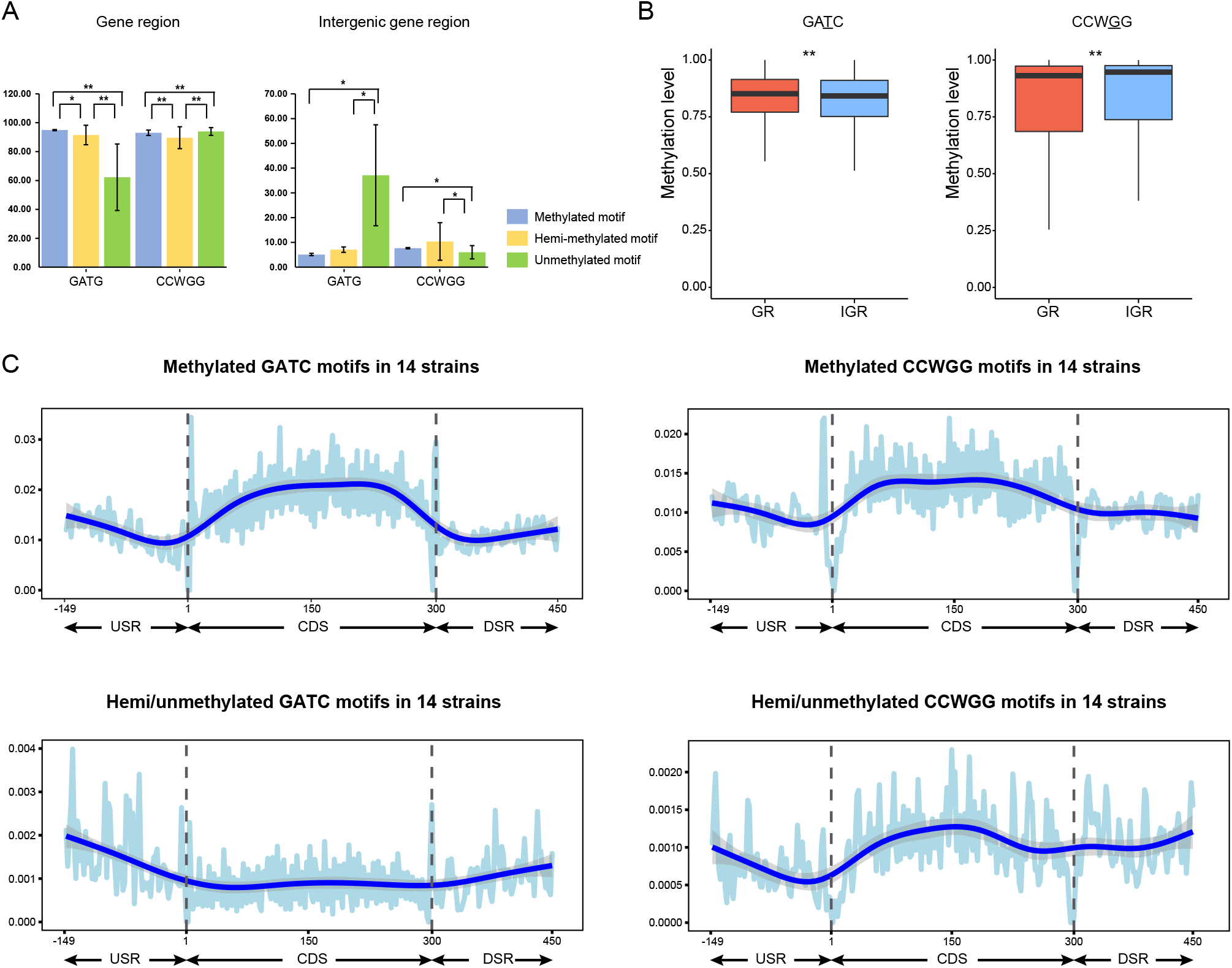
Distribution of GA<u>T</u>C/CCW<u>G</u>G motifs with different methylation patterns in GRs and IGRs. **A**. Bar plots showing the ratio of G**A**<u>T</u>C/C**C**W<u>G</u>G motifs with different methylation patterns in GRs and IGRs of 14 *K. pneumonia* strains. The blue, yellow and green bars indicate the ratios of methylated, hemi-methylated and un-methylated motifs in GRs and IGRs. **B**. Box plots showing the methylated levels of G**A**<u>T</u>C/C**C**W<u>G</u>G motifs in GRs and IGRs (red and blue boxes). **C**. Frequency distribution of the methylated (top) and hemi/un-methylated (bottom) G**A**<u>T</u>C/C**C**W<u>G</u>G motifs in GRs and IGRs of 14 *K. pneumonia* strains. The dotted grey lines represent the position of the start and stop codons. GRs, gene regions; IGRs, intergenic regions.

The analysis of the G**A**<u>T</u>C motifs’ density in 5’ upstream region (USR), coding sequence (CDS) and 3’ downstream region (DSR) also supported the above conclusion: the methylated motifs displayed a higher density in GRs, while the hemi-/un-methylated motifs presented a higher density in IGRs (both 5’ USR and 3’ DSR) (Figure 4C). We further explored the hemi/un-methylated G**A**<u>T</u>C sites shared in the 5’ USR regions of 14 *K. pneumoniae* strains: 13 hemi/un-methylated G**A**<u>T</u>C sites corresponding to 11 genes were observed (Table S11). We also detected 12 high-density hemi/un-methylated clusters (no less than three consecutive hemi/un-methylated motifs in at least two strains) in the 5’ USR regions (Table S12).

We then analyzed the sequence conservation of the DNA fragments (20 nt on both sides) containing hemi/un-methylated or methylated motifs to determine the causes leading to their differential distributions in IGRs. The results showed that the fragments with hemi/un-methylated G**A**<u>T</u>C motifs (44 nt) exhibit higher conservation than those with methylated G**A**<u>T</u>C motifs in IGRs (Figure S12).

### Methylation kinetic analysis revealing different re-methylated rates of GA<u>T</u>C and CCW<u>G</u>G motifs during growth cycle

Since there is no demethylase in bacteria, the *in vivo* methylation kinetics characterization is based on the dynamic equilibrium between the replication-mediated passive demethylation and MTases-catalyzed re-methylation [18]. To explore the features of methylation kinetics of G**A**<u>T</u>C and C**C**W<u>G</u>G motifs during the growth cycle, we first characterized the methylomes of the NTUH-K2044 and 11492 strains at the exponential phase (1 h), transition-to-stationary phase (4 h) and stationary phase (24 h) (**Figure 5A**, Table S13-14). By comprehensively analyzing genome-wide sequencing coverages and the fractions of methylated reads (methylated-read ratio/FRAC value), G**A**<u>T</u>C and C**C**W<u>G</u>G motifs were found to exhibit distinct kinetic features during the growth cycle (Figure 5B, Figure S13). As for G**A**<u>T</u>C motifs, the methylated-read ratios were more than 90% throughout the genomes in the three phases (Figure 5B), although the sequencing coverages varied on the genomes during the growth cycles (Figure S13), indicating that the G**A**<u>T</u>C motifs could be re-methylated in a very short time after the passive demethylation caused by replication: *i.e*., the re-methylated rate was almost identical to the passive demethylation rate.

**Figure 5.**
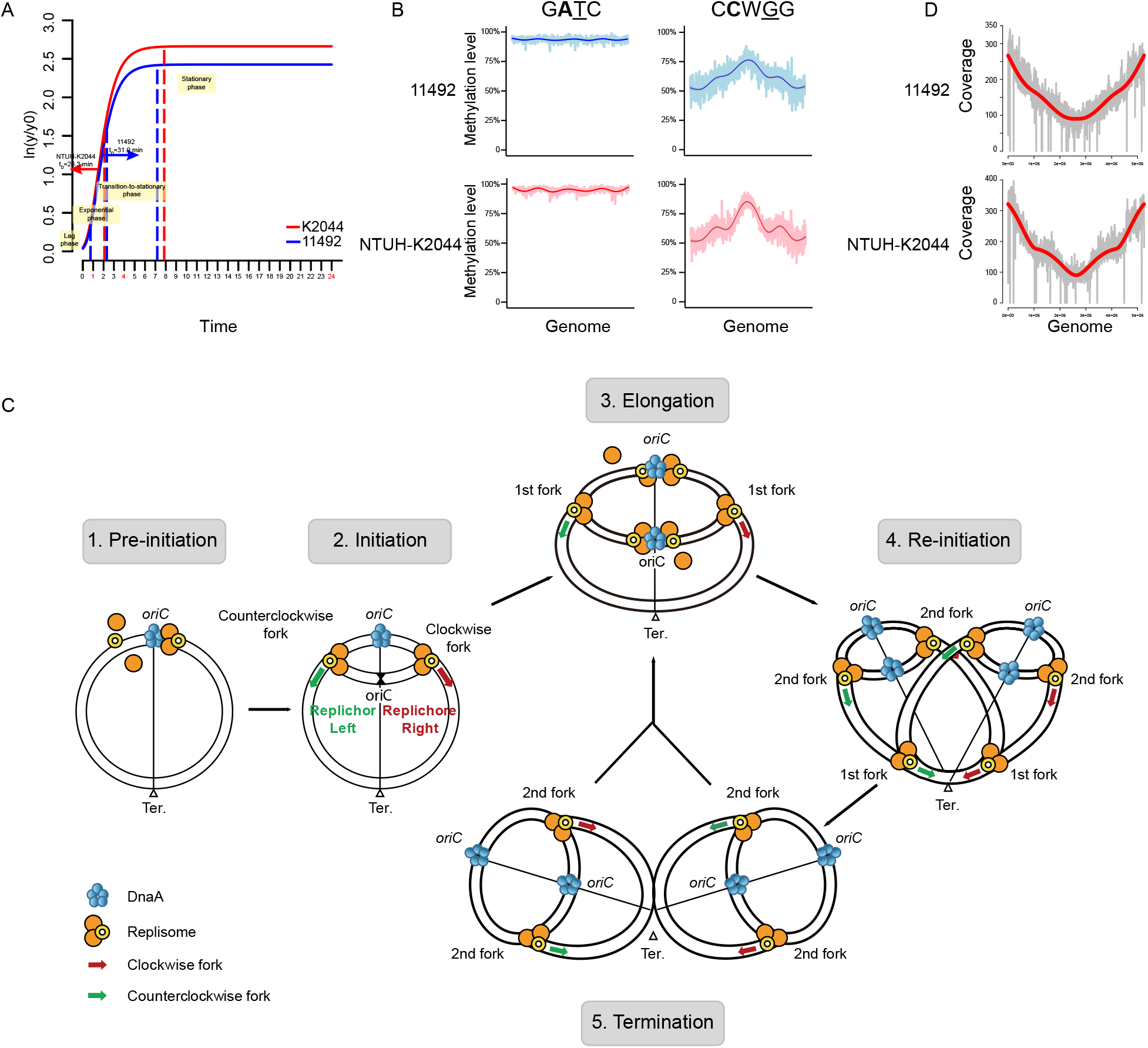
Dynamic methylation analysis of GA<u>T</u>C and CCW<u>G</u>G motifs of NTUH-K2044 and 11492. **A**. Growth curves of *K. pneumonia* NTUH-K2044 (red) and 11492 (blue). X-axis represents the growth time, including lag phase, exponential phase, transition-to-stationary phase and stationary phase. Y-axis represents the logarithms of normalized O.D. value. At 1, 4 and 24 h, aliquots of bacterial cultures were collected and sequenced using PacBio and Illumina platforms. The doubling time (tD) of the two strains are also labelled in the plot. **B**. Genome-wide methylation level versus genome position for the two motifs (G**A**<u>T</u>C and C**C**W<u>G</u>G) at exponential phase. The bold fitting lines approximate the average methylation levels across the genomes (5 kb window size). **C**. Schematic diagram showing the process of DNA replication in exponential phase. (1) The origin replication complex binds to oriC region. (2) The first round of replication will be initiated when original replication complex binds to *oriC* region. (3) The second round of replication begins before completion of the first round replication. (4) Six replication forks are generated in one bacterium. (5-6) The first round of replication completes, followed by a new cell replication cycle. **D**. Fitting of the genomic coverages with the mathematical model of replication. The genomic coverage plots of 11492 (upper) and NTUH-K2044 (lower) strains are shown in grey; the mathematically fitting curves are shown in red.

In contrast, the C**C**W<u>G</u>G motif showed a much slower re-methylation rate. First, in the exponential phase (1 h), the methylated-read ratio of C**C**W<u>G</u>G motifs in the *oriC* regions (55.66 ± 18.54%) was much lower than that in the replication termination (*Ter*) regions (82.63 ± 12.71%) (Figure S13B). Secondly, the average methylated-read ratio in the transition-to-stationary phase (54.50 ± 27.14%, 4 h) was close to that of the exponential phase (60.50 ± 17.71%, 1 h), but not the stationary phase (80.28 ± 19.05 %, 24 h) (Figure S13B). This finding suggested a much slower re-methylation rate compared to the replication-mediated passive demethylation rate for C**C**W<u>G</u>G motifs, which resulted in the above differences in the methylated-read ratios.

To quantify the re-methylation time per motif in the exponential phase, we first investigated the replication kinetics of the aforementioned two *K. pneumoniae* strains. Figure S13 showed that the genome sequencing coverage in the *oriC* regions was more than twice as much as that of the *Ter* regions (2.93 and 3.02 folds for the 11492 and NTUH-K2044 strains, respectively), indicating that the next round of replication was re-initiated before the replication termination in the exponential phase. We then constructed a replication model (Figure 5C) by fitting genome coverage data, and obtained the ratio (tD/tR) of doubling time (tD) to replication termination time (tR) (see “Materials and methods” for detail). The doubling time (tD) in the exponential phase also reflects the re-initiation time. Interestingly, we obtained the same tD/tR values (0.59) for the two *K. pneumoniae* strains (Figure 5D), suggesting a similar regulatory mechanism for replication cycle, which should be intrinsically carried by *K. pneumoniae* strains. Two tD values were then calculated through fitting the growth curves (~34.64 min for strain 11492; ~27.95 min for NTUH-K2044, Figure 5A), therefore we could infer the t_R_ from the t_D_/t_R_ values (~58.71 and ~47.36 min for strain 11492 and NTUH-K2044, respectively).

We further obtained the re-methylation time per motif (**Table 6**) through simulating the dynamic processes of passive demethylation and re-methylation in the exponential phase based on the five parameters, such as methylation-read fraction of each motif (*M_(x)_*), initial methylation-read fraction of each motif 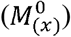, t_D_, t_R_ and distribution density of the first replication forks (*P*_(x_1_)_, see “Materials and methods” for detail).

**Table 6.** Average methylation time per motif in NTUH-K2044 and 11492.

In general, the mean re-methylation time of 6mA was shorter (3.52 and 3.46 min for G**A**<u>T</u>C and GR**A**CRAC motifs, respectively) than that of 5mC (9.23 and 4.55 min for C**C**W<u>G</u>G and MT**C**<u>G</u>AK motifs, respectively) in the exponential phase (Table 6). As for a certain motif, the re-methylation time in the NTUH-K2044 strain was shorter than that in the 11492 strain (Figure S14). In addition, the flanking bases could influence the re-methylation rates of the G**A**<u>T</u>C and C**C**W<u>G</u>G motifs. When the flanking bases were C/G rather than T/A, G**A**<u>T</u>C motif showed a faster re-methylation rate, which was totally reversed for the C**C**W<u>G</u>G motif (Table 6, Figure S15).

### Slower re-methylation rate of GATC motifs in intergenic and *oriC* regions at the exponential phase

To investigate the role of G**A**<u>T</u>C motifs in transcriptional regulation of *K. pneumoniae* strains, we analyzed the re-methylation rates of the motifs in IGRs (including 5’ USRs and 3’ DSR) at the exponential phase (**Figure 6A**). The results indicated the slower re-methylation motifs in IGRs (3.94 ± 5.82 min) than that in GRs (3.50 ± 5.10 min). Since most 5’ USRs in bacteria are overlapped with the promoter regions and involved in the transcriptional regulation [26], we further explored the COG functional category of genes with top 5% slow re-methylation sites (> 7.04 min/motif) in 5’ USRs (Table S15). Four enriched functional categories (“Cell cycle control, cell division, chromosome partitioning”, “Carbohydrate transport and metabolism”, “Intracellular trafficking, secretion, and vesicular transport” and “Translation, ribosomal structure and biogenesis”) were observed (Table S15), of which at least 5% of genes contained a slow re-methylation sites (> 7.04 min/motif) in 5’ USRs. Among them, the “carbohydrate transport and metabolism” functional category owned the most genes (22 genes) with slow re-methylation rates in 5’ USRs, including three hemi/un-methylated motifs shared in 14 *K. pneumoniae* strains and even one hemi/un-methylated motif-clusters.

**Figure 6.**
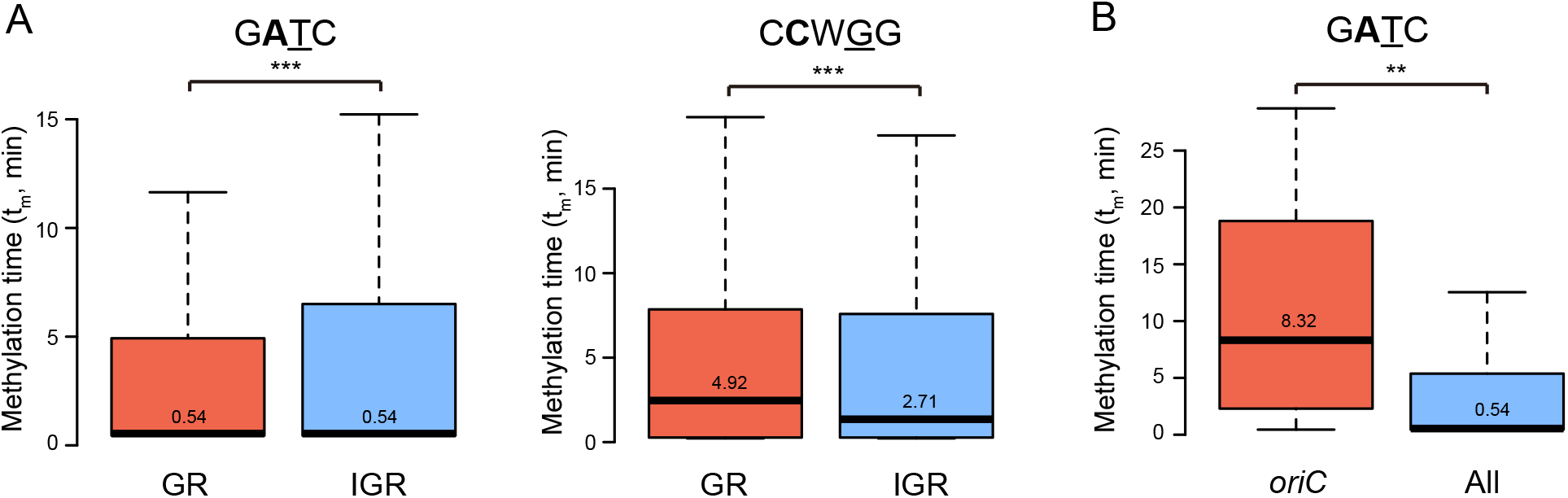
Comparison of re-methylation times of GA<u>T</u>C/CCW<u>G</u>G motifs in different genome regions. **A**. Box plot shows the methylation time of G**A**<u>T</u>C motifs in *oriC* and other regions. **B**. Box plot shows the methylation time of G**A**<u>T</u>C and C**C**W<u>G</u>G motifs in GRs and IGRs. ** *P* < 0.01, *** *P* < 0.001. GRs, gene regions; IGRs, intergenic regions.

To further explore the role of G**A**<u>T</u>C motifs in replication regulation of *K. pneumoniae* strains, we analyzed the re-methylation rates of the motifs in *oriC* regions (Figure 6B). The G**A**<u>T</u>C motifs in *oriC* region had the highest distribution density (~34 sites/kb) (Figure S8) and slower re-methylation rates (10.35 ± 8.69 min). Among the 14 G**A**TC motifs in *oriC* region, eight and seven motifs with slow remethylation rates (> 7.04 min/motif) were enriched in the upstream sequence of the fourth DnaA binding site (DnaA box) and the adjacent AT-rich region in strain 11492 and NTUH-K2044, respectively (Figure S16).

## Discussion

In this study, we characterized the precision methylome of 14 *K. pneumoniae* strains with different serotypes, MLSTs, CGs, viscosity/virulence, and drug-resistances, using SMRT and the bisulfite sequencing techniques, and identified 15 DNA methylation motifs (eight novel ones) corresponding to 13 R-M system MTases and two orphan MTases, respectively. Two motifs (G**A**<u>T</u>C and C**C**W<u>G</u>G) and their respective orphan MTases (Dam and Dcm) appeared to be the most important since they were present in all *K. pneumoniae* strains with the most extensive distributions on genomes in relative to other modified motifs (Table 2, 4). This feature was also reported in almost all members of the *Enterobacteriaceae* family [21,24]. Functional analysis in previous reports indicated that these two motifs, especially G**A**<u>T</u>C, could exert multiple functions including transcriptional regulation, cell-cycle controlling, and mismatch repair in *E.coli* and *Salmonella enterica* [15–17,19–21]. We also demonstrated their high-density distributions on *K. pneumoniae* genome comparing with that on simulated genome (Figure 3, Figure S9-10), which was probably caused by purifying selection (Figure S11) leading to their evolutionary conservation (Figure S12) as reported in other bacterial genome [27]. This well-conserved feature also implies an essential role of these two motifs in *K. pneumoniae*.

The hemi/un-methylated G**A**<u>T</u>C motifs were found to possess the tendency of IGRs (including 5’ USR and 3’ DSR) localization (Figure 4A, C). Since promoters in *K. pneumoniae* were also predicted to be located across the 5’ USR regions (Figure S17), thus, these hemi/un-methylated G**A**<u>T</u>C motifs in 5’ USR might be protected from the methylation by competitively binding of some regulators to the promoter regions [17,19]. In *E.coli* strains, this feature facilitates the epigenetic regulation of the downstream gene expression [28]. The status of these motifs suggested similar epigenetic mechanisms in *K. pneumoniae* strains, and might be the consequence of selection during long-term evolution (Figure S11), which was supported by our findings that the genomic fragments with hemi/un-methylated G**A**<u>T</u>C motifs in IGRs had higher sequence conservation (Figure S12).

Importantly, by establishing a mathematical model to simulate the dynamic process of passive demethylation and re-methylation of each motif in the exponential phase, we obtained the re-methylation time of each motif throughout the whole genome (Table 6, Figure S14). Our studies revealed that the motifs at different genomic locations showed different re-methylation rates. We could reasonably infer that the slower re-methylation rates in some sites/regions might also be due to the competitive binding of certain proteins in order to prevent methylation [17,19]. Thus, the slow re-methylation rates could precisely reflect the methylation-mediated epigenetic regulations at these sites/regions *in vivo*.

There are two methylation-mediated epigenetic regulations: transcription and replication, which were examined by methylation kinetic analysis in our study. Firstly, transcriptional regulation analysis indicated that the G**A**<u>T</u>C motifs in IGRs presented the slower re-methylation rates than those in GRs (Figure 6). This is in agreement with the distribution characteristics of the G**A**TC motifs in IGRs: lower methylatedread ratios and more hemi/un-methylated motifs (Figure 4A, 4B). As described above, most IGRs in bacteria overlap with the promoter regions and participate in the transcriptional regulation [26]. Thus, the promoter regions with slow re-methylation motifs should be the locations where the G**A**<u>T</u>C motifs with hemi/un-methylated status function as transcription regulators in *K. pneumoniae* strains. Similarly, this should also be the consequence of the competitive binding between Dam and some transcription regulatory proteins (such as OxyR) to these sites/regions [15,28,29], which is analogous to the epigenetic transcriptional regulation mediated by the competition between DNA methyltransferases and CTCF in CpG islands of eukaryotic cells [30]. Secondly, replication regulation analysis identified 7 - 8 slow G**A**<u>T</u>C motifs enriched in the upstream sequence of the fourth DnaA binding site (DnaA box) and the adjacent AT-rich region (Figure S16). SeqA has been shown to prefer to bind to these motifs for the purpose of lowering re-methylation rates [31] and preventing the initiation cascade for chromosome replication [17,31]. Therefore, re-methylation of these motifs in the fourth DnaA box and AT-rich regions should represent the main rate-limiting steps for triggering initiation of DNA replication in *K. pneumoniae* strains. Our findings also uncovered many promoter regions with slower re-methylation motifs in *K. pneumoniae* strains (Figure 6), therefore, it is reasonably speculated that epigenetic regulation in bacteria is very complex rather than simple as we believed.

Compared with G**A**<u>T</u>C motif, C**C**W<u>G</u>G motif has different distribution characteristics in GRs and IGRs (Figure 4A, B). We then performed the COG functional analysis for the genes with hemi/un-methylated C**C**WGG sites in IGRs, since previous studies have shown the epigenetic regulation of hemi/un-methylated sites in IGRs in bacteria [32]. Our findings showed that the top three enriched functional categories for the genes with hemi/un-methylated sites in IGRs were “Replication, recombination and repair [L]”, “Cell motility [N]”, and “Coenzyme transport and metabolism [H]” (Figure S18A), suggesting the possible epigenetic regulation of C**C**W<u>G</u>G sites in *K. pneumoniae*. On the other hand, they showed totally different COG categories for the genes with hemi/un-methylated G**A**TC sites in IGRs (Figure S18B), suggesting different epigenetic regulations between the two main methylated sites in *K. pneumoniae*.

We also explored the of methylation kinetic feature of C**C**W<u>G</u>G motifs during growth cycle. There were 1961 C**C**W<u>G</u>G sites changed in methylation during growth phase: 1954 hemi/un-methylated sites in exponential phase became methylated in stationary phase; seven unmethylated sites in exponential phase became hemimethylated in stationary phase. No methylated sites in exponential phase became hemi/un-methylated in stationary phase. The dynamic changes in methylation status might be due to the active and inactive replications in exponential and stationary phases [13]: active replication at exponential phase could result in more hemi/un-methylated sites on the genome because of the replication-mediated passive demethylation; inactive replication at stationary phases could lead to more methylated sites on the genome due to nearly total MTase-catalyzed methylation. Incidentally, the seven hemi-methylated C**C**W<u>G</u>G sites in stationary phase are located on the genes encoding for four small subunit ribosomal RNAs, two mobile element proteins and one possible transcription regulator (Table S16), which may be necessary for the survival of bacteria [33].

Importantly, eight novel MTases and related motifs were detected, including five Type I and three Type II MTases (Table 5, Table S6-7). These novel MTases and cognate REases on the genomes form restriction-modification (R-M) systems. It is known that R-M system could protect the bacterial cells by cleaving the foreign phage DNAs [9–14]. We then investigated the hemi/un-methylated sites in IGRs, since previous studies have shown the epigenetic regulation of hemi/un-methylated sites in IGRs of some bacteria [32]. Among the eight novel methylated sites, the MT**C**<u>G</u>AK motifs are shared by the NTUH-K2044 and 11492 strains, which owned the most hemi/un-methylated sites (56.03 and 56.77% for NTUH-K2044 and 11492 strains, Table 4). More than 60% of un-methylated MT**C**<u>G</u>AK motifs are shared by the two strains (Table S17), which are mainly enriched in “Transcription [K]”, “Inorganic ion transport and metabolism [P]”, and “Replication, recombination and repair [L]” COG categories (Figure S19). The CC**A**YN_7_<u>T</u>TYG motif is also owned by two strains (23 and 11311) (Table 3). Three hemi-methylated CC**A**YN_7_<u>T</u>TYG motifs were detected to be shared by the two strains, which were associated with *ascB, deoC*, and *fecA* genes, respectively (Table S18). These hemi/un-methylated MT**C**<u>G</u>AK and CC**A**YN_7_<u>T</u>TYG motifs in IGRs are shared by two strains, which might exert the epigenetic regulation of related gene expression. In addition, we also performed COG analysis for the other six newly reported methylated sites, which only existed in one *K. pneumoniae* strain (Figure S19). There is a limitation for these hemi/un-methylated sites in only one *K. pneumoniae* strain, since they may be derived from replication-mediated passive demethylation [34].

Previous studies on MTase kinetics mainly focused on *in vitro* analysis [35–37]. Several reported analyses about the *in vivo* methylation kinetics, including one publication about bacteria, only studied the overall, but not single-motif methylation kinetics [38–40]. Our study is the first to characterize the *in vivo* methylation kinetics at single motif resolution in whole genome, thus, offer an efficient means and valuable resources for a better understanding of epigenetic regulation in bacteria.

## Materials and methods

### Strain information, growth curve, and phenotypic characterization

The information of 14 *K. pneumoniae* strains used in this study was shown in Table S1. The strains were cultured overnight in Luria-Bertani (LB) medium at 37 °C. 1 ml of overnight culture suspensions was then transferred to a flask containing 200 ml of LB medium and cultured in a shaker at 200 rpm. The growth curve of bacteria was determined by recording OD_600_ values at different time-points. This experiment was performed in triplicate.

Drug susceptibility test was performed by VITEK 2 (bioMérieux, Durham, NC, USA), and the drug-resistance phenotypes were determined by Clinical and Laboratory Standards Institute (CLSI) standards (https://clsi.org/standards/products/microbiology/documents/m100/). The string test of *K. pneumoniae* strains was conducted, and hypermucoviscosity was defined by viscous strings with a length of more than 5 mm [41].

### Genomic DNA Extraction, sequencing, assembly, correction and annotation

The genomic DNA was extracted using TIANamp Bacteria Genomic DNA Kit (catalog No. DP302, TIANGEN, Beijing, China). The whole-genome sequencing was implemented by PacBio RSII platform using P6/C4 chemistry (Pacific Biosciences, CA, USA). Each strain was sequenced with 1-2 SMART cells with genome coverage of more than 50× (Table S2).

*De novo* assembly of the genome was performed using Hierarchical Genome Assembly Process 3 (HGAP3) in SMRT Portal (version 2.2.0). Gap closing was completed by PBJelly [42]. On the basis of Blast results, genome circularization was finished by manually removing the contigs overlapped regions.

To correct the polymer-errors, we re-sequenced the strains using Illumina sequencing (Table S3). Paired-end libraries were prepared as previous description, and clean reads were obtained after eliminating redundant and low-quality raw reads. Paired reads were extracted and then mapped onto the assembled genome sequences to obtain unique mapped reads using BWA. Pilon v.1.13 was subsequently used to polish genome sequences using unique map reads.

Genome sequences were annotated in the Subsystem Technology (RAST) pipeline with Rapid Annotation [43]. Unannotated genes were then predicted by alignment in the NCBI non-redundant (NR) database using BLAST. The protein functions were annotated by Clusters of Orthologous Groups (COG). tRNAs and rRNAs were predicted by tRNAscan-SE and “search_for_rnas” tools, respectively.

### Genome structure and phylogenetic analysis

The average nucleotide identity (ANI) and coverage were calculated by ANI on EzGenome (http://www.ezbiocloud.net/tools/ani) and online BLAST (https://blast.ncbi.nlm.nih.gov/Blast.cgi). Multiple alignments of genomic sequences were performed using the Mauve multiple alignment software [23].

Single nucleotide polymorphisms (SNPs) were detected by MUMmer based on the 14 genomes in this study and 62 published genomes (Table S4) using HS11286 as reference. Prank [44] was used to annotate the protein coding genes of the 76 *K. pneumonia* genomes, and then used Roary to predict 3173 core genes [45]. The SNPs (117,142 SNPs) in core genes were used to construct a phylogenetic tree on the basis of maximum likelihood using FastTree [46], followed by decoration using software evolview v2 (http://www.evolgenius.info/evolview/) [47].

### Genome-wide detection of 6mA and the related motifs using the SMRT sequencing data

The SMRT Portal (version 2.2.0) was applied to detect genome-wide 6mA modification and related motifs using the standard settings in the RS_Modification_and_Motif_Analysis. 1 protocol as previously described (https://smrt-analysis.readthedocs.io/en/latest/SMRT-Pipe-Reference-Guide-v2.2.0/). We then identified the motifs through selecting the top 1000 kinetic hits and subjected a window of ± 20 base around the detected modified base to MEME-ChIP [48], followed by comparing them with the predicted MTase targeting motif sequences in REBASE [24].

There are three methylated patterns of motifs: methylated, hemi-methylated and un-methylated motifs. The methylated/hemi-methylated/un-methylated motifs indicate the sites with methylated nucleotides on both/one/no strands.

### Genome-wide detection of 5mC and the related motifs using the bisulfite sequencing data

The 5mC methylation was detected by bisulfite sequencing (Table S5). Trimmomatic (v0.32) [49] was used to trim adapters and low-quality bases using default parameters. Clean reads were mapped on the reference genomes by Bismark (version 0.12.2) [50]. We then identified the motifs through subjecting a window of ± 20 base around the detected modified base to MEME-ChIP [48], followed by comparing with the predicted MTase targeting motif sequences in REBASE [24].

### MTases cloning and verification

The predicted MTase genes were amplified from bacterial genomic DNAs using the gene-specific primers (Table S16), and cloned into plasmid pRRS as previously described [14]. The corresponding methylation sensitive restriction-site sequences (used to detect the activity of MTases) were included in the 3’-end oligonucleotides. The recombinant plasmids were then transformed into bacterial host ER2796 (not containing known MTase genes), followed by bacterial culture overnight. The plasmid DNAs were prepared using QIAprep Spin Miniprep Kit (catalog No. 27104, QIAGEN, Beijing, China). The appropriate restriction enzymes were then used to determine the presence or absence of methylated motifs in recombinant plasmid. The digestion reaction were conducted for 4 h at 37 °C and run on 1% agarose gels. The methylation motifs were also double detected by sequencing the recombinant plasmid by SMRT/bisulfite sequencing.

### Density distributions of GA<u>T</u>C and CCW<u>G</u>G motifs on the *K. pneumoniae* and simulated genomes

The simulated genome was generated based on the same length and GC content as those in the *K. pneumoniae* genomes. We then calculated the number of motifs in 1kb consecutive segments, so as to determine the density of GA<u>T</u>C and CCW<u>G</u>G motifs across the *K. pneumoniae* and simulated genomes.

We defined high/low-density regions by calculating the motif number in each 2kb-size non-overlap sliding window on the genome. Through normal distribution analysis using the pnorm function in R, the top and tail 5% of regions were defined as high-density and low-density regions, respectively.

### Ka/Ks analysis of the GA<u>T</u>C and CCW<u>G</u>G “motif sequences” and corresponding “scramble sequences”

We firstly extracted the minimum DNA sequences (2~3 codons) containing the motifs from the open reading frames of genes, in order to obtain “motif sequences”. The corresponding “scramble sequences” were obtained by random shuffling (eliminating the “motif sequences”). The top-10 frequent “scramble sequences” were used in subsequent analysis. We then identified the reference sequences of “motif sequences”/ “scramble sequences” on reference genome (HS11286) by multiple sequence alignment (mafft) [51]. The “motif sequences”, “scramble sequences” and corresponding reference sequences of each strain were respectively concatenated, and their Ka/ks ratios were calculated by paraAT [52].

### Sequence conservation analysis of the GA<u>T</u>C and CCW<u>G</u>G motifs

We extracted the methylated motifs and their flanking sequences (20 nt) in IGRs from one genome, and then obtained the corresponding sequences of 13 other genomes by multiple sequence alignment. Using the above 14 sequences, the conservation scores were calculated by phylop [53]. The conservation scores of the hemi-/un-methylated motif with 20-nt flanking sequences in IGRs were calculated by the same method.

### Calculation of the genomic replication time (t_R_) through simulating genomic coverage plots

The genomic coverage plots reflect the accumulated copy numbers across genomes. Since the replication forks always proceed from origin of replication (*oriC*) to doubling point (*s_DP_*), only a few ones can get close to the doubling point in the exponential phase. As a result, the sequencing coverage in *oriC* region is much higher than that in doubling point. In each cell, the copy number of each site (*s*) depends on its relative position to the first replication fork (*s_1_*) and the minimal time of successive initiations (re-initiation). The re-initiation time is consistent with the doubling time (*t_D_*) in the exponential phase, reflecting the growth rate of *K. pneumoniae* strains.

After initiation, the replication forks should go forward bi-directly at similar speeds in the exponential phase. Therefore, the genome coverage plots are symmetrical near the doubling point (*s_DP_*). We divided the coverage plots into the left (*s* ≤ *s_DP_*) and right (*s ≥ s_DP_*) parts *s ≤ s_DP_s ≥ s_DP_* by doubling point, and simulated the curves independently. We then obtained the relative position (*x* ∈ (0,1]) of each genomic site shown as equation (1). *S_G_* represents the half length of genome.

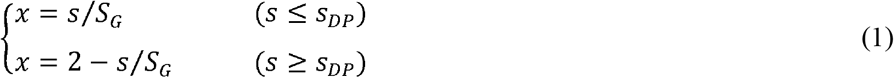

We subsequently calculated the relative distance of the first replication fork when another one at *oriC* is re-initiated (Δ*x_R_*) as equation (2). *t_R_* represents the genomic replication time.

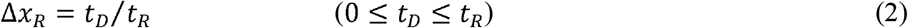

By constructing the replication frequency matrix of genomic sites (*x*) with different distances from the first replication fork (*x_1_*, the copy number of the genomic site (*f*_(*x,x*_1_)_ can be determined as equation (3). When the genomic site is in front of the first replication fork (*x* ≥ *x*_1_) its copy number is one; when the site is between the first and second replication forks (*x*_1_ – Δ*x_R_* ≤ *x* ≤ *x*_1_) its copy number is two. The rest was deduced by analogy.

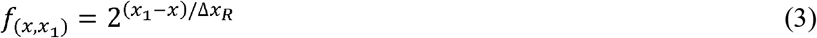

Next, we used the B distribution to assess cell population density with different genomic positions of the first replication forks (*x*_1_ ∈ [0,1]). Since *x* is not continuous data (with the step size of 1/*S_G_*), the cell population density of the first replication forks (*P*_(*x*_1_)_) at each genomic site was determined by the difference of accumulated densities (*I_x_*(*α, β*)) of the adjacent sites.

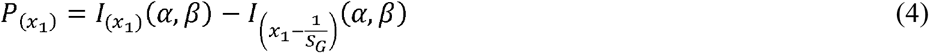

The genome coverage plots could be determined as the integral of the copy numbers of each genomic site in the cellular populations (*H*(*x*)) as shown in the equation (5). The integral was achieved based on *P*_(*x*_1_)_ and the relative distance between *x* and *x*_1_ (*x*_1_ – *x*). By substituting equation (2)–(4) into equation (5), we established the mathematical model to fit the genome coverage plots as equation (6).

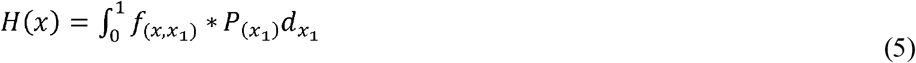

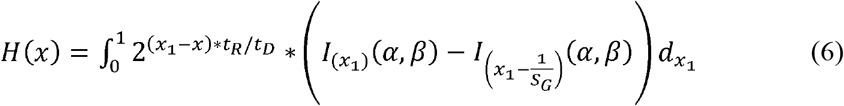

We then substituted equation (1) into equation (6), and repeatedly fit the genome coverage plots by selecting continuous parameters with step sizes of 0.1. We finally obtained the optimal solutions of Δ*x_R_* and B distribution parameters (*α, β*) through Goodness of Fit Test. Since *t_D_* could be calculated from growth curve of each *K. Pneumoniae* strain, we could obtain the replication time (*t_R_*) by the equation (2).

### Calculation of the re-modification time of motifs

In the exponential phase, methylation-read fraction (*M_(x)_*) of each motif was determined by its initial methylation-read fraction 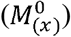, replication-induced passive de-methylation and methyltransferase-catalyzed re-methylation. Based on *M_(x)_* in the exponential phase and B distribution parameters (*α, β*), we calculated the proceeding distance of the corresponding replication fork when the hemi-methylated motif was re-methylated (Δ*x_M_*). As shown in the equation (7), Δ*x_M_* corresponds to the ratio of mean re-methylation time (*t_M_*) to the genomic replication time (*t_R_*).

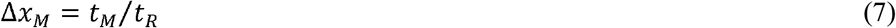

When the motif *x* is located at downstream of replication fork (*x_j_* ≤ *x*), its methylation-read fraction (*M_(x)_*) remain unchanged because of no replication-induced passive de-methylation. When the motif *x* is located at upstream of replication fork with a longer distance (*x_j_ ≥ x* + Δ*x_M_*), the methylation-read fraction is also unchanged because of the completion of re-methylation of motif *x*. Thus, we only considered the de-methylation effect of the replication forks in a certain range (*x* ≤ *x_j_* ≤ (*x* + Δ*x_M_*)) on *M_(x)_* in the mathematical derivation. *j* represents the serial number of the replication fork affecting *M_(x)_*.

If the hemi-methylated motifs can be fast re-modified before next replication fork (*t_M_* ≤ *t_D_*), we only need to evaluate the de-methylation effect of one replication for all replication forks in the range of *x* ≤ *x_j_* ≤ *x* + Δ*x_M_*. Here we include the first replication forks (*x*_1_) and other effective replication forks (*x_j_*) in the range. We then substituted the accumulated B distribution density of first replication forks (*x* + (*j* – 1)Δ*x_R_* ≤ *x*_1_ ≤ *x* + Δ*x_M_* + (*j* – 1)Δ*x_R_*) for that of the corresponding effective replication forks (*x* ≤ *x_j_* ≤ *x* + Δ*x_M_*). Due to semi-conservative replication, we further obtain Δ*x_M_* from equation (8). *N* represents the upper limit of *j*, which is determined by Δ*x_R_* and the relative genomic position (*x*) of methylated motifs.

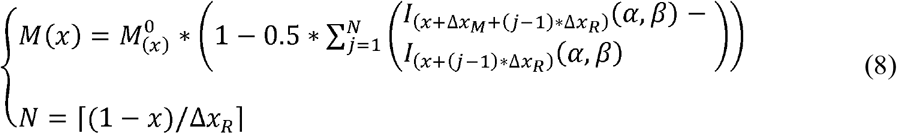

If the methylation rate is slow, the hemi-methylated motifs may not be remodified when the next replication fork crosses them (*t_M_* > *t_D_*). In this case, the effect of multiple passive de-methylation on *M_(x)_* should be considered. *n* represents upper limit of multiple passive de-methylation times (*i*) as equation (9), which is determined by the ratio of mean re-methylation time (*t_M_*) to re-initiation time(*t_D_*).

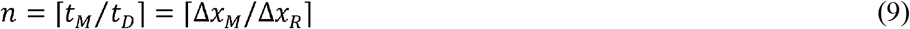

We then converted genomic position of effective replication forks (*x_j_*) into that of the corresponding first replication forks (*x*_1_), and substituted accumulated B distribution density of *x*_1_ (*x* + (*j* – 1) * Δ*x_R_* ≤ *x*_1_ ≤ *x* + *j* * Δ*x_R_*) for the *x_j_* population density (*x* ≤ *x_j_* ≤ *x* + Δ*x_M_*). Considering the differential influences on *M_(x)_* with the case of *j* < *n* and *n* ≤ *j* ≤ *N*, we calculated Δ*x_M_* as the equation (10) by comprehensive calculations.

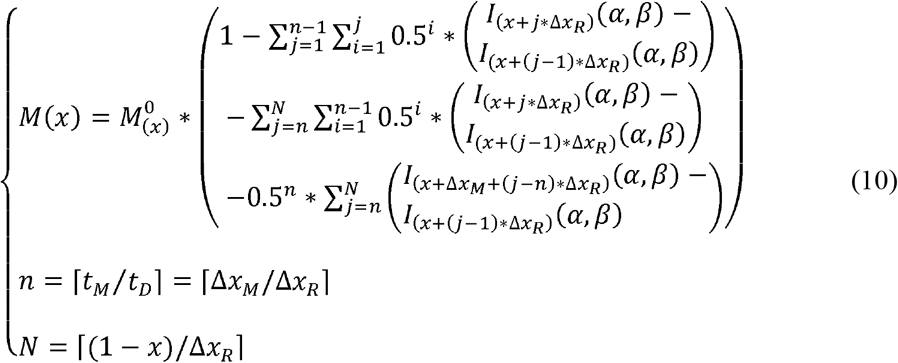

Based on the growth curve and genome coverage plots of the sequencing data, we deduce some parameters in different *K. pneumoniae* strains, such as *t_D_, t_R_* and *l_x_*(*α, β*). The mean re-methylation time (*t_M_*) of each motif could be further calculated by substituting the above parameters, the equation (2) and (7), as well as the methylation-read fraction in exponential phase (*M_(x)_*) and stationary phase 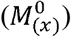 into the equation (10).

## Data availability

The genome data have been deposited in National Genomics Data Center (CRA003482) and NCBI (PRJNA477755) for strains of NTUH-K2044, 11492, 11420, 11454, 12208, 11311, 23, 11305, N201205880, 309074, 13190, 283747, 721005, 11021, and 11305.

## CRediT author statement

**Jing Fu**: Validation, Writing - Original Draft, Resources. **Ju Zhang**: Methodology, Writing - Original Draft, Resources. **Li Yang**: Software, Formal analysis, Writing - Original Draft. **Nan Ding**: Formal analysis, Writing - Original Draft, Visualization. **Liya Yue**: Validation, Writing - Original Draft, Visualization. **Xiangli Zhang**: Formal analysis, Visualization. **Dandan Lu**: Formal analysis, Visualization. **Xinmiao Jia**: Visualization. **Cuidan Li**: Visualization, Data Curation. **Chongye Guo**: Visualization. **Zhe Yin**: Resources. **Xiaoyuan Jiang**: Resources. **Yongliang Zhao**: Writing - Review & Editing. **Fei Chen**: Conceptualization, Writing - Review & Editing, Funding acquisition, Project administration, Supervision. **Dongsheng Zhou**: Conceptualization, Writing - Review & Editing, Project administration, Supervision.

## Acknowledgements

This work was supported by the National Key Program for Infectious Disease of China [2018ZX10302301-004-003] and the National Natural Science Foundation of China [31770870, 31801093].

## Supplementary material

**Figure S1 Phylogenetic analysis of 76 *K. pneumoniae* strains**

The strains shown in red letters represent the 14 *K. pneumoniae* strains we sequenced; the strains shown in black letters indicate the 62 *K. pneumoniae* strains downloaded from NCBI. The colored strips and stars represent various types of MLSTs and serotypes of *K. pneumoniae* strains.

**Figure S2 Genomic structural comparison of the 14 *K. pneumoniae* strains**

Genomes are arranged from the top to bottom in the order of: NTUH-K2044; 11492; 11420; 11454; 11311; 12208; 11311; 23; 11021; N201205880; 309074; 11305; 13190; 283747 and 721005. The same colors represent the homologous fragments as identified by the Mauve program.

**Figure S3 Preferred flanking sequences of the unmethylated, hemimethylated, and methylated MTC<u>G</u>AK motifs in NTUH-K2044 and 11492**

The sequence logos show the preferences of 10 nucleotides flanking the MT**C**<u>G</u>AK motifs in the two strains of NTUH-K2044 and 11492. The size of colored letters represents the probability of occurrence.

**Figure S4 Characterization of seven novel 6mA MTases specificities by using restriction digestion and SMRT sequencing**

**A**. Schematic diagram shows the whole MTase gene and its methylation motif sequence in the recombinant plasmids. We respectively cloned the MTases genes into pRRS plasmids. Type I MTases contain modification (M) and specificity (S) subunits, which are shown as blue and green bars. Type II MTases only include MTase subunits (M). The predicted methylation motif sequences are located at the downstream of the stop codon of the MTases. To identify the activity of MTases, we introduced some methylation-sensitive restriction enzymes, which recognition motifs share six bases (red dotted boxes) with the MTase motifs. If the A bases (red bold letter) are methylated by the MTases in the methyltransferase deficient *E. coli* strain ER2796, the restriction enzymes will fail to cut the corresponding motif sequences. **B.** Electrophoretogram identifying methylation activity of the MTases. As for KamB, the circular plasmid pRRS-KamB could not be cleave into linear plasmid by the methylation sensitive restriction enzymes, MfeI and BstBI (lane 4: circular plasmid control, lane 5: linear plasmid control, lane 6: plasmid pRRS-KamB cut by MfeI, lane 7: plasmid pRRS-KamB cut by BstBI), demonstrating that MTase KamB could successfully methylate TTCAN7TTC motif. As for KamC/D/E/F, since there were two restriction sites on the recombinant plasmids, the circular plasmid pRRS-KamC/D/E/F could be cleave into a linear fragment if the motifs were methylated (lane 8/11/14//17: circular plasmid control, lane 9/12/15/18: linear plasmid control, lane 10: plasmid pRRS-KamC cut by PciI, lane 13: plasmid pRRS-KamD cut by ScaI, lane 16: plasmid pRRS-KamE cut by BstBI, lane 19: plasmid pRRS-KamF cut by ScaI). Similarly, as for KamA/G, there were three restriction sites on the recombinant plasmids, the circular plasmid pRRS-KamA/G could be cleave into two linear fragment if the motifs were methylated (lane 1/20: circular plasmid control, lane 2/21: linear plasmid control, lane 3: plasmid pRRS-KamA cut by BfuAI, lane 22: plasmid pRRS-KamG cut by ScaI). For KamG, we also observed cleaved bands on the gel due to the uncomplete methylation (indicated as black arrows). **C.** IPD ratio plot shows the seven methylation motifs in the plasmids. SMRT sequencing was adopted to confirm the specificities of MTases. The purple and orange bars show the IPD ratios on plus and minus strands, respectively. The red rectangle indicates the motif sequence for each MTase. The average IPD values of the methylated bases on plus and minus strands are between four and eight.

**Figure S5 Characterization of 5mC MTases specificity using restriction digestion**

**A.** Schematic diagram shows the MTase gene and its methylation motif sequence in the recombinant plasmid. We cloned KcmA gene and predicted motif sequence (MT**C**<u>G</u>AK: at the downstream of the stop codon of the KcmA gene) into pRRS plasmid. The modification bases are marked as red bold letters. **B.** Electrophoretogram identifying the methylation activity of KcmA (a 5mC Mtase). The KcmA gene was cloned into the pRRS plasmid and expressed in E. coli ER2796. Recombinant plasmid DNA (pRRS-KcmA) was prepared and digested by the restriction enzyme BspDI. The products were resolved on an agarose gel for analysis. Lane M: Takara trans2K plusII ladder; Lane 1: recombinant plasmid pRRS-KcmA as positive control; Lane 2: linear recombinant plasmid pRRS-KcmA as negative control (digested by sbf I); Lane 3: pRRS-KcmA digested by BspD I to verify the methylation activity of KcmA; Lane 4: Takara trans2K plusII ladder.

**Figure S6 Seven predicted MTases without methylation activity**

**A.** Schematic diagram shows the seven predicted MTase genes and corrsponding methylation motif in the recombinant plasmids. **B.** Electrophoretogram identifying methylation activity of the MTases. lane 1/5/8/13/18/21/25/29: circular plasmid control; lane 2/6/9/14/19/22/30: linear plasmid control; lane 3: plasmid pRRS-M2B2 cut by MfeI; lane 4: plasmid pRRS-M2B2 cut by BstBI; lane 7: plasmid pRRS-M3B1 cut by BfuAI; lane 10: plasmid pRRS-M4B1 cut by BfuAI; lane 11: plasmid pRRS-M4B2 cut by MfeI; lane 12: plasmid pRRS-M4B2 cut by BstBI; lane 15: plasmid pRRS-M5B1 cut by BfuAI; lane 16: plasmid pRRS-M5B2 cut by MfeI; lane 17: plasmid pRRS-M5B2 cut by BstBI; lane 20: plasmid pRRS-M8D cut by BfuAI; lane 23: plasmid pRRS-M10E cut by BspEI; lane 24: plasmid pRRS-M10E cut by BstBI; lane 27: plasmid pRRS-M11E cut by BspEI; lane 28: plasmid pRRS-M11E cut by BstBI; lane 31: plasmid pRRS-M15G cut by ScaI.

**Figure S7 COG categories of coding genes with GA<u>T</u>C and CCW<u>G</u>G motifs in high-density and low-density regions**

X-axis shows the functional classes. Y-axis shows the number of genes in each functional class.

**Figure S8 Distribution of GA<u>T</u>C (upper panel) and CCW<u>G</u>G (low panel) motifs flanking the *oriC* region among the 14 *K. pneumonia* strains.**

The color intensity indicates the number of motif in 1kb window size.

**Figure S9 Density distribution of the GA<u>T</u>C/CCW<u>G</u>G motifs on the NTUH-K2044 genome and simulated genomes**

The orange histograms show the density distribution of G**A**<u>T</u>C/C**C**W<u>G</u>G motifs on the simulated genomes of strain NTUH-K2044. The green histograms show the density distribution of G**A**TC/C**C**WGG on the NTUH-K2044 genome.

**Figure S10 Density distribution of the GA<u>T</u>C/CCW<u>G</u>G motifs on the 13 *K. pneumoniae* genomes and random generated genomes**

The orange histograms show the density distribution of G**A**<u>T</u>C/C**C**W<u>G</u>G motifs on the random generated genomes. The green histograms show the density distribution of G**A**TC/C**C**WGG on the 13 *K. pneumoniae* genomes.

**Figure S11 Ka/Ks ratios for GA<u>T</u>C/CCW<u>G</u>G “motif sequences” and “scramble sequences” of the 14 *K. pneumoniae* genomes**

Box plot showing the Ka/Ks ratios of G**A**<u>T</u>C/C**C**W<u>G</u>G “motif sequences” of the *K. pneumoniae* genomes. Red boxes indicate the Ka/Ks ratios of G**A**<u>T</u>C/C**C**W<u>G</u>G “motif sequences”; blue boxes indicate the corresponding “scramble sequences” as the controls.

**Figure S12 Boxplot of conservations scores for methylated and hemi/unmethylated GA<u>T</u>C/CCW<u>G</u>G motifs**

Box plot shows the conservation values of G**A**<u>T</u>C/C**C**W<u>G</u>G motifs of the *K. pneumoniae* genomes. Red boxes indicate the conservation values of methylated G**A**<u>T</u>C/C**C**W<u>G</u>G motifs and their flanking regions (20 nt); blue boxes indicate the hemi/un-methylated motifs and their flanking regions (20 nt) as the controls.

**Figure S13 Genome-wide sequencing coverage and methylation level of GA<u>T</u>C and CCW<u>G</u>G motifs during cell cycles of NTUH-K2044 and 11492**

**A.** Genome-wide sequencing coverage versus genome position at three stages (1, 4 and 24 h) in the cell cycle of two *K. pneumonia* strains. The replication bidirectionally begins from the origin (O) and completes at terminus (T) (i. e., doubling point: in the middle of genome). The bold lines approximate the average coverage across the genomes. Ratios of the average coverage at *oriC* to that at doubling point are labeled in the figure. **B.** Genome-wide methylation level versus genome position for the two motifs (G**A**<u>T</u>C, C**C**W<u>G</u>G, MT**C**<u>G</u>AK and GR**A**CRAC) at the three growth stages in the cell cycle of two *K. pneumonia* strains. The bold lines approximate the average methylation levels across the genomes (5kb window size).

**Figure S14 Box plot showing the re-methylation time of motifs in strain 11492 and NTUH-K2044**

Box plots separately represent re-methylation time of motifs in strain 11492 (red) and NTUH-K2044 (blue). The black diamond mark represents the mean, and the transverse line represents the median.

**Figure S15 Preferred sequences flanking the GA<u>T</u>C and CCW<u>G</u>G motifs with fast and slow methylation rates in NTUH-K2044 and 11492.**

**A.** Preferred sequences flanking G**A**<u>T</u>C motifs with fast and slow methylation rates. The fast methylation rate means that the FRAC values for the motif are more than 0.95; the slow methylation rate means that the FRAC values for the motif are less than 0.9. Left panel: x-axis shows the G**A**<u>T</u>C motif flanked by five nucleotides; y-axis indicates the ratios of A/T/C/G. Right panel: the sequence logos show preferences of five nucleotides flanking the G**A**<u>T</u>C motifs at stationary phase in strain NTUH-K2044 and 11492. **B.** Preference of flanking nucleotides of C**C**W<u>G</u>G motif. Quick methylation mode was defined as the FRAC values for 1h, 4h, and 24h samples were all above 0.85, and the slow methylation mode was defined as the FRAC values for 1h, 4h, and 24h samples were all below 0.55. X-axis indicated the positions of nucleotides, and Y-axis indicated the ratios of A/T/C/G.

**Figure S16 Re-methylation time of GATC motifs in the *oriC* region (381 nt) of NTUH-K2044 and 11492**

The DnaA boxes (blue), AT-rich region (yellow) and G**A**<u>T</u>C motifs (red) are labeled on the sequence. The re-methylation times of G**A**<u>T</u>C motifs in 11492 and NTUH-K2044 stains are marked in purple and green.

**Figure S17 Distribution of the promoters in the upstream regions of 14 *K. pneumoniae* genomes**

X-axis shows the distance from the start codon; Y-axis shows the number of promoters locating corresponding positions of the region.

**Figure S18 COG distributions of genes with hemi/un-methylated GA<u>T</u>C/CCW<u>G</u>G motifs in intergenic regions**

Box plots indicated the COG categories of coding genes with methylated and hemi/un-methylated G**A**<u>T</u>C/C**C**W<u>G</u>G motifs in intergenic regions. X-axis shows the functional classes. Y-axis shows the ratio of genes in each functional class.

**Figure S19 COG distributions of genes with upstream hemi/un-methylated sites of novel motifs in intergenic regions**

X-axis shows the functional classes. Y-axis shows the ratio of genes in each functional class. [C] Energy production and conversion, [D] Cell cycle control, cell division, chromosome partitioning, [E] Amino acid transport and metabolism, [F] Nucleotide transport and metabolism, [G] Carbohydrate transport and metabolism, [H] Coenzyme transport and metabolism, [I] Lipid transport and metabolism, [J] Translation, ribosomal structure and biogenesis, [K] Transcription, [L] Replication, recombination and repair, [M] Cell wall/membrane/envelope biogenesis, [N] Cell motility, [O] Post-translational modification, protein turnover, and chaperones, [P] Inorganic ion transport and metabolism, [Q] Secondary metabolites biosynthesis, transport, and catabolism, [R] General function prediction only, [S] Function unknown, [T] Signal transduction mechanisms, [U] Intracellular trafficking, secretion, and vesicular transport, [V] Defense mechanisms.

**Figure S20 Accuracy evaluation of the mathematical model**

**A.** Comparison between the fitting and real methylated-read ratios (FRACmodel and FRACPacBio) of GATC motif (5kb window). **B.** Comparison between the fitting and real methylated-read ratios (FRAC_model_ and FRAC_PacBio_) of CCWGG motif (5kb window). **C.** Boxplot showing the percentage errors of GATC and CCWGG motifs.

**Table S1 Clinical information of 14 *K. pneumoniae* isolates**

**Table S2 Sequencing data of the 14 *K. pneumoniae* strains using SMRT technology**

**Table S3 Sequencing data of the 14 *K. pneumoniae* strains using Illumina Hiseq platform**

**Table S4 Genomic information of the online 62 *K. pneumoniae* strains used in the phylogenetic tree**

**Table S5 Bisulfite sequencing data of the 14 *K. pneumoniae* strains**

**Table S6 22 predicted MTase genes and the corresponding 15 methylation motifs**

**Table S7 Detailed information of the motifs and corresponding DNA MTases among the 14 *K. pneumoniae* strains**

**Table S8 Fishers’ exact test of the unmethylated and methylated MTC<u>G</u>AK sites**

**Table S9 Distribution of GATC motifs in GRs and IGRs among 14 *K. pneumoniae* strains**

**Table S10 Distribution of CCW<u>G</u>G motifs in GRs and IGRs among 14 *K. pneumoniae* strains**

**Table S11 Summary of genes with upstream hemi/un-methylated GA<u>T</u>C sites shared in the 14 *K. pneumoniae* strains**

**Table S12 Summary of genes with upstream hemi/un-methylated GA<u>T</u>C motif clusters in the 14 *K. pneumoniae* strains**

**Table S13 SMRT sequencing data of the samples at three growth time points (1, 4 and 24 h) of 11492 and NTUH-K2044**

**Table S14 Bisulfite sequencing data of the samples at three growth time points (1, 4 and 24 h) of 11492 and NTUH-K2044**

**Table S15 Slow re-methylated GA<u>T</u>C motifs in the 5’ USR of genes enriched in four COG categories**

**Table S16 The seven CCWGG sites changed in methylation state during the growth phase of NTUH-K2044**

**Table S17 Summary of the genes with upstream hemi/un-methylated MTCGAK sites shared in the NUTH-K2044 and 11492 strains**

**Table S18 Summary of the genes with upstream hemi/un-methylated CCAYN_7_TTYG sites shared in the 23 and 11311 strains**

**Table S19 Oligonucleotide primers used in this study**

## References

[1] Guo Y, Wang S, Zhan L, Jin Y, Duan J, Hao Z, et al. Microbiological and clinical characteristics of hypermucoviscous *Klebsiella pneumoniae* isolates associated with invasive infections in China. Front Cell Infect Microbiol 2017; 7: 24.

[2] Lam MMC, Wyres KL, Duchene S, Wick RR, Judd LM, Gan YH, et al. Population genomics of hypervirulent *Klebsiella pneumoniae* clonal-group 23 reveals early emergence and rapid global dissemination. Nat Commun 2018; 9: 2703.

[3] Yeh KM, Kurup A, Siu LK, Koh YL, Fung CP, Lin JC, et al. Capsular serotype K1 or K2, rather than magA and rmpA, is a major virulence determinant for *Klebsiella pneumoniae* liver abscess in Singapore and Taiwan. J Clin Microbiol 2007; 45: 466–71.

[4] Bialek-Davenet S, Criscuolo A, Ailloud F, Passet V, Jones L, Delannoy-Vieillard AS, et al. Genomic definition of hypervirulent and multidrug-resistant *Klebsiella pneumoniae* clonal groups. Emerg Infect Dis 2014; 20: 1812–20.

[5] Hammond AW, Gerard GF, Chatterjee DK. Cloning the KpnI restriction-modification system in *Escherichia coli*. Gene 1991; 97: 97–102.

[6] Valinluck B, Lee NS, Ryu J. A new restriction-modification system, KpnBI, recognized in *Klebsiella pneumoniae*. Gene 1995; 167: 59–62.

[7] Mehling JS, Lavender H, Clegg S. A Dam methylation mutant of *Klebsiella pneumoniae* is partially attenuated. FEMS Microbiol Lett 2007; 268: 187–93.

[8] Fang CT, Yi WC, Shun CT, Tsai SF. DNA adenine methylation modulates pathogenicity of *Klebsiella pneumoniae* genotype K1. J Microbiol Immunol Infect 2017; 50: 471–7.

[9] Fang G, Munera D, Friedman DI, Mandlik A, Chao MC, Banerjee O, et al. Genome-wide mapping of methylated adenine residues in pathogenic *Escherichia coli* using single-molecule real-time sequencing. Nat Biotechnol 2012; 30: 1232–9.

[10] Bottacini F, Morrissey R, Roberts RJ, James K, van Breen J, Egan M, et al. Comparative genome and methylome analysis reveals restriction/modification system diversity in the gut commensal *Bifidobacterium breve*. Nucleic Acids Res 2018; 46: 1860–77.

[11] Lluch-Senar M, Luong K, Llorens-Rico V, Delgado J, Fang G, Spittle K, et al. Comprehensive methylome characterization of *Mycoplasma genitalium* and *Mycoplasma pneumoniae* at single-base resolution. PLoS Genet 2013; 9: e1003191.

[12] Hargreaves KR, Thanki AM, Jose BR, Oggioni MR, Clokie MR. Use of single molecule sequencing for comparative genomics of an environmental and a clinical isolate of *Clostridium difficile* ribotype 078. BMC Genomics 2016; 17: 1020.

[13] Murray IA, Clark TA, Morgan RD, Boitano M, Anton BP, Luong K, et al. The methylomes of six bacteria. Nucleic Acids Res 2012; 40: 11450–62.

[14] Zhu L, Zhong J, Jia X, Liu G, Kang Y, Dong M, et al. Precision methylome characterization of *Mycobacterium tuberculosis complex* (MTBC) using PacBio single-molecule real-time (SMRT) technology. Nucleic Acids Res 2016; 44: 730–43.

[15] Cota I, Bunk B, Sproer C, Overmann J, Konig C, Casadesus J. OxyR-dependent formation of DNA methylation patterns in OpvABOFF and OpvABON cell lineages of *Salmonella enterica*. Nucleic Acids Res 2016; 44: 3595–609.

[16] Cohen NR, Ross CA, Jain S, Shapiro RS, Gutierrez A, Belenky P, et al. A role for the bacterial GATC methylome in antibiotic stress survival. Nat Genet 2016; 48: 581–6.

[17] Wolanski M, Donczew R, Zawilak-Pawlik A, Zakrzewska-Czerwinska J. oriC-encoded instructions for the initiation of bacterial chromosome replication. Front Microbiol 2014; 5: 735.

[18] Kozdon JB, Melfi MD, Luong K, Clark TA, Boitano M, Wang S, et al. Global methylation state at base-pair resolution of the Caulobacter genome throughout the cell cycle. Proc Natl Acad Sci U S A 2013; 110: E4658–67.

[19] Lobner-Olesen A, Hansen FG, Rasmussen KV, Martin B, Kuempel PL. The initiation cascade for chromosome replication in wild-type and Dam methyltransferase deficient *Escherichia coli* cells. EMBO J 1994; 13: 1856–62.

[20] Lahue RS, Su SS, Modrich P. Requirement for d(GATC) sequences in *Escherichia coli* mutHLS mismatch correction. Proc Natl Acad Sci U S A 1987; 84: 1482–6.

[21] Blow MJ, Clark TA, Daum CG, Deutschbauer AM, Fomenkov A, Fries R, et al. The Epigenomic Landscape of Prokaryotes. PLoS Genet 2016; 12: e1005854.

[22] Lee CR, Lee JH, Park KS, Kim YB, Jeong BC, Lee SH. Global dissemination of carbapenemase-producing *Klebsiella pneumoniae*: epidemiology, genetic context, treatment options, and detection methods. Front Microbiol 2016; 7: 895.

[23] Darling AE, Mau B, Perna NT. progressiveMauve: multiple genome alignment with gene gain, loss and rearrangement. PLoS One 2010; 5: e11147.

[24] Roberts RJ, Vincze T, Posfai J, Macelis D. REBASE--a database for DNA restriction and modification: enzymes, genes and genomes. Nucleic Acids Res 2015; 43: D298–9.

[25] Hurst LD. The Ka/Ks ratio: diagnosing the form of sequence evolution. Trends Genet 2002; 18: 486–487

[26] de Jong A, Pietersma H, Cordes M, Kuipers OP, Kok J. PePPER: a webserver for prediction of prokaryote promoter elements and regulons. BMC Genomics 2012; 13: 299.

[27] Luo H, Gao F, Lin Y. Evolutionary conservation analysis between the essential and nonessential genes in bacterial genomes. Sci Rep 2015; 5: 13210.

[28] Waldron DE, Owen P, Dorman CJ. Competitive interaction of the OxyR DNA-binding protein and the Dam methylase at the antigen 43 gene regulatory region in *Escherichia coli*. Molecular Microbiology 2002; 44: 509–20.

[29] Campbell JL, Kleckner N. *E. coli* oriC and the dnaA gene promoter are sequestered from *dam* methyltransferase following the passage of the chromosomal replication fork. Cell 1990; 62: 967–79.

[30] Chang SW, Chao WR, Ruan A, Wang PH, Lin JC, Han CP. A promising hypothesis of c-KIT methylation/ expression paradox in c-KIT (+) squamous cell carcinoma of uterine cervix-CTCF transcriptional repressor regulates c-KIT proto-oncogene expression. Diagn Pathol 2015; 10: 207.

[31] Nievera C, Torgue JJ, Grimwade JE, Leonard AC. SeqA blocking of DnaA-oriC interactions ensures staged assembly of the *E. coli* pre-RC. Mol Cell 2006; 24: 581–92.

[32] Camacho EM, Casadesus J. Regulation of traJ transcription in the Salmonella virulence plasmid by strand-specific DNA adenine hemimethylation. Mol Microbiol 2005; 57: 1700–18.

[33] Lopez Sanchez MIG, Cipullo M, Gopalakrishna S, Khawaja A and Rorbach J. Methylation of ribosomal RNA: a mitochondrial perspective. Front. Genet 2020; 11: 761.

[34] Radman-Livaja M, Liu CL, Friedman N, Schreiber SL, Rando OJ. Replication and active demethylation represent partially overlapping mechanisms for erasure of H3K4me3 in budding yeast. PLOS Genetics 2010; 6: e1000837.

[35] Coffin SR, Reich NO. Modulation of Escherichia coli DNA methyltransferase activity by biologically derived GATC-flanking sequences. J Biol Chem 2008; 283: 20106–16.

[36] Pollak AJ, Chin AT, Brown FL, Reich NO. DNA looping provides for “intersegmental hopping” by proteins: a mechanism for long-range site localization. J Mol Biol 2014; 426: 3539–52.

[37] Barel I, Naughton B, Reich NO, Brown FLH. Specificity versus processivity in the sequential modification of DNA: a study of DNA adenine methyltransferase. J Phys Chem B 2018; 122: 1112–20.

[38] Zheng Y, Sweet SM, Popovic R, Martinez-Garcia E, Tipton JD, Thomas PM, et al. Total kinetic analysis reveals how combinatorial methylation patterns are established on lysines 27 and 36 of histone H3. Proc Natl Acad Sci U S A 2012; 109: 13549–54.

[39] Wodarz D, Boland CR, Goel A, Komarova NL. Methylation kinetics and CpG-island methylator phenotype status in colorectal cancer cell lines. Biol Direct 2013; 8: 14.

[40] Hojfeldt JW, Laugesen A, Willumsen BM, Damhofer H, Hedehus L, Tvardovskiy A, et al. Accurate H3K27 methylation can be established de novo by SUZ12-directed PRC2. Nat Struct Mol Biol 2018; 25: 225–32.

[41] Siu LK, Yeh KM, Lin JC, Fung CP, Chang FY. *Klebsiella pneumoniae* liver abscess: a new invasive syndrome. The Lancet Infectious Diseases 2012; 12: 881–7.

[42] English AC, Richards S, Han Y, Wang M, Vee V, Qu J, et al. Mind the gap: upgrading genomes with Pacific Biosciences RS long-read sequencing technology. PLoS One 2012; 7: e47768.

[43] Brettin T, Davis JJ, Disz T, Edwards RA, Gerdes S, Olsen GJ, et al. RASTtk: a modular and extensible implementation of the RAST algorithm for building custom annotation pipelines and annotating batches of genomes. Sci Rep 2015; 5: 8365.

[44] Loytynoja A, Goldman N. webPRANK: a phylogeny-aware multiple sequence aligner with interactive alignment browser. BMC Bioinformatics 2010; 11: 579.

[45] Page AJ, Cummins CA, Hunt M, Wong VK, Reuter S, Holden MT, et al. Roary: rapid large-scale prokaryote pan genome analysis. Bioinformatics 2015; 31: 3691–3.

[46] Price MN, Dehal PS, Arkin AP. FastTree 2-approximately maximum-likelihood trees for large alignments. PLoS One 2010; 5: e9490.

[47] He Z, Zhang H, Gao S, Lercher MJ, Chen WH, Hu S. Evolview v2: an online visualization and management tool for customized and annotated phylogenetic trees. Nucleic Acids Res 2016; 44: W236–41.

[48] Ma W, Noble WS, Bailey TL. Motif-based analysis of large nucleotide data sets using MEME-ChIP. Nat Protoc 2014; 9: 1428–50.

[49] Bolger AM, Lohse M, Usadel B. Trimmomatic: a flexible trimmer for Illumina sequence data. Bioinformatics 2014; 30: 2114–20.

[50] Krueger F, Andrews SR. Bismark: a flexible aligner and methylation caller for Bisulfite-Seq applications. Bioinformatics 2011; 27: 1571–2.

[51] Katoh K, Standley DM. MAFFT multiple sequence alignment software version 7: improvements in performance and usability. Mol Biol Evol 2013; 30: 772–80.

[52] Zhang Z, Xiao J, Wu J, Zhang H, Liu G, Wang X, et al. ParaAT: a parallel tool for constructing multiple protein-coding DNA alignments. Biochem Biophys Res Commun 2012; 419: 779–81.

[53] Cooper GM, Stone EA, Asimenos G, Program NCS, Green ED, Batzoglou S, et al. Distribution and intensity of constraint in mammalian genomic sequence. Genome Res 2005; 15: 901–13.

