## Supplemental data for "Precision Methylome and *in vivo* Methylation Kinetics Characterization of *Klebsiella Pneumoniae*"

**Running title**: *Fu J et al / Dynamic Methylome of Klebsiella Pneumoniae*

**Supplementary figures**

**Figure S1**

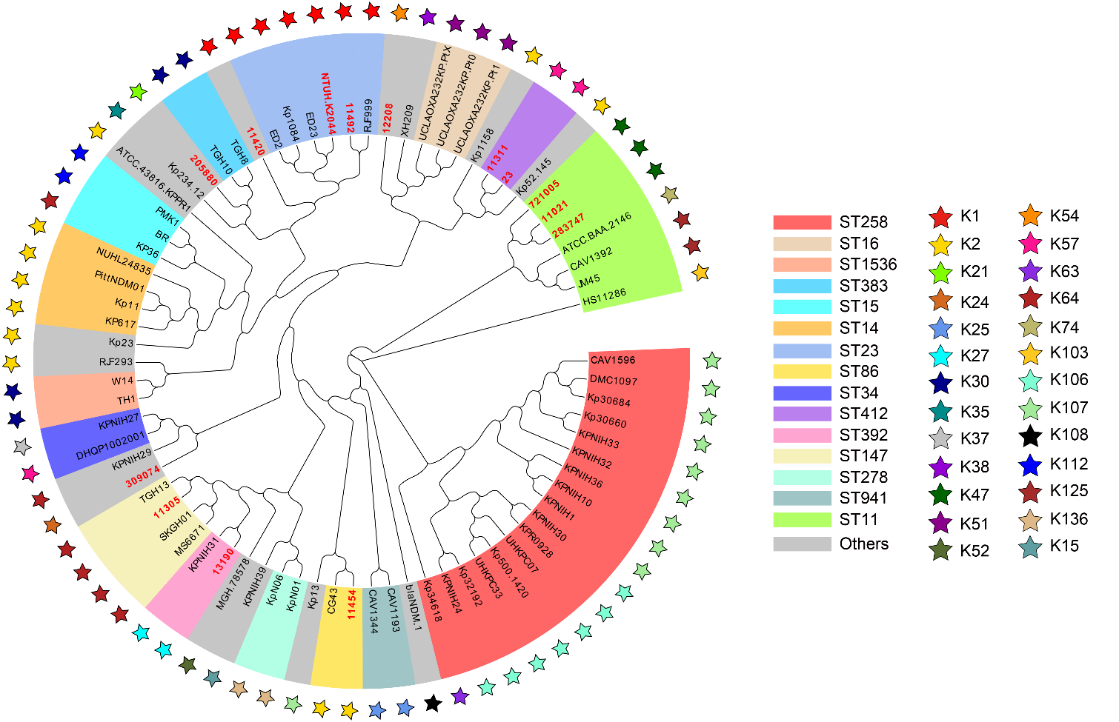

**Figure S1 Phylogenetic analysis of 76 *K. pneumoniae* strains (including our 14 and 62 online *K. pneumoniae* strains)**

The strains shown in red letters represent the 14 *K. pneumoniae* strains we sequenced; the strains shown in black letters indicate the 62 *K. pneumoniae* strains downloaded from NCBI. The colored strips and stars represent various types of MLSTs and serotypes of *K. pneumoniae* strains.

**Figure S2**

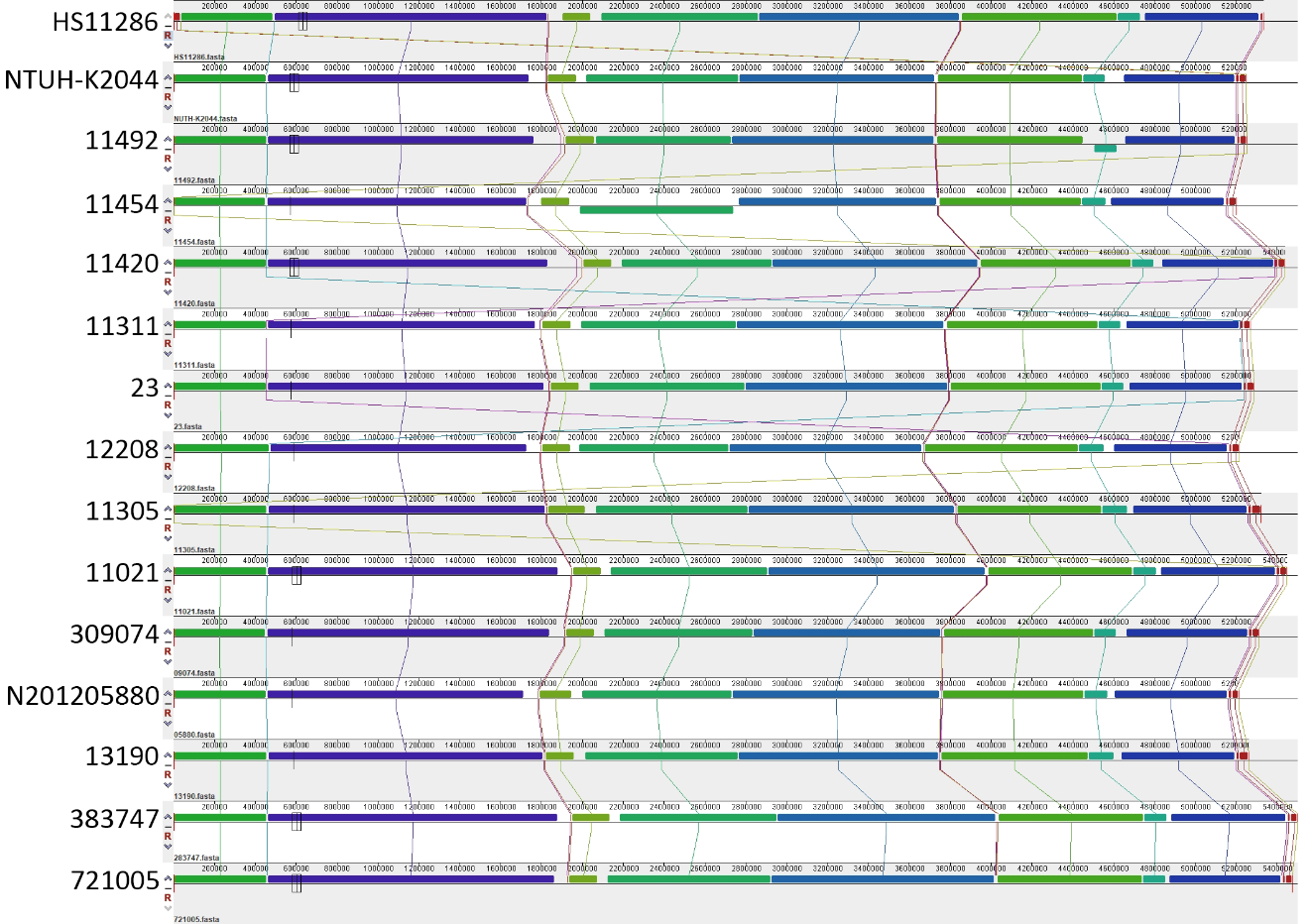

**Figure S2** **Genomic structural comparison of the 14 *K. pneumoniae* strains**

Genomes are arranged from the top to bottom in the order of: NTUH-K2044; 11492; 11420; 11454; 11311; 12208; 11311; 23; 11021; N201205880; 309074; 11305; 13190; 283747 and 721005. The same colors represent the homologous fragments as identified by the Mauve program.

**Figure S3**

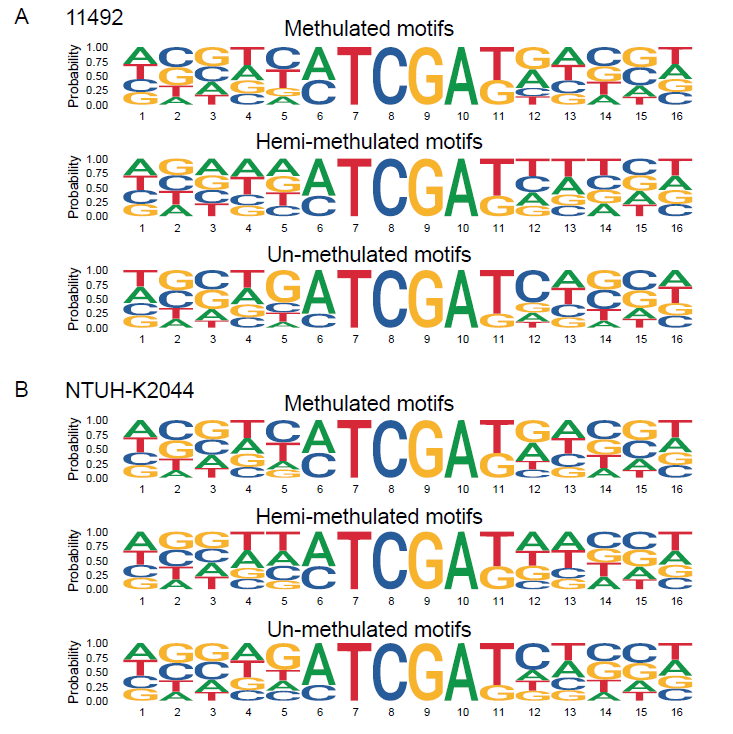

### Figure S3 Preferred flanking sequences of the methylated, hemi-methylated, and un-methylated MTCGAK motifs in *K. pneumoniae* strains 11492 and NTUH-K2044.

The sequence logos show the preferences of 10 nucleotides flanking the MT**C**GAK motifs at stationary phase in strain 11492 (A) and NTUH-K2044 (B).

**Figure S4**

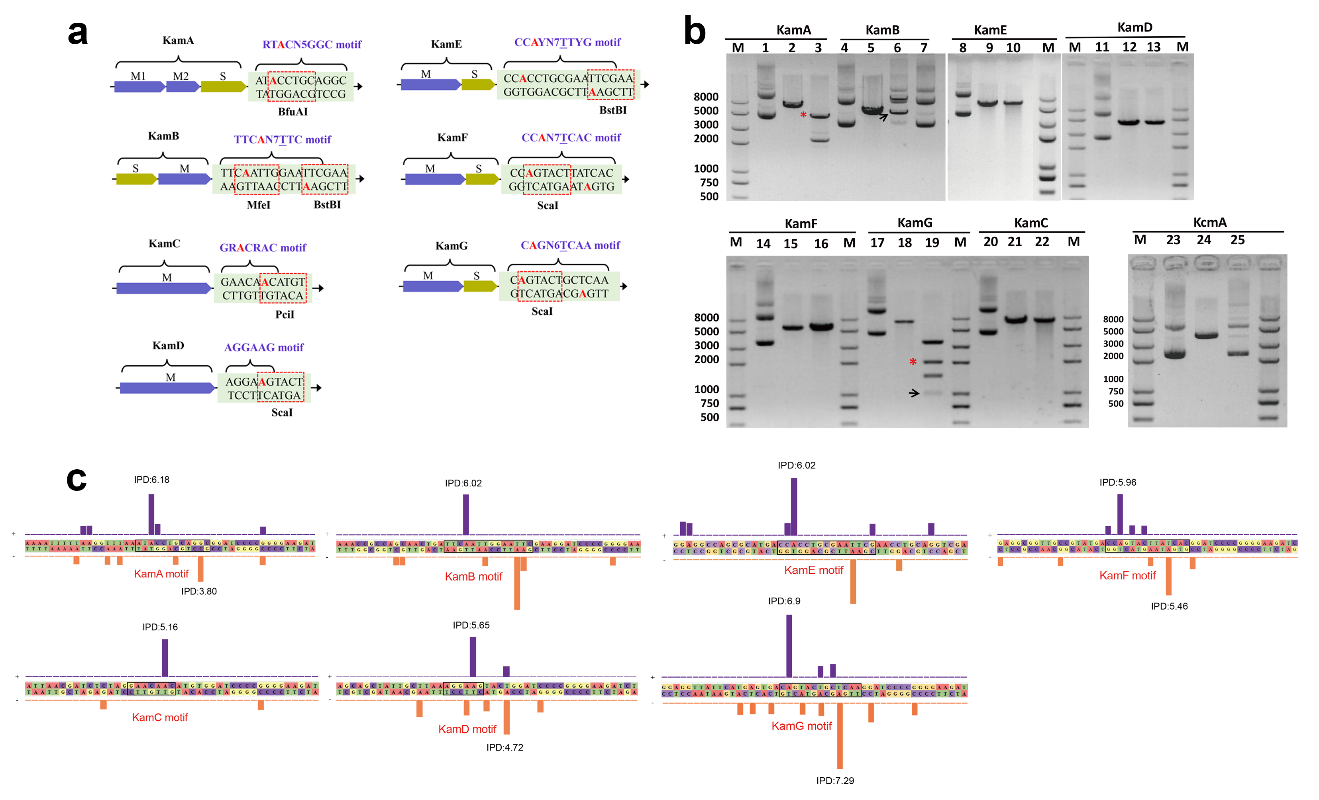

**Figure S4** **Characterization of seven novel 6mA MTases specificities by using restriction digestion and SMRT sequencing**

**(A)** Schematic diagram shows the whole MTase gene and its methylation motif sequence in the recombinant plasmids. We respectively cloned the MTases genes into pRRS plasmids. Type I MTases contain modification (M) and specificity (S) subunits, which are shown as blue and green bars. Type II MTases only include MTase subunits (M). The predicted methylation motif sequences are located at the downstream of the stop codon of the MTases. To identify the activity of MTases, we introduced some methylation-sensitive restriction enzymes, which recognition motifs share six bases (red dotted boxes) with the MTase motifs. If the A bases (red bold letter) are methylated by the MTases in the methyltransferase deficient *E. coli* strain ER2796, the restriction enzymes will fail to cut the corresponding motif sequences. **(B)** Electrophoretogram identifying methylation activity of the MTases. As for KamB, the circular plasmid pRRS-KamB could not be cleave into linear plasmid by the methylation sensitive restriction enzymes, MfeI and BstBI (lane 4: circular plasmid control, lane 5: linear plasmid control, lane 6: plasmid pRRS-KamB cut by MfeI, lane 7: plasmid pRRS-KamB cut by BstBI), demonstrating that MTase KamB could successfully methylate TTCAN_7_TTC motif. As for KamC/D/E/F, since there were two restriction sites on the recombinant plasmids, the circular plasmid pRRS-KamC/D/E/F could be cleave into a linear fragment if the motifs were methylated (lane 8/11/14//17: circular plasmid control, lane 9/12/15/18: linear plasmid control, lane 10: plasmid pRRS-KamC cut by PciI, lane 13: plasmid pRRS-KamD cut by ScaI, lane 16: plasmid pRRS-KamE cut by BstBI, lane 19: plasmid pRRS-KamF cut by ScaI). Similarly, as for KamA/G, there were three restriction sites on the recombinant plasmids, the circular plasmid pRRS- KamA/G could be cleave into two linear fragment if the motifs were methylated (lane 1/20: circular plasmid control, lane 2/21: linear plasmid control, lane 3: plasmid pRRS-KamA cut by BfuAI, lane 22: plasmid pRRS-KamG cut by ScaI). For KamG, we also observed cleaved bands on the gel due to the uncomplete methylation (indicated as black arrows). **(c)** IPD ratio plot shows the seven methylation motifs in the plasmids. SMRT sequencing was adopted to confirm the specificities of MTases. The purple and orange bars show the IPD ratios on plus and minus strands, respectively. The red rectangle indicates the motif sequence for each MTase. The average IPD values of the methylated bases on plus and minus strands are between four and eight.

**Figure S5**

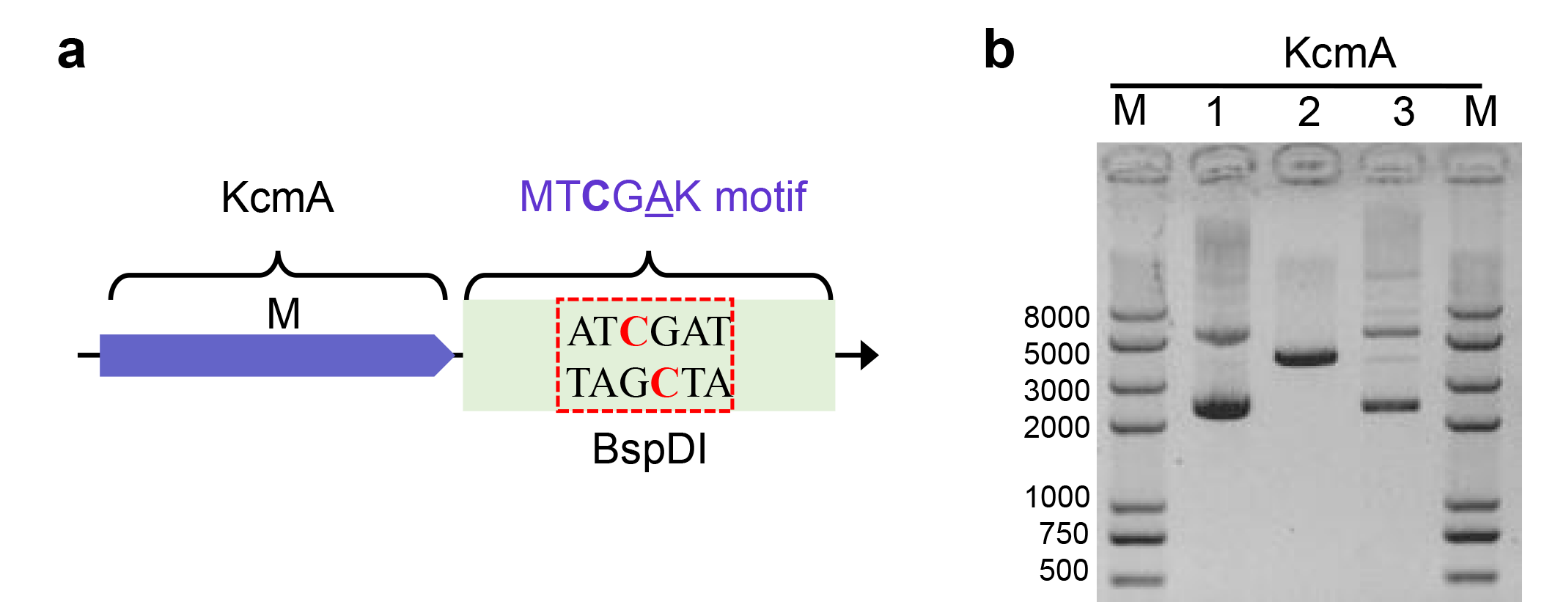

**Figure S5 Characterization of one 5mC MTases specificity using restriction digestion**

**(A)** Schematic diagram shows the MTase gene and its methylation motif sequence in the recombinant plasmid. We cloned KcmA gene and predicted motif sequence (MT**C**GAK: at the downstream of the stop codon of the KcmA gene) into pRRS plasmid. The modification bases are marked as red bold letters. **(B)** Electrophoretogram identifying the methylation activity of KcmA (a 5mC Mtase). The KcmA gene was cloned into the pRRS plasmid and expressed in *E. coli* ER2796. Recombinant plasmid DNA (pRRS-KcmA) was prepared and digested by the restriction enzyme BspDI. The products were resolved on an agarose gel for analysis.

Lane M: Takara trans2K plusII ladder; Lane 1: recombinant plasmid pRRS-KcmA as positive control; Lane 2: linear recombinant plasmid pRRS-KcmA as negative control (digested by sbf I); Lane 3: pRRS-KcmA digested by BspDI to verify the methylation activity of KcmA; Lane 4: Takara trans2K plusII ladder.

**Figure S6**

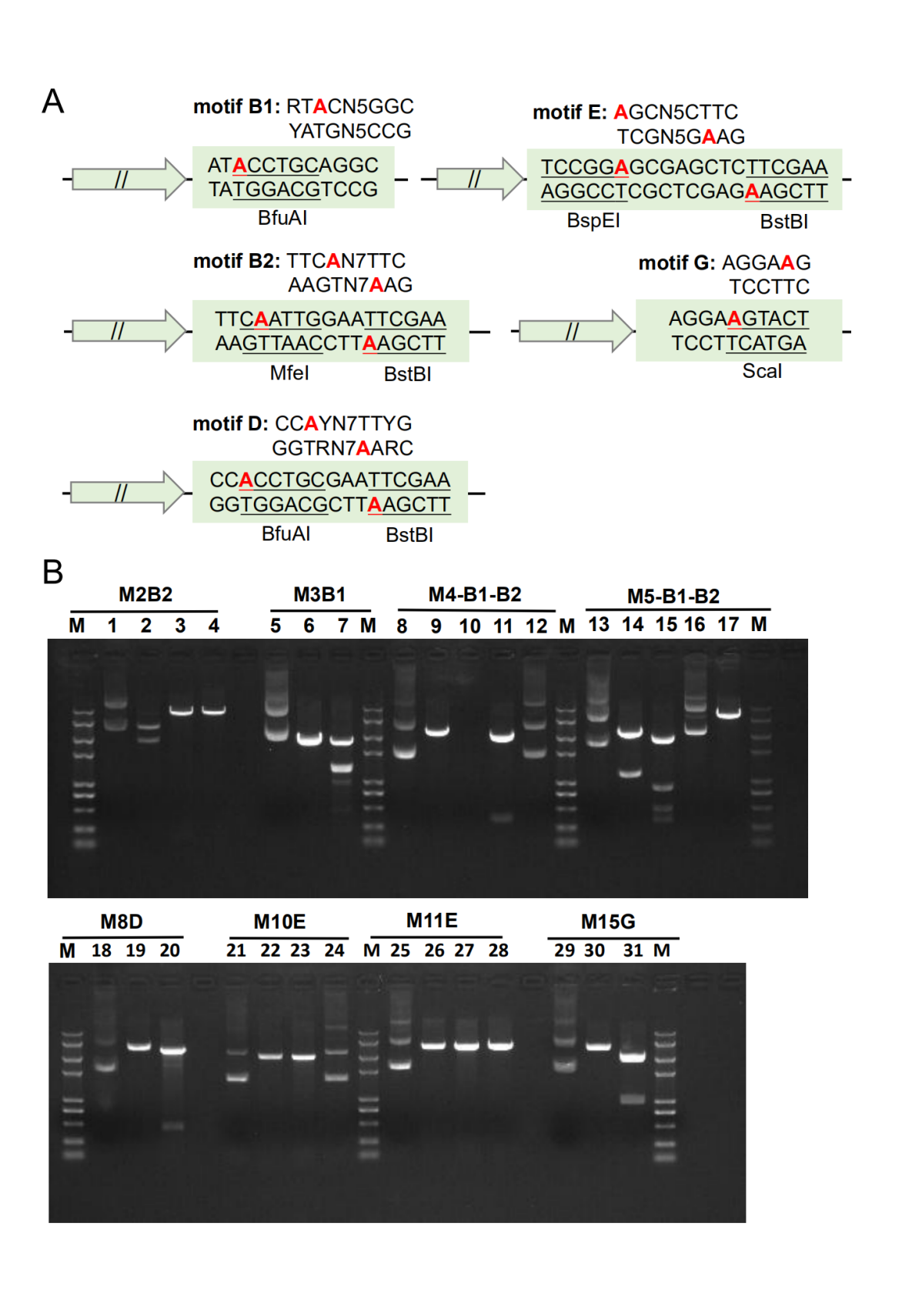

**Figure S6 The seven predicted MTases did not methylate the corresponding motifs**

In this study, we predicted 22 MTases that might be responsible for the 15 methylated motifs from *K. pneumoniae* genomes in the REBASE database. Crossover validations identified the corresponding 15 MTases that can specifically recognize and methylate the respective 15 motifs The other seven predicted MTases did not methylate the corresponding motifs.

**(A)** Schematic diagram shows the seven predicted MTase genes and corrsponding methylation motif in the recombinant plasmids. **(B)** Electrophoretogram identifying methylation activity of the MTases. lane 1/5/8/13/18/21/25/29: circular plasmid control; lane 2/6/9/14/19/22/30: linear plasmid control; lane 3: plasmid pRRS-M2B2 cut by MfeI; lane 4: plasmid pRRS-M2B2 cut by BstBI; lane 7: plasmid pRRS-M3B1 cut by BfuAI; lane 10: plasmid pRRS-M4B1 cut by BfuAI; lane 11: plasmid pRRS-M4B2 cut by MfeI; lane 12: plasmid pRRS-M4B2 cut by BstBI; lane 15: plasmid pRRS-M5B1 cut by BfuAI; lane 16: plasmid pRRS-M5B2 cut by MfeI; lane 17: plasmid pRRS-M5B2 cut by BstBI; lane 20: plasmid pRRS-M8D cut by BfuAI; lane 23: plasmid pRRS-M10E cut by BspEI; lane 24: plasmid pRRS-M10E cut by BstBI; lane 27: plasmid pRRS-M11E cut by BspEI; lane 28: plasmid pRRS-M11E cut by BstBI; lane 31: plasmid pRRS-M15G cut by ScaI.

**Figure S7**

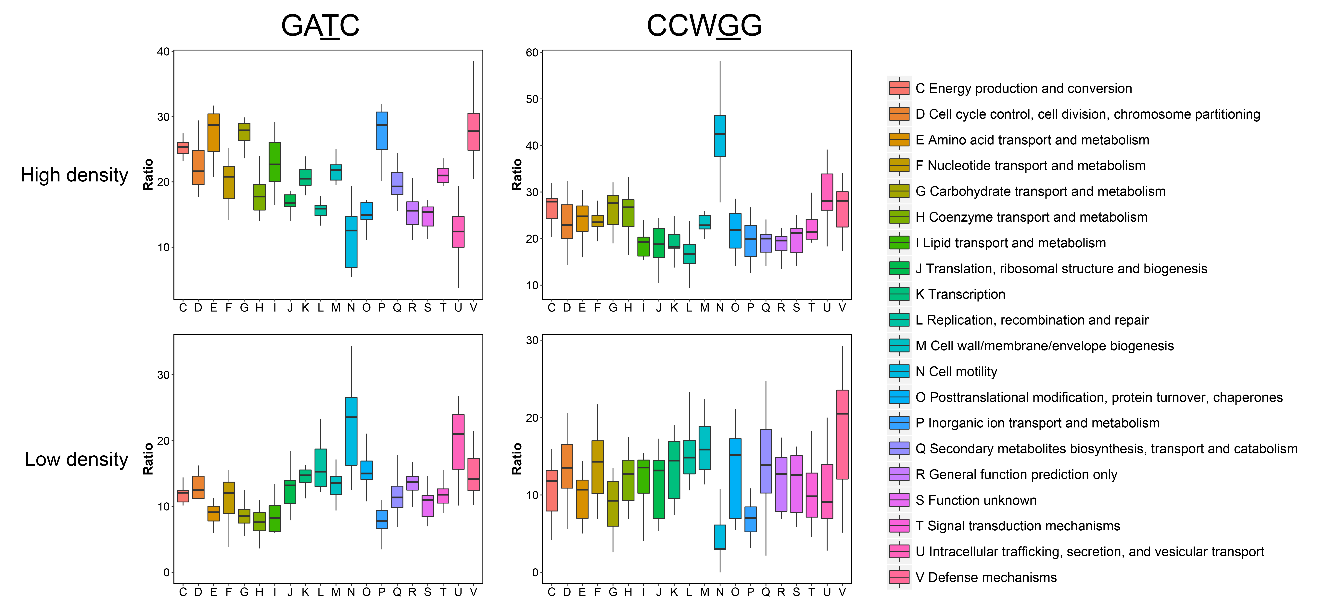

**Figure S7** **COG distributions of coding genes with GATC and CCWGG motifs**

Box plots indicated the COG categories of coding genes with motifs. Functional classes of genes containing the motif G**A**TC and C**C**WGG motifs across COG categories. X-axis shows the functional classes. Y-axis shows the number of genes in each functional class.

**Figure S8**

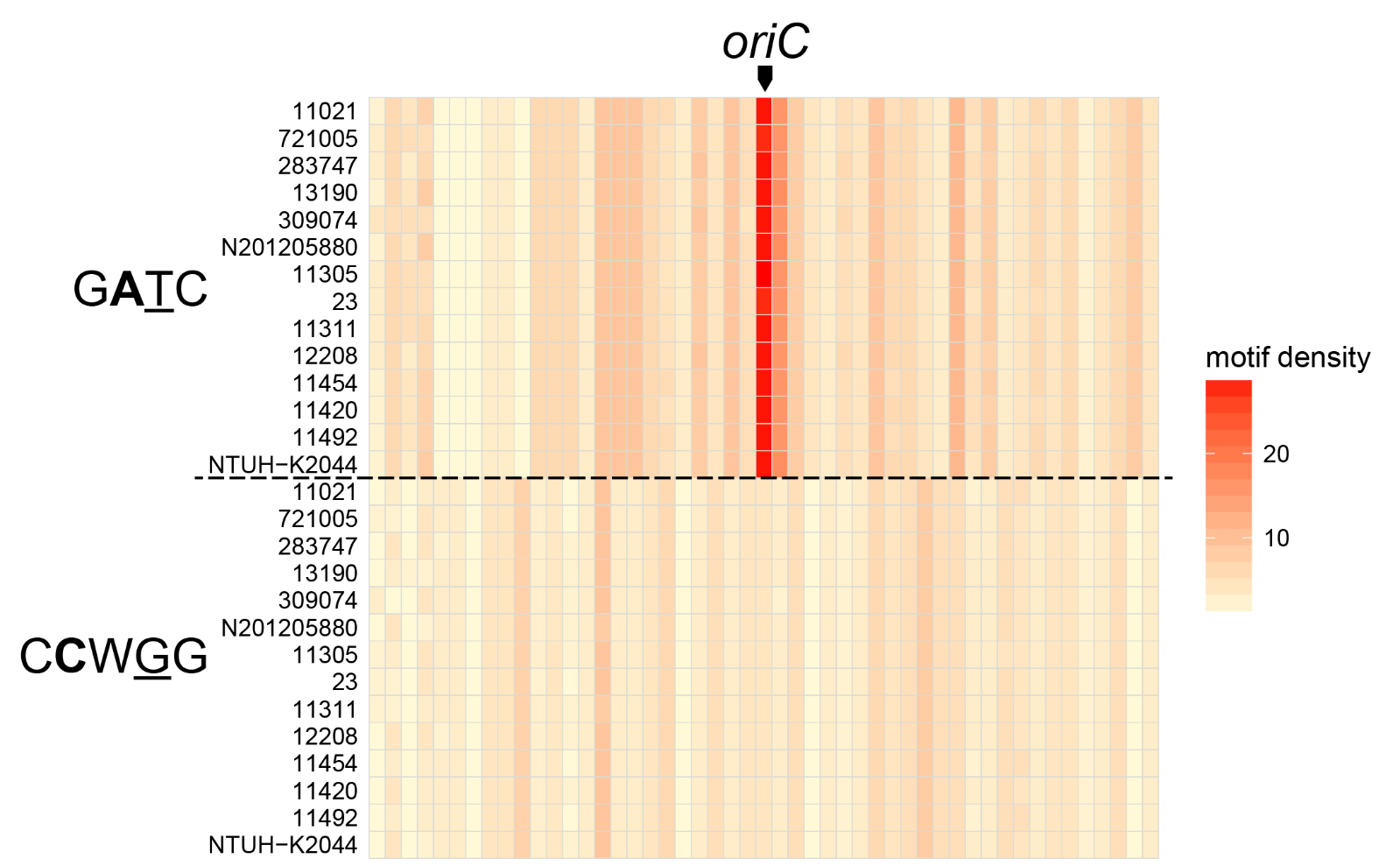

**Figure S8 Heatmap showing the distribution of GATC (upper panel) and CCWGG (low panel) motifs flanking the oriC region among the 14 *K. pneumonia* strains**

The color intensity indicates the number of motif in 1kb window size.

**Figure S9**

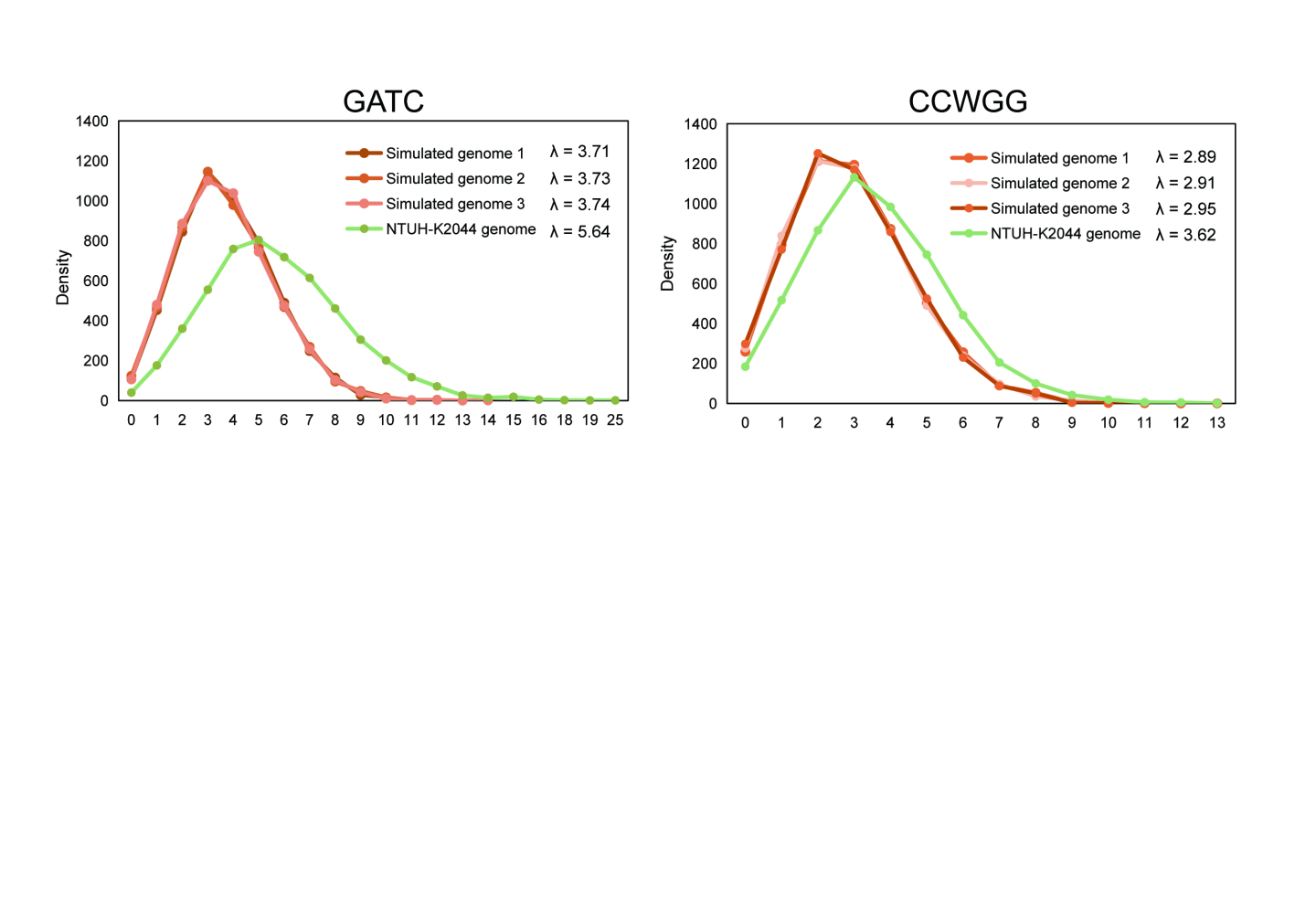

**Figure S9 Density distribution of the GATC/CCWGG motifs on the NTUH-K2044 genome and simulated genomes**

**Figure S10**
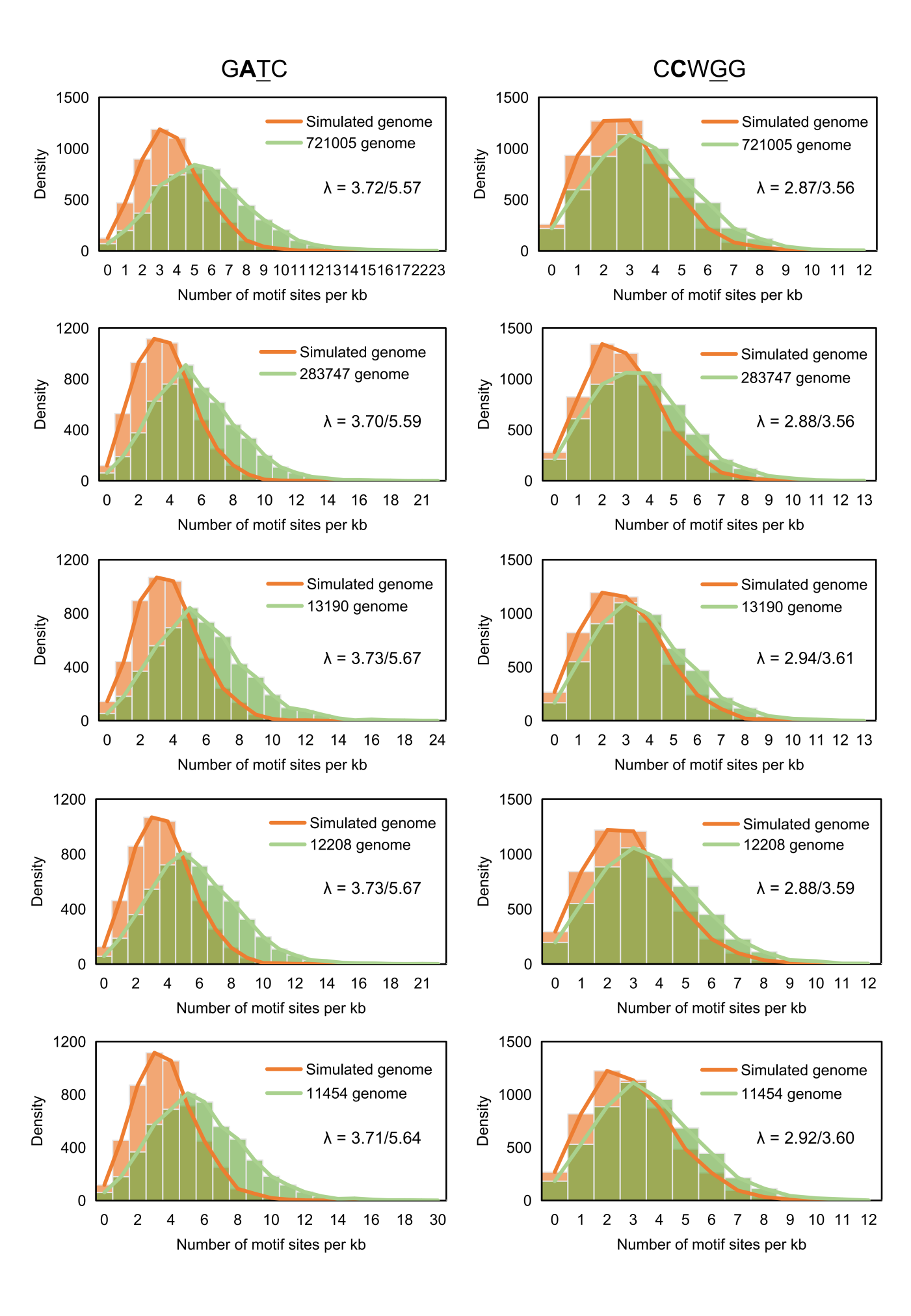

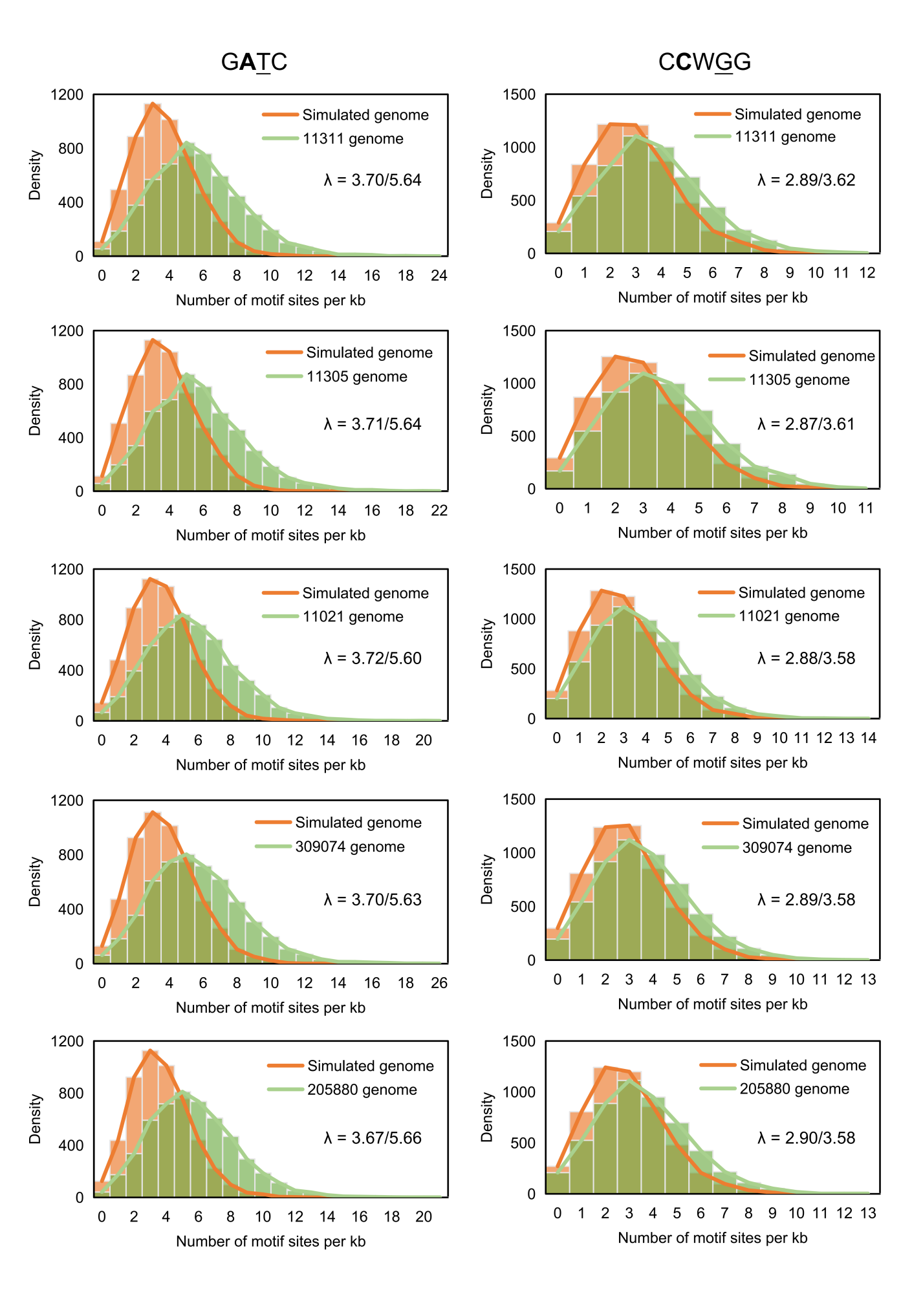

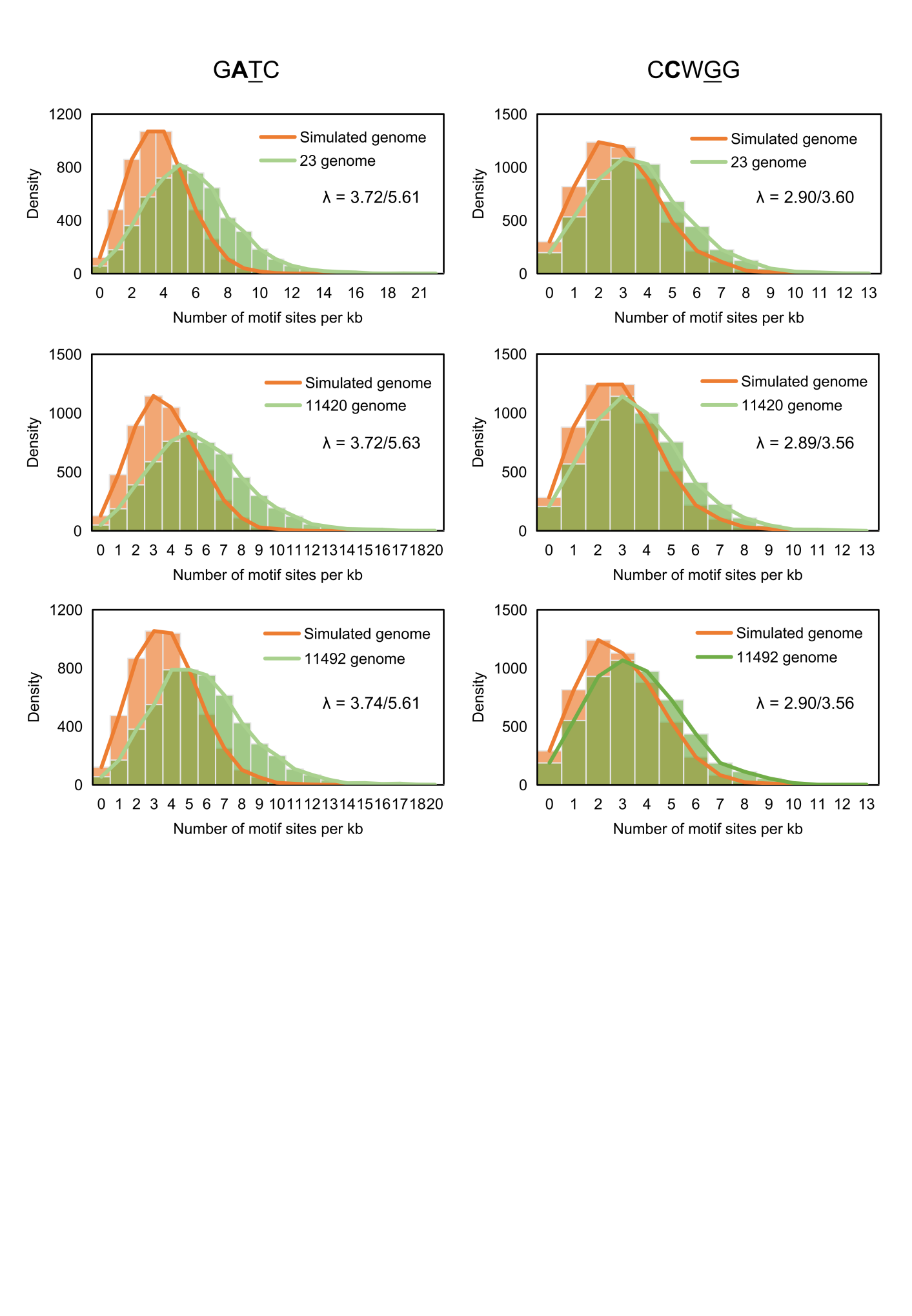

**Figure S10 Density distribution of the GATC/CCWGG motifs on the 13 *K. pneumoniae* genomes and random generated genomes**

The orange histograms show the density distribution of GATC/CCWGG motifs on the random generated genomes. The green histograms show the density distribution of GATC/CCWGG on the 13 *K. pneumoniae* genomes.

**Figure S11**

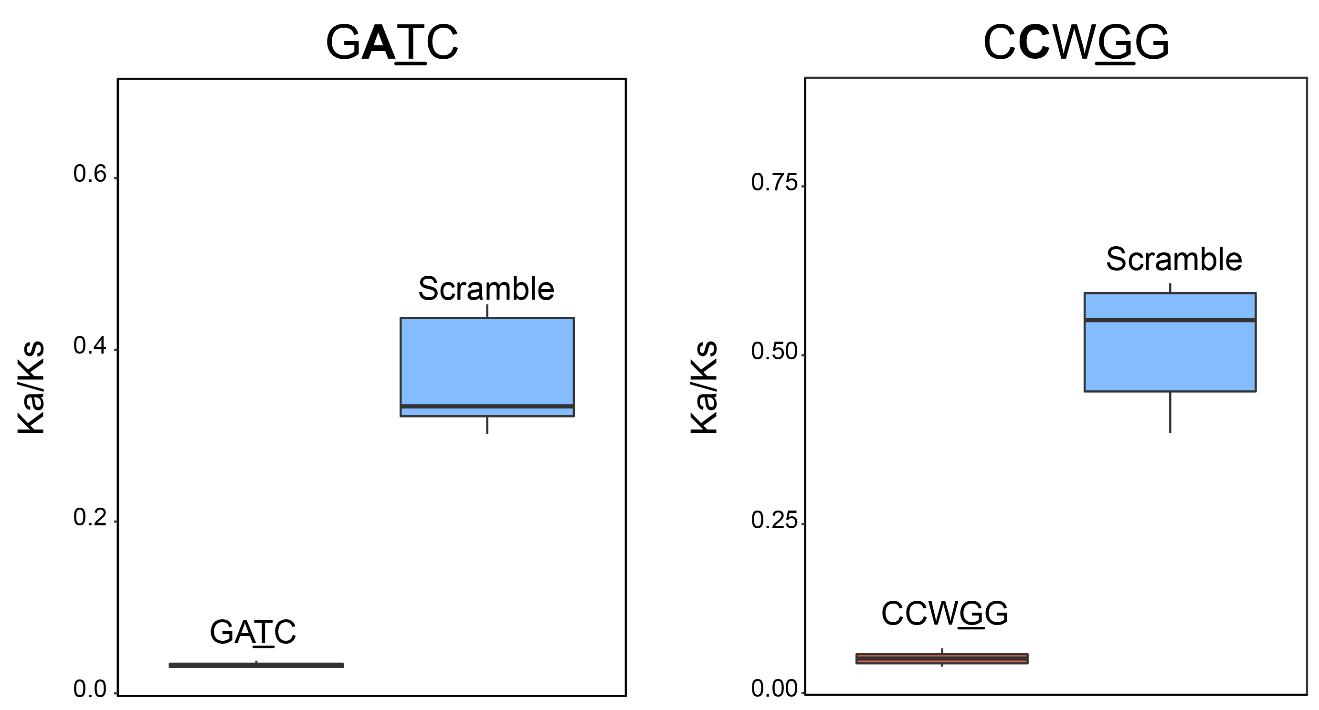

**Figure S11 Ka/Ks ratios for GATC/CCWGG “motif sequences” and “scramble sequences” of the 14 *K. pneumoniae* genomes**

Boxplots showing the Ka/Ks ratios of G**A**TC/C**C**WGG “motif sequences” of the *K. pneumoniae* genomes. Red boxes indicate the Ka/Ks ratios of G**A**TC/C**C**WGG “motif sequences”. Blue boxes indicate the corresponding “scramble sequences” as the controls.

**Figure S2**

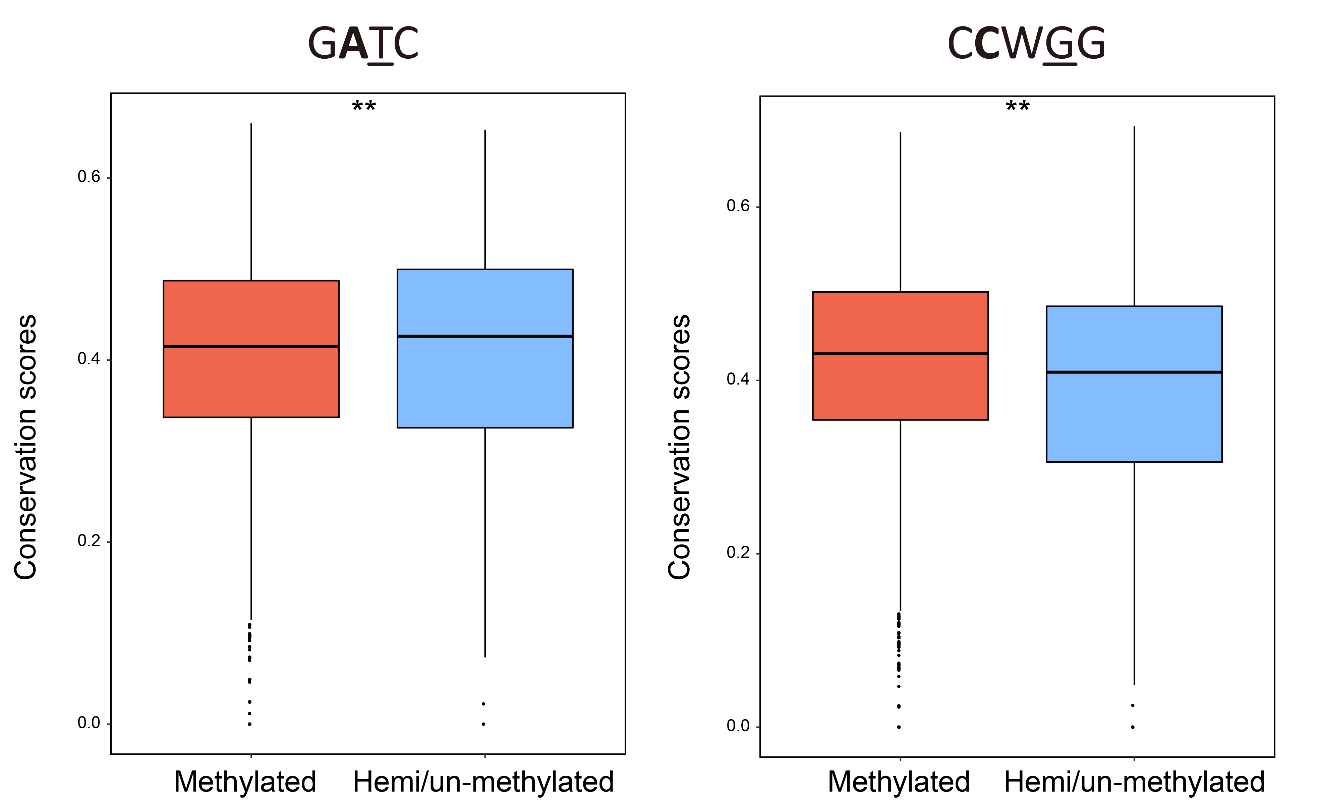

**Figure S12 Boxplots of conservations scores for methylated and hemi/un-methylated motifs and their flanking regions in IGRs**

Boxplots show the conservation values of G**A**TC/C**C**WGG motifs of the *K. pneumoniae* genomes. Red boxes indicate the conservation values of methylated G**A**TC/C**C**WGG motifs and their flanking regions (20 nt) in IGRs; blue boxes indicate the hemi/un-methylated motifs and their flanking regions (20 nt) as the controls in IGRs.

**Figure S13**

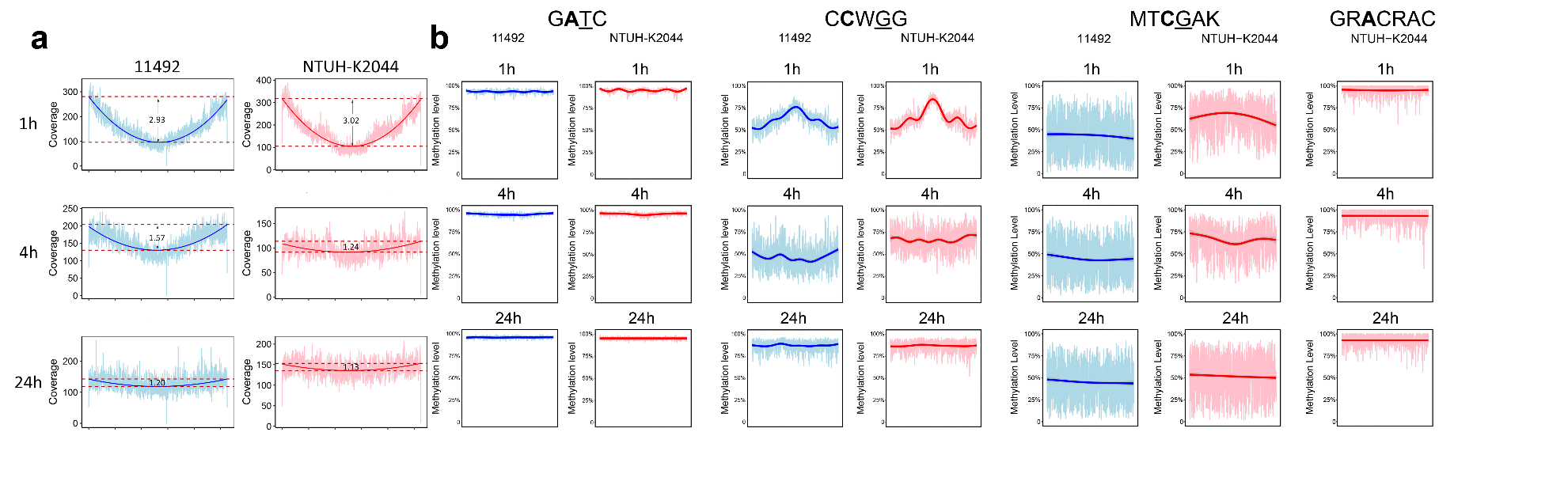

**Figure S13 Genome-wide sequencing coverage and methylation level of GATC and CCWGG motifs during cell cycles of two *K. pneumonia* strains**

**(a)** Genome-wide sequencing coverage versus genome position at three stages (1, 4 and 24 h) in the cell cycle of two *K. pneumonia* strains. The replication bidirectionally begins from the origin (O) and completes at terminus (T) (i. e., doubling point: in the middle of genome). The bold lines approximate the average coverage across the genomes. Ratios of the average coverage at oriC to that at doubling point are labeled in the figure. **(b)** Genome-wide methylation level versus genome position for the two motifs (G**A**TC, C**C**WGG, MT**C**GAK and GR**A**CRAC) at the three growth stages in the cell cycle of two *K. pneumonia* strains. The bold lines approximate the average methylation levels across the genomes (5kb window size).

**Figure S14**

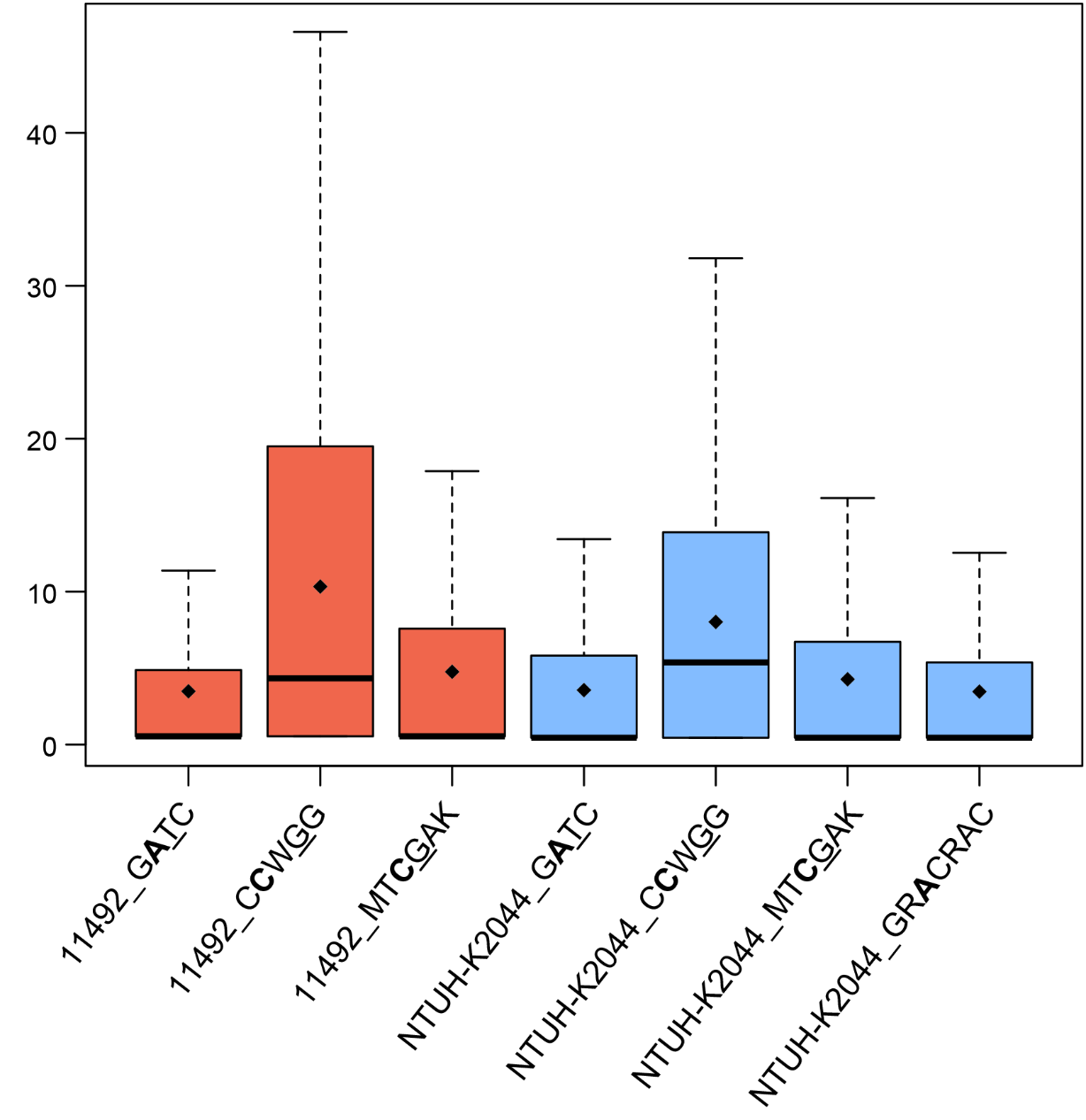

### Figure S14 Boxplots showing the re-methylation time of motifs in strain 11492 (red) and NTUH-K2044 (blue)

**Figure S15**

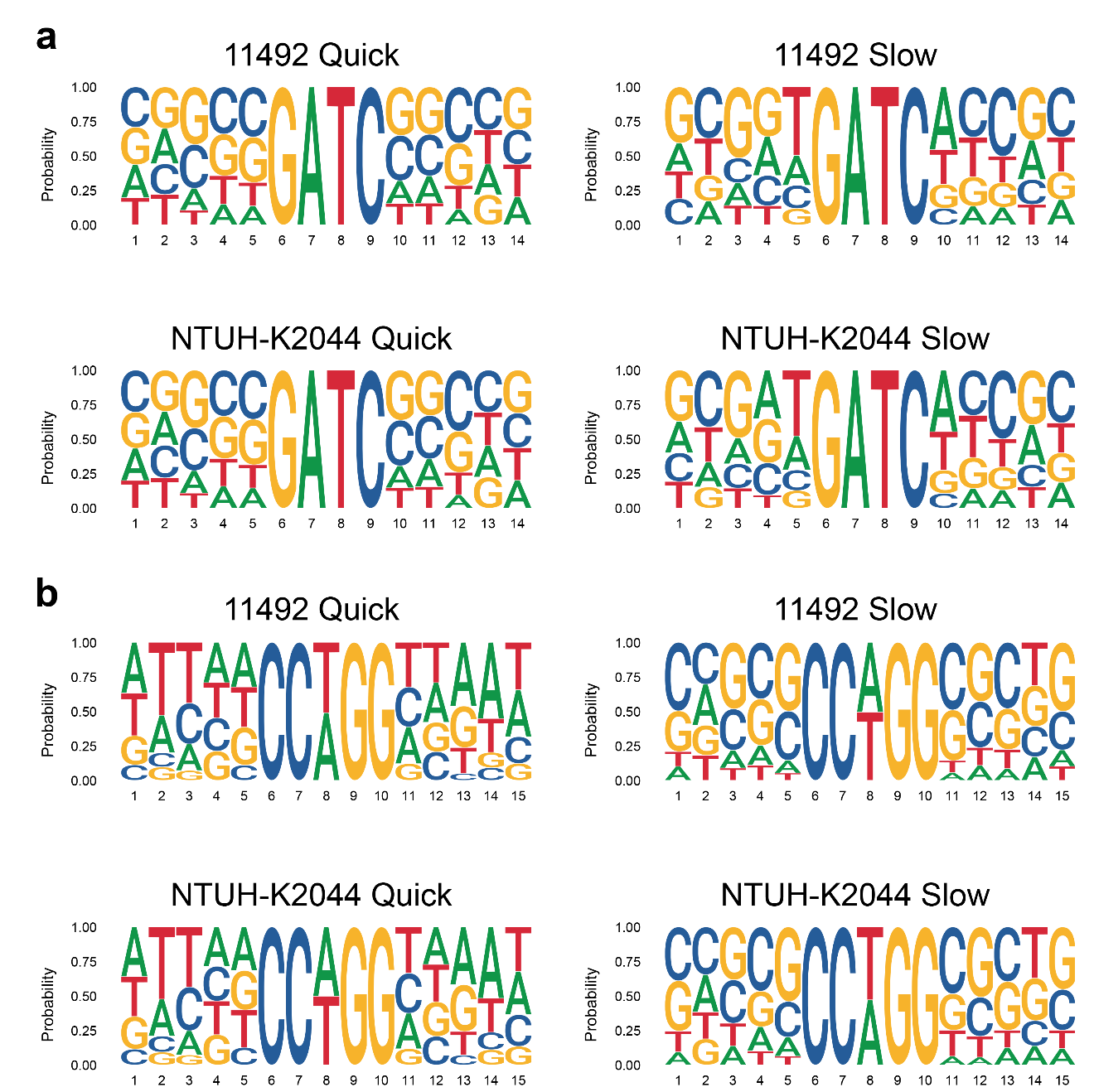

**Figure S15 Preferred sequences flanking the GATC and CCWGG motifs with fast and slow methylation rates in the two *K. pneumoniae* strains.**

**(a)** Preferred sequences flanking G**A**TC motifs with fast and slow methylation rates. The fast methylation rate means that the FRAC values for the motif are more than 0.95; the slow methylation rate means that the FRAC values for the motif are less than 0.9. Left panel: X-axis shows the G**A**TC motif flanked by five nucleotides; Y-axis indicates the ratios of A/T/C/G. Right panel: the sequence logos show preferences of five nucleotides flanking the GATC motifs at stationary phase in strain NTUH-K2044 and 11492. **(b)** Preference of flanking nucleotides of C**C**WGG motif. Quick methylation mode was defined as the FRAC values for 1 h, 4 h, and 24 h samples were all above 0.85, and the slow methylation mode was defined as the FRAC values for 1 h, 4 h, and 24 h samples were all below 0.55. X-axis indicated the positions of nucleotides, and Y-axis indicated the ratios of A/T/C/G.

**Figure S16**

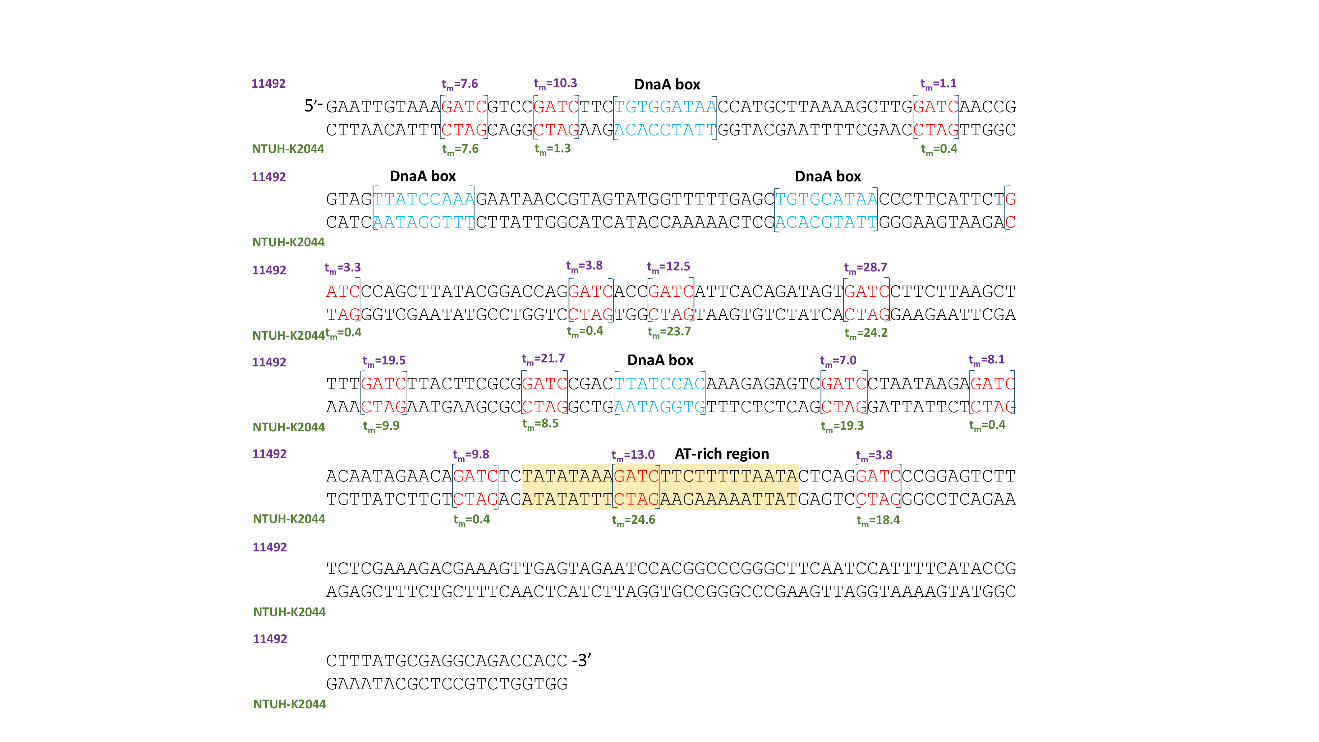

**Figure S15 Re-methylation time of GATC motifs in the oriC region (381 nt) of two *K. pneumoniae* strains**

The DnaA boxes (blue), AT-rich region (yellow) and G**A**TC motifs (red) are labeled on the sequence. The re-methylation times of GATC motifs in 11492 and NTUH-K2044 stains are marked in purple and green.

**Figure S17**

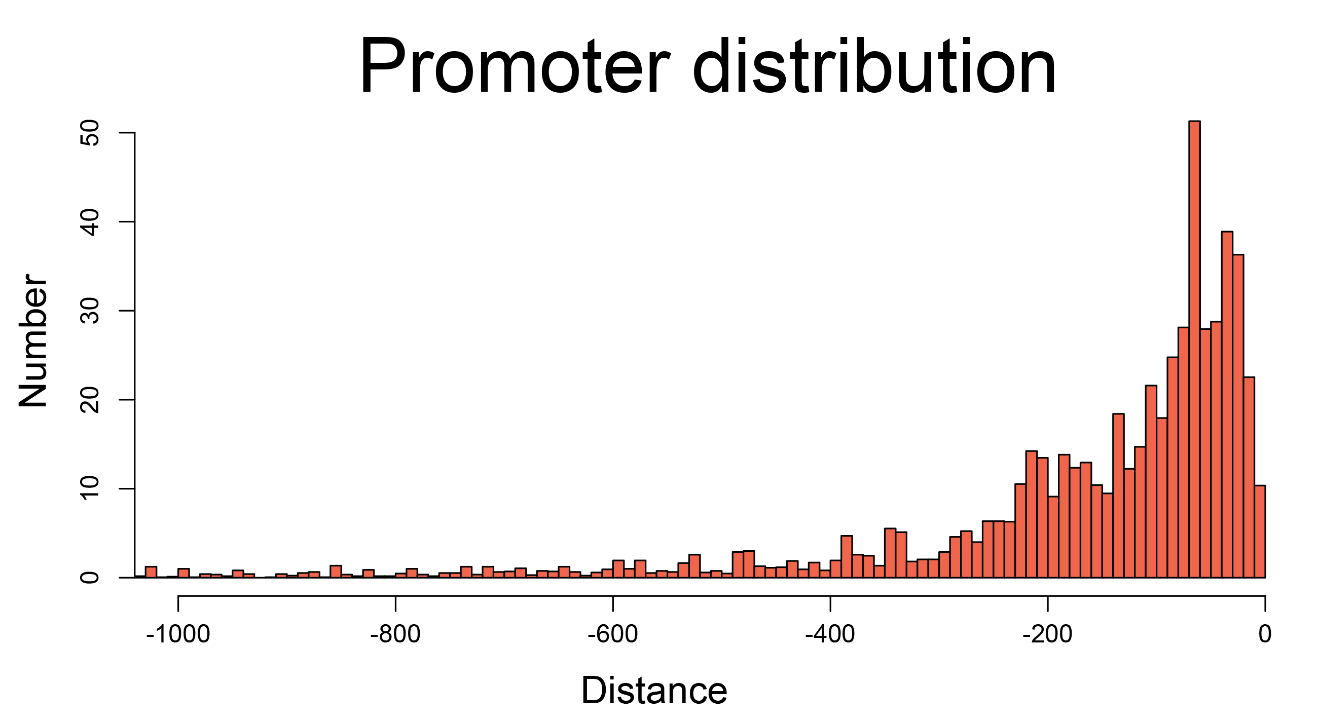

**Figure S16 Distribution of the promoters in the upstream regions of 14 *K. pneumoniae* genomes**

X-axis shows the distance from the start codon; Y-axis shows the number of promoters locating corresponding positions of the region.

**Figure S18**

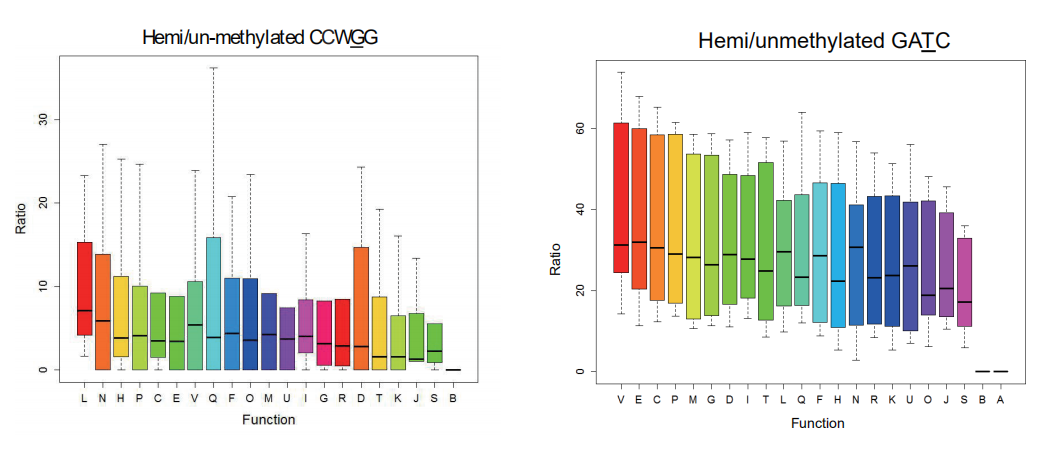

**Figure S18** **COG distributions of genes with hemi/un-methylated GATC/CCWGG motifs in intergenic regions**

Box plots indicated the COG categories of coding genes with methylated and hemi/un-methylated G**A**TC**/**C**C**WGG motifs in intergenic regions. X-axis shows the functional classes. Y-axis shows the ratio of genes in each functional class.

**Figure S19**

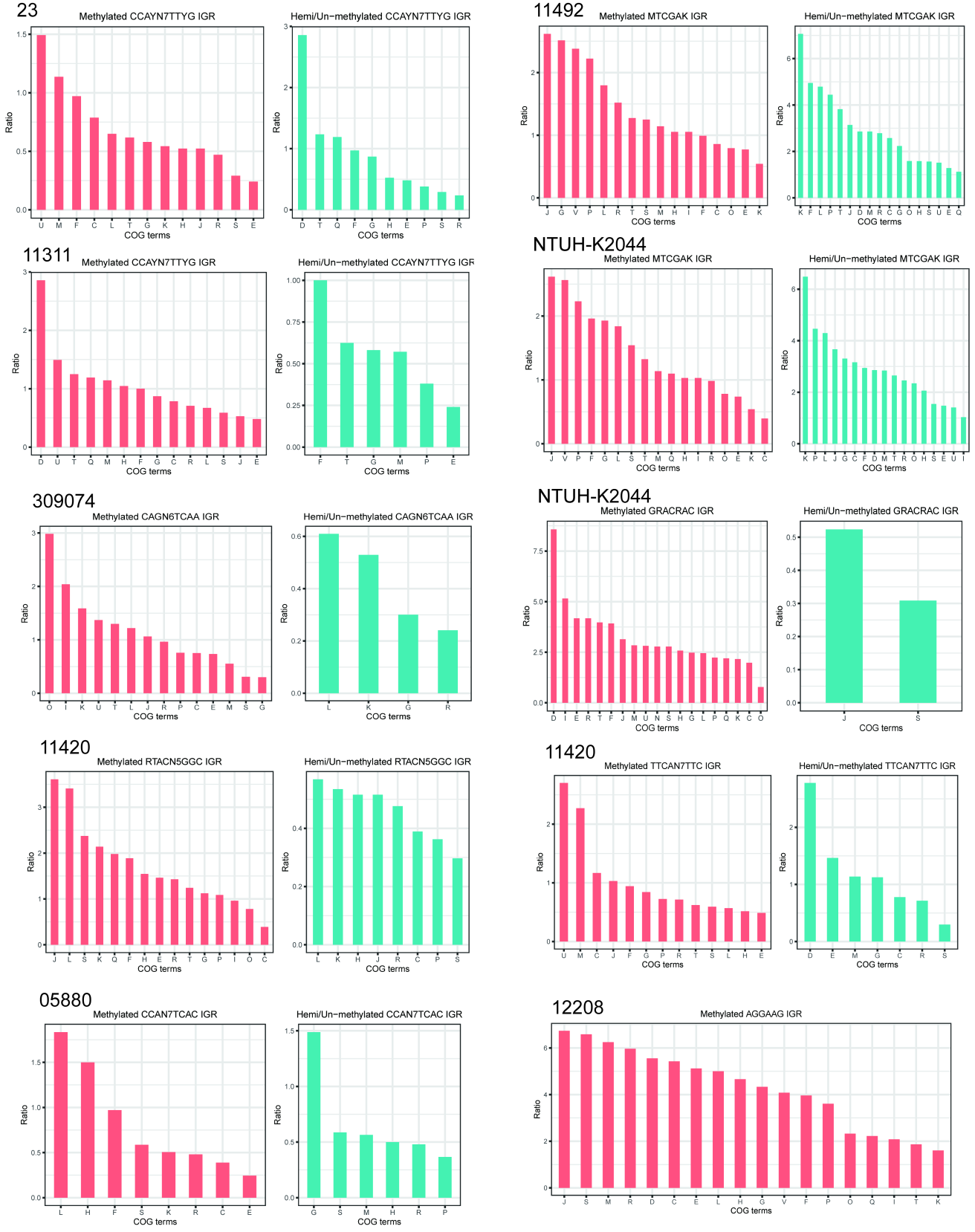
**Figure S19 COG distributions of genes with upstream hemi/un-methylated sites of novel motifs in intergenic regions**

X-axis shows the functional classes. Y-axis shows the ratio of genes in each functional class. [C] Energy production and conversion, [D] Cell cycle control, cell division, chromosome partitioning, [E] Amino acid transport and metabolism, [F] Nucleotide transport and metabolism, [G] Carbohydrate transport and metabolism, [H] Coenzyme transport and metabolism, [I] Lipid transport and metabolism, [J] Translation, ribosomal structure and biogenesis, [K] Transcription, [L] Replication, recombination and repair, [M] Cell wall/membrane/envelope biogenesis, [N] Cell motility, [O] Post-translational modification, protein turnover, and chaperones, [P] Inorganic ion transport and metabolism, [Q] Secondary metabolites biosynthesis, transport, and catabolism, [R] General function prediction only, [S] Function unknown, [T] Signal transduction mechanisms, [U] Intracellular trafficking, secretion, and vesicular transport, [V] Defense mechanisms.

**Figure S20**

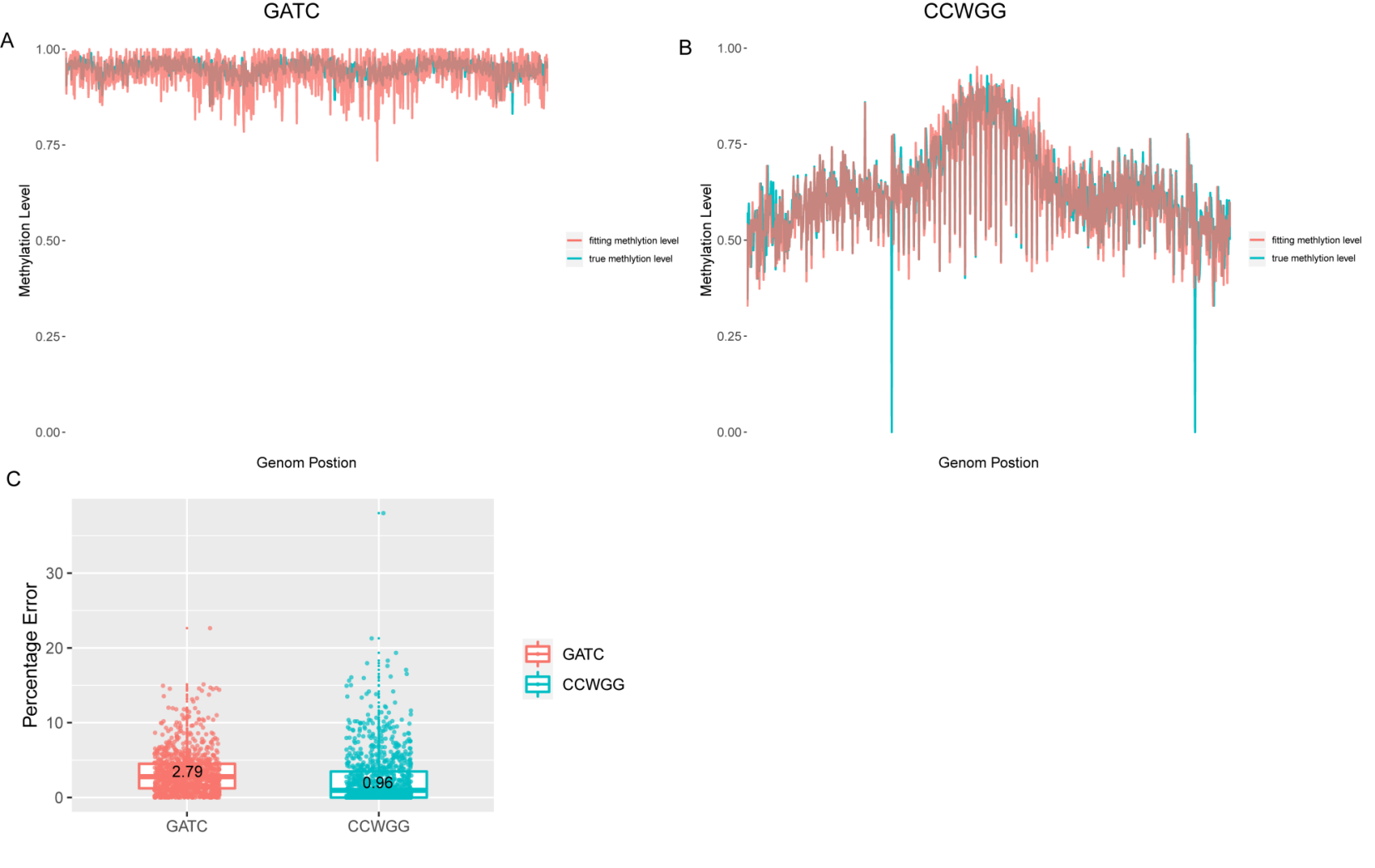

**Figure S20 Evaluating the accuracy of the re-modification time model**

**(A)** Fitting methylated-read ratio (FRACmodel) and real methylated-read ratio (FRACPacBio) comparison of GATC motif (5kb window). **(B)** Fitting methylated-read ratio (FRACmodel) and real methylated-read ratio (FRACPacBio) comparison of CCWGG motif (5k window). **(C)** Boxplot showing the percentage errors of GATC and CCWGG motifs.

**Supplementary tables**

### Table S1 Clinical information of 14 *K. pneumoniae* isolates

| **Strains No.** | **Samples** | **Capsular**  **type** | **MLST** | **Clonal group (CG)** | **Strains string test** | **Drug [sensibility](C:/Program%20Files%20(x86)/Youdao/Dict/7.3.0.0817/resultui/dict/?keyword=sensibility)*^a^*** | | | | |
| --- | --- | --- | --- | --- | --- | --- | --- | --- | --- | --- |
|  |  |  |  |  |  | **CFZ** | **CAZ** | **IPM** | **CIP** | **AK** |
| NTUH-K2044 | abscess/meningitis | K1 | 23 | 23 | + | I | S | I | S | S |
| 11492 | blood/abscess | K1 | 23 | 23 | + | R | R | R | R | S |
| 11420 | [ascites](C:/Program%20Files%20(x86)/Youdao/Dict/7.3.0.0817/resultui/dict/?keyword=ascites)/sputum | K1 | 1265 | 23 | + | R | R | R | S | S |
| 11454 | blood/abscess | K2 | 86 | / | + | S | S | S | S | S |
| 12208 | sputum | K54 | 4 | 29 | - | R | R | R | S | S |
| 11311 | blood/abscess | K57 | 412 | / | + | S | S | S | S | S |
| 23 | blood/abscess | K57 | 412 | / | + | S | S | S | S | S |
| 11305 | blood/cervical secretion | K64 | 38 | 147 | + | R | R | R | R | S |
| N201205880 | sputum | K21 | 86 | / | - | R | R | R | R | R |
| 309074 | sputum | K24 | 10 | / | - | R | R | R | R | R |
| 13190 | sputum | K27 | 40 | 147 | - | R | R | R | R | R |
| 283747 | sputum | K47 | 4 | 258 | - | R | R | R | R | R |
| 721005 | [urine](C:/Program%20Files%20(x86)/Youdao/Dict/7.3.0.0817/resultui/dict/?keyword=urine) | K47 | 4 | 258 | - | R | R | R | R | R |
| 11021 | [urine](C:/Program%20Files%20(x86)/Youdao/Dict/7.3.0.0817/resultui/dict/?keyword=urine) | K47 | 4 | 258 | - | R | R | R | R | R |

*Note: ^a^* CFZ: cefazolin; CAZ: ceftazidime; IPM: [imipenem](C:/Program%20Files%20(x86)/Youdao/Dict/7.3.0.0817/resultui/dict/?keyword=imipenem); CIP: [ciprofloxacin](C:/Program%20Files%20(x86)/Youdao/Dict/7.3.0.0817/resultui/dict/javascript:;); AK: [amikacin](C:/Program%20Files%20(x86)/Youdao/Dict/7.3.0.0817/resultui/dict/?keyword=amikacin); S: susceptible; R: resistant; I: intermediate.

### Table S2 Sequencing data of the 14 *K. pneumoniae* strains using SMRT technology

| **Sample** | **Cell** | **Number of Bases (bp)** | **Mean Read Length (bp)** | **Mean Subread length (bp)** | **Coverage** |
| --- | --- | --- | --- | --- | --- |
| NTUH-K2044 | 1 | 886,769,768 | 15,421 | 12,257 | 117X |
| 11492 | 2 | 952,838,022 | 8,164 | 7,142 | 165X |
| 11420 | 1 | 541,500,849 | 10,787 | 3,374 | 80X |
| 11454 | 2 | 773,967,122 | 8,457 | 2,732 | 95X |
| 12208 | 1 | 892,071,700 | 13,096 | 7,507 | 134X |
| 11311 | 1 | 891,654,097 | 11,116 | 7,490 | 124X |
| 23 | 2 | 737,867,481 | 7,954 | 2,862 | 82X |
| 11305 | 1 | 666,166,026 | 9,720 | 7,355 | 103X |
| N201205880 | 1 | 615,336,756 | 14,233 | 7,527 | 60X |
| 309074 | 1 | 1,101,489,716 | 12,287 | 8,500 | 160X |
| 13190 | 1 | 952,259,010 | 14,438 | 9,004 | 67X |
| 283747 | 1 | 623,152,844 | 14,379 | 8,926 | 95X |
| 721005 | 1 | 1,152,468,329 | 12,851 | 8,130 | 151X |
| 11021 | 1 | 522,202,437 | 14,804 | 7,477 | 55X |

### Table S3 SMRT Sequencing data of the samples at three growth time points (1, 4 and 24 h) of two *K. pneumoniae* strains (11492 and NTUH-K2044)

| **Sample** | **Cell** | **Number of Bases (bp)** | **Mean Read Length (bp)** | **Mean Subread length (bp)** | **Coverage** |
| --- | --- | --- | --- | --- | --- |
| 11492_1h | 1 | 1,463,421,074 | 14,095 | 8,209 | 160X |
| 11492_4h | 1 | 1,290,931,013 | 12,382 | 9,168 | 159X |
| 11492_24h | 1 | 1,061,611,880 | 10,129 | 6,510 | 132X |
| NTUH-K2044_1h | 1 | 1,446,229,850 | 14,195 | 8,384 | 177X |
| NTUH-K2044_4h | 1 | 681,653,271 | 11,213 | 9,378 | 100X |
| NTUH-K2044_24h | 1 | 1,446,068,018 | 13,125 | 9,085 | 145X |

### Table S4 Sequencing data of the 14 *K. pneumoniae* strains using Illumina Hiseq platform

| **Sample** | **Clean**  **Reads** | **Duplication**  **Deleted** | **Filter** | **Mate Paired** | **Mapped** | **Unique Mapped** | **Depth** | **Coverage**  **(10X)** |
| --- | --- | --- | --- | --- | --- | --- | --- | --- |
| NTUH-K2044 | 22,870,446 | 20,436,158 | 19,919,947 | 19,425,742 | 18,118,021 | 17,934,586 | 493 | 99.41% |
| 11492 | 9,548,040 | 8,274,524 | 8,203,572 | 8,140,190 | 7,962,700 | 7,874,315 | 218 | 98.96% |
| 11420 | 8,535,078 | 8,520,932 | 8,247,425 | 8,044,246 | 7,849,334 | 7,753,861 | 168 | 99.22% |
| 11454 | 15,106,774 | 15,037,602 | 14,464,415 | 13,990,144 | 13,921,432 | 13,755,535 | 317 | 99.20% |
| 12208 | 5,809,476 | 5,734,298 | 5,476,698 | 4,906,182 | 4,906,182 | 4,719,438 | 126 | 99.89% |
| 11311 | 7,722,008 | 6,170,366 | 6,063,244 | 5,960,572 | 5,858,494 | 5,797,267 | 160 | 99.21% |
| 23 | 8,726,760 | 8,706,494 | 8,385,358 | 8,135,502 | 8,024,562 | 7,935,682 | 177 | 99.19% |
| 11305 | 9,133,612 | 7,855,376 | 7,789,872 | 7,730,848 | 7,315,970 | 7,224,924 | 202 | 99.04% |
| N201205880 | 7,448,260 | 7,334,122 | 7,277,008 | 7,128,778 | 7,128,718 | 6,762,212 | 243 | 99.99% |
| 309074 | 10,483,754 | 9,279,180 | 9,253,276 | 9,004,180 | 9,004,180 | 6,745,943 | 286 | 95.31% |
| 13190 | 8,748,960 | 8,589,348 | 8,121,571 | 7,853,518 | 5,482,369 | 5,369,274 | 144 | 98.45% |
| 283747 | 10,371,170 | 10,225,530 | 9,890,486 | 9,312,452 | 9,312,452 | 8,885,307 | 239 | 99.99% |
| 721005 | 6,790,294 | 6,739,216 | 6,715,998 | 6,715,998 | 6,482,828 | 6,147,004 | 245 | 99.05% |
| 11021 | 7,191,852 | 7,102,130 | 7,045,938 | 6,902,228 | 6,902,228 | 6,592,975 | 246 | 99.99% |

### Table S5 Genomic information of the online 62 *K. pneumoniae* strains (NCBI) used in the phylogenetic tree

| **Strain** | **Accession** | **Size (Mb)** | **GC%** | **Genes** | **Proteins** | **Plasmids** | **Host** | **Isolate** | **City** | **Country** |
| --- | --- | --- | --- | --- | --- | --- | --- | --- | --- | --- |
| ATCC.43816.KPPR1 | NZ_CP009208 | 5.37483 | 57.4 | 5259 | 5074 | 0 | clinical | - | - | USA |
| ATCC.BAA.2146 | NZ_CP006659 | 5.7815 | 56.9738 | 5883 | 5705 | 4 | patients | urine | - | USA |
| blaNDM.1 | NZ_CP009114 | 5.51033 | 57.2905 | 5555 | 5378 | 2 | patients | blood or any other sterile source | - | - |
| BR | NZ_CP015990 | 5.42072 | 57.3423 | 5308 | 5102 | 1 | Homo sapiens | - | hangzhou | China |
| CAV1193 | NZ_CP013322 | 5.8747 | 57.0929 | 5944 | 5735 | 5 | Homo sapiens | ventilated airway | - | - |
| CAV1344 | NZ_CP011624 | 5.81001 | 57.0961 | 5874 | 5666 | 5 | patients | - | - | - |
| CAV1392 | NZ_CP011578 | 5.55249 | 57.2436 | 5618 | 5444 | 3 | patients | - | - | - |
| CAV1596 | NZ_CP011647 | 5.62052 | 57.1393 | 5664 | 5506 | 4 | patients | - | - | - |
| CG43 | NC_022566 | 5.16686 | 57.6 | 5098 | 4920 | 0 | Homo sapiens | - | Taiwan | China |
| DHQP1002001 | CP016811 | 5.55846 | 57.3358 | 5584 | 5322 | 2 | Homo sapiens | Urine | California | USA |
| DMC1097 | NZ_CP011976 | 5.71069 | 57.0422 | 5770 | 5606 | 3 | human | blood | Metropolitan | - |
| ED2 | NZ_CP016813 | 5.41296 | 57.4 | 5258 | 5076 | 0 | Homo sapiens | blood | Taiwan | China |
| ED23 | NZ_CP016814 | 5.5874 | 57.3144 | 5514 | 5311 | 1 | Homo sapiens | blood | Taiwan | China |
| HS11286 | NC_016845 | 5.68232 | 57.1373 | 5867 | 5779 | 6 | human | sputum | Shanghai | China |
| JM45 | NC_022082 | 5.60317 | 57.2405 | 5669 | 5387 | 2 | human | blood | - | - |
| Kp1084 | NC_018522 | 5.38671 | 57.4 | 5227 | 4995 | 0 | patients | - | Taiwan | China |
| Kp11 | CP016923 | 5.8056 | 56.7804 | 5707 | 5492 | 4 | Homo sapiens | Urine | Pennsylvania | USA |
| Kp1158 | NZ_CP006722 | 5.34104 | 57.2 | 5190 | 5026 | 0 | Homo sapiens | - | Taiwan | China |
| Kp13 | NZ_CP003999 | 5.73989 | 56.8079 | 5699 | 5486 | 6 | patients | - | - | Southern Brazil |
| Kp23 | CP016926 | 5.48385 | 57.1811 | 5421 | 5219 | 3 | Homo sapiens | Tracheal Aspirate | Minnesota | USA |
| Kp234.12 | NZ_CP011313 | 5.64883 | 56.7773 | 5597 | 5388 | 3 | Homo sapiens | blood | - | Germany |
| Kp30660 | NZ_CP006923 | 5.54094 | 57.146 | 5599 | 5419 | 5 | patients | - | - | - |
| Kp30684 | NZ_CP006918 | 5.41722 | 57.4316 | 5481 | 5324 | 3 | patients with urinary tract infections | - | - | - |
| Kp32192 | NZ_CP010361 | 5.6208 | 57.2065 | 5697 | 5481 | 3 | Homo sapiens | Excreted bodily substance | - | USA |
| Kp34618 | NZ_CP010392 | 5.64991 | 57.2079 | 5714 | 5553 | 4 | Homo sapiens | Bodily fluid | - | USA |
| KP36 | CP017385 | 5.75921 | - | 5671 | 5460 | 3 | Homo sapiens | urine | Taiwan: Tainan | China |
| Kp500.1420 | NZ_CP011980 | 5.54764 | 57.2924 | 5644 | 5468 | 4 | human | urine | North_East_Ohio | USA |
| Kp52.145 | NZ_FO834906 | 5.65568 | 57.0991 | 5574 | 5316 | 2 | Human | - | Java | - |
| KP617 | NZ_CP012753 | 5.69605 | 56.8612 | 5599 | 5411 | 2 | patients | - | Seoul | Korea |
| KpN01 | NZ_CP012987 | 5.68063 | 56.9887 | 5710 | 5505 | 4 | Homo sapiens | urine | Calgary | Canada |
| KpN06 | NZ_CP012992 | 5.65049 | 57.0185 | 5682 | 5477 | 4 | Homo sapiens | blood | Calgary | Canada |
| KPNIH1 | NZ_CP008827 | 5.76661 | 57.1432 | 5868 | 5711 | 3 | patients | - | - | USA |
| KPNIH10 | NZ_CP007727 | 5.76782 | 57.1433 | 5867 | 5708 | 3 | patients | - | New York | USA |
| KPNIH24 | NZ_CP008797 | 5.73456 | 57.1176 | 5815 | 5651 | 3 | Patients | throat/groin | - | - |
| KPNIH27 | NZ_CP007731 | 6.13244 | 56.7058 | 6188 | 5950 | 5 | Patients | groin | - | - |
| KPNIH29 | NZ_CP009863 | 5.5206 | 57.1769 | 5513 | 5297 | 2 | patients | perirectal | - | - |
| KPNIH30 | NZ_CP009872 | 5.48415 | 57.2625 | 5568 | 5418 | 3 | patients | perirectal | - | - |
| KPNIH31 | NZ_CP009876 | 5.47358 | 57.2555 | 5444 | 5254 | 3 | patients | urine | - | - |
| KPNIH32 | NZ_CP009775 | 5.85831 | 57.007 | 5906 | 5739 | 3 | patients | perirectal | - | - |
| KPNIH33 | NZ_CP009771 | 5.69522 | 57.2179 | 5707 | 5540 | 3 | patients | urine | - | - |
| KPNIH36 | NZ_CP014647 | 5.68147 | 57.207 | 5793 | 5633 | 3 | patient | rectal/perirectal | - | USA |
| KPNIH39 | NZ_CP014762 | 5.77967 | 56.9455 | 5800 | 5585 | 3 | Homo sapiens | - | - | USA |
| KPR0928 | NZ_CP008831 | 5.43679 | 57.3174 | 5512 | 5365 | 2 | Patients | sputum | - | - |
| MGH.78578 | NC_009648 | 5.69489 | 57.1666 | 5665 | 5450 | 5 | Patients | sputum | - | - |
| MS6671 | NZ_LN824133 | 5.9802 | 56.5426 | 5954 | 5749 | 6 | Homo sapiens | urine | - | - |
| NUHL24835 | NZ_CP014004 | 5.53519 | 57.2067 | 5466 | 5293 | 2 | - | urine | - | - |
| PittNDM01 | NZ_CP006798 | 5.8123 | 56.7812 | 5712 | 5511 | 4 | patients | urine | Pittsburgh, PA | USA |
| PMK1 | NZ_CP008929 | 5.99074 | 56.5508 | 5879 | 5672 | 4 | the first infected neonatal case | - | - | - |
| RJF293 | NZ_CP014008 | 5.45059 | 57.1955 | 5338 | 5144 | 1 | Homo sapiens | blood | Shanghai | China |
| RJF999 | NZ_CP014010 | 5.69083 | 57.1064 | 5572 | 5375 | 1 | Homo sapiens | blood | Shanghai | China |
| SKGH01 | NZ_CP015500 | 6.08846 | 56.5414 | 6035 | 5778 | 5 | Homo sapiens | Urine | - | United Arab Emirates |
| TGH10 | NZ_CP012744 | 5.35546 | 57.3 | 5379 | 5158 | 0 | Homo sapiens | wound swab | Greece | - |
| TGH13 | CP012745 | 5.34458 | 57.4 | 5316 | 5125 | 0 | Homo sapiens | rectal | - | Greece |
| TGH8 | NZ_CP012743 | 5.43972 | 57 | 5436 | 5208 | 0 | Homo sapiens | CVC | Greece | - |
| TH1 | NZ_CP016159 | 5.49547 | 57.1624 | 5411 | 5218 | 2 | human | fecal | Beijing | China |
| UCLAOXA232KP.Pt0 | NZ_CP012560 | 5.31274 | 57.5 | 5294 | 5081 | 0 | Homo sapiens | Respiratory | California | USA |
| UCLAOXA232KP.Pt1 | CP012561 | 5.88049 | 56.9253 | 5921 | 5658 | 6 | Homo sapiens | abdominal drainage | California | USA |
| UCLAOXA232KP.PtX | CP012568 | 5.79855 | 57.018 | 5819 | 5573 | 4 | Homo sapiens | Rectal | California | USA |
| UHKPC07 | NZ_CP011985 | 5.52378 | 57.3213 | 5615 | 5462 | 3 | human | urine | - | UH |
| UHKPC33 | NZ_CP011989 | 5.66312 | 57.1333 | 5762 | 5581 | 4 | human | urine | Cleveland_OH | USA |
| W14 | NZ_CP015753 | 5.49548 | 57.1624 | 5407 | 5213 | 2 | Homo sapiens | fecal | Beijing | China |
| XH209 | NZ_CP009461 | 5.11888 | 57.6 | 5054 | 4889 | 0 | patient | blood | Hangzhou, Zhejiang | China |

### Table S6 Bisulfite sequencing data of the 14 *K. pneumoniae* strains

| **Strain No.** | **Clean pair reads*^a^*** | **Genome depth (X)** | **BS-con Rate*^b^*** | **Number of C**  **(≥ 20X)** | **C covered*^c^***  **(≥ 20X)** | **Number of methylated C*^d^* (≥ 20X)** |
| --- | --- | --- | --- | --- | --- | --- |
| NTUH-K2044 | 5,120,161 | 190 | 99.96% | 1,558,949 | 99.37% | 19,489/19,457 |
| 11492 | 7,816,589 | 309 | 99.88% | 1,542,407 | 99.10% | 21,570/21,574 |
| 11420 | 4,961,543 | 184 | 99.97% | 1,628,192 | 99.24% | 19,531/19,524 |
| 11454 | 6,801,540 | 263 | 99.45% | 1,545,707 | 99.17% | 16,146/16,195 |
| 12208 | 6,329,978 | 213 | 99.99% | 1,613,576 | 98.70% | 19,903/19,884 |
| 11311 | 9,262,793 | 389 | 99.94% | 1,557,774 | 99.27% | 19,283/19,276 |
| 23 | 3,902,961 | 146 | 99.99% | 1,586,682 | 99.27% | 19,365/19,382 |
| 11305 | 4,704,639 | 177 | 99.92% | 1,532,062 | 99.07% | 19,096/19,102 |
| N201205880 | 8,519,543 | 270 | 100% | 1,707,387 | 98.79% | 20,891/20,875 |
| 309074 | 8,439,345 | 354 | 99.97% | 1,594,371 | 99.19% | 18,185/18,207 |
| 13190 | 6,286,874 | 269 | 99.72% | 1,585,130 | 98.08% | 19,694/19,635 |
| 283747 | 6,093,694 | 190 | 99.90% | 1,597,379 | 98.68% | 19,431/19,424 |
| 721005 | 8,553,164 | 244 | 99.60% | 1,665,462 | 98.46% | 20,578/20,541 |
| 11021 | 6,137,777 | 197 | 99.98% | 1,644,827 | 98.48% | 19,059/19,003 |

*Note: ^a^* “Clean pair reads” represents the paired reads mapped uniquely to the reference genome by Bismark. *^b^* “BS-con Rate” indicates the bisulfite conversion rate. *^c^* “C covered” indicates the proportion of mapped C sites (≥ 20X) over total C sites in the reference genome. *^d^* The number of methylated C on the plus and minus strands of chromosomes and plasimds.

### Table S7 Bisulfite sequencing data of the samples at three growth time points (1, 4 and 24 h) of two *K. pneumoniae* strains (11492 and NTUH-K2044)

| Sample | Clean pair  Reads^a^ | Genome depth (X) | BS-con Rate^b^ | Number of C (≥ 20X) | C covered^c^ (≥ 20X) | Number of methylated  C^d^ (≥ 20X) |
| --- | --- | --- | --- | --- | --- | --- |
| 11492_1h | 53,139,694 | 2706X | 99.36% | 1,542,690 | 99.12% | 25,054/24,943 |
| 11492_4h | 40,388,108 | 2061X | 99.83% | 1,543,117 | 99.15% | 23,525/23,466 |
| 11492_24h | 45,586,499 | 2299X | 98.89% | 1,543,566 | 99.18% | 24,713/24,609 |
| NTUH-K2044_1h | 6,049,372 | 245X | 100% | 1,501,581 | 95.59% | 20,435/20,414 |
| NTUH-K2044_4h | 5,266,802 | 223X | 100% | 1,498,879 | 95.42% | 16,896/16,752 |
| NTUH-K2044_24h | 83,926,262 | 4168X | 98.63% | 1,551,123 | 98.75% | 29,835/29,882 |

*Note: ^a^* “Clean pair reads” represents the paired reads mapped uniquely to the reference genome by Bismark. *^b^* “BS-con Rate” indicates the bisulfite conversion rate. *^c^* “C covered” indicates the proportion of mapped C sites (≥ 20X) over total C sites in the reference genome. ^d^ The number of methylated C on the plus and minus strands of chromosomes and plasimds.

### Table S8 The 22 predicted MTase genes and the corresponding 15 methylated motifs

| R-**M Type** | **Candidate MTase genes*** | | **Strain** | **Motif** | | **Rebase** |
| --- | --- | --- | --- | --- | --- | --- |
| **Type II** | **K2044peg364/11492peg365/11420peg366/**  **11454peg356/12208/peg379/11311peg358/23peg361/**  **11305peg365/N201005880peg363/309074peg363/**  **13190peg374/283747peg368/721005peg375/**  **11021peg372** | **M1A** | **14 strains** | **GATC** | A | 100% aa identity to M.Kpn43816Dam |
| **Type I** | **11420peg4701.peg4700.peg4699** | **M2B1** | **11420** | **RTACN_5_GGC** | B1 | **New** |
| Type I | M/S.Eco448ORF25820P | M3B1 |  |  |  |  |
| Type II | M.KpnSWU01ORFJP | M4B1 |  |  |  |  |
| Type II | M.Sen39523DndAP | M5B1 |  |  |  |  |
| Type I | M1/M2/S.KpnHSL4ORFAP | M2B2 |  | **TTCAN_7_TTC** | B2 | **New** |
| **Type I** | **11420peg5619.peg5618** | **M3B2** |  |  |  |  |
| Type II | M.KpnSWU01ORFJP | M4B2 |  |  |  |  |
| Type II | M.Sen39523DndAP | M5B2 |  |  |  |  |
| **Type I** | **11305peg4529.peg4530** | **M6C** | **11305** | **CCAGN_7_RTTC** | C | 100% aa identity to M.KpnAATI |
| **Type I** | **23peg4539.peg4540/11311peg4473.peg4474** | **M7D** | **23/11311** | **CCAYN_7_TTYG** | D | **New** |
| Type II | M.Sen39523DndAP | M8D |  |  |  |  |
| **Type I** | **11454peg1670.peg1671** | **M9E** | **11454** | **AGCN_5_CTTC** | E | 99.9% aa identity to M.KpnGH01II |
| Type III | M.Kpn214ORFGP | M10E |  |  |  |  |
| Type II | M.Sen39523DndAP | M11E |  |  |  |  |
| **Type I** | **13190peg4519.peg4520** | **M12** | **13190** | **CCAGN_7_RTTC** | C | 100% aa identity to M.KpnAATI |
| **Type I** | **13190peg4583.peg4582** | **M13** |  | **GGCAN_8_TCG** | F | 100% aa identity to M.KpnAATIV |
| **Type II** | **12208peg4487** | **M14** | **12208** | **AGGAAG** | G | **New** |
| Type II | M.Sen39523DndAP | M15 |  |  |  |  |
| **Type I** | **N201205880peg5335.peg5336** | **M17** | **N201205880** | **CCAN_7_TCAC** | I | **New** |
| **Type I** | **309074peg650.peg649** | **M18** | **309074** | **CATCN_6_TTYG** | J | 100% aa identity to M.Kpn39795II |
| **Type I** | **309074peg4560.peg4561** | **M20** |  | **CTAN_5_GTAA** | M | 99.8% aa identity to M.Kpn35657I |
| **Type I** | **309074peg4841.peg4840** | **M21** |  | **CAGN_6_TCAA** | N | **New** |
| **Type II** | **K2044peg4432** | **M24** | **NTUH-K2044** | **GRACRAC** | P | **New** |
| **Type II** | **K2044peg4434/11492peg4298** | **M27** | **NTUH-K2044**  **/11492** | **MTCGAK** | R | **New** |
| **Type II** | **K2044peg1760/11492peg1825/11420peg1936/**  **11454peg1716/12208peg1747/11311peg1742/**  **23peg1802/11305peg1826/N201205880peg1726/**  **309074peg1868/13190peg1805/283747peg1927/**  **721005peg1933/11021peg1937** | **M32** | **14 strains** | **CCWGG** | V | 100% aa identity to M.Kpn62629II |

### Table S9 Oligonucleotide primers used in this study

| **Primer^a^** | **Target gene** | **5’-3’ sequence^b^** | **RE* for cloning** | **Methylation-sensitive RE for verification** |
| --- | --- | --- | --- | --- |
| pRRS_F |  | TCATTAGGCACCCCAGGCT | - | - |
| pRRS_R |  | TTCCCAGTCACGACGTTGTA | - | - |
| M2B1_F | 11420peg4699-peg4671 | CG CCTGCAGGTTAAGGTTAATCATATGACACTGATTAACCTAAAAGATCTCG | SbfI |  |
| M2B1_R |  | CG GGATCC**GCCTGCAGGTAT**TTAAACCTTAAAAATTTTTGTTAGCTGC | BamHI | BfuAI |
| M3B2_F | 11420peg5618-peg5619 | CG CCTGCAGGTTAAGGTTAATCATATGGCACAACAGACACACAC | SbfI |  |
| M3B2_R |  | CG GGATCCTTC**GAATTCCAATTGAA**TCAGTTGCTGGCGGTTTCTG | BamHI | MfeI, BstBI |
| M7D_F | 23peg4539-peg4540 | CC CGAAGACATAGCTTTAAGGTTAATCATATGGCCGCAATCTCTTTCGA | BbsI |  |
| M7D_R |  | CG CCTGCAGGTT**CGAATTCGCAGGTGG**TCATGCGCTGGCCTCCTGTG | SbfI | BfuAI, BstBI |
| M14G_F | 12208peg4490 | CG CCTGCAGGTTAAGGTTAATCATATGAACACTGCCAATTTAAA | SbfI |  |
| M14G_R |  | CG GGATCCAGTA**CTTCCT**TTAAGCAATAGCTGCTTT | BamHI | ScaI |
| M17I_F | 05880peg5452-peg5453 | CG CCTGCAGGTTAAGGTTAATCATATGAACGACAAAATCAGTC | SbfI |  |
| M17I_R |  | CG GGATCC**GTGAATAGTACTGG**TCATACGGCAACCGCCTCAT | BamHI | ScaI |
| M21N_F | 09074peg4167-peg4168 | CG CCTGCAGGTTAAGGTTAATCATATGGCCATTAAGAAAACCGA | SbfI |  |
| M21N_R |  | CG GGATCC**TTGACGAGTACTG**TCACTCATGAATAACCTCCT | BamHI | ScaI |
| M24P_F | NTUH-K2044peg619 | CG CCTGCAGGTTAAGGTTAATCATATGAGTTTAAATAACATTCA | SbfI |  |
| M24P_R |  | CG GGATCCACAT**GTTGTTC**CTAGAGATCGTTAATCC | BamHI | PciI |
| M27R_F | 11492peg4298 | CG CCTGCAGGTTAAGGTTAATCATATGAAAAAAGTATCTTGTGT | SbfI |  |
| M27R_R |  | CG GGATCC**ATCGAT**TCACTTTGTTGCGACCAAGT | BamHI | BspDI |
| M2_F |  | CG CCTGCAGG TTAAGGTTAATCATATGACTAATAAGAAAATCA | SbfI |  |
| M2B2_R |  | CG GGATCC **GAATTC CAATTGAA**TTAAACCTTAAAAATTTTTGTTAGCTGC | BamHI | MfeI, EcoRI |
| M3_F |  | CG CCTGCAGG TTAAGGTTAATCATATGGCACAACAGACACACAC | SbfI |  |
| M3B1_R |  | CG GGATCC**GCCTGCAGGTAT**TCAGATCCTTCCACCCACCC | BamHI | BfuAI |
| M4_F |  | CG CCTGCAGG TTAAGG TTAATCATATGTCCCGATTTATCCAGGG | SbfI |  |
| M4B1_R |  | CG GGATCC **GCCTGCAGGTAT** TTACGCGGCCTCCGGGAACT | BamHI | BfuAI |
| M4B2_R |  | CG GGATCC **GAATTC CAATTGAA**TTACGCGGCCTCCGGGAACT | BamHI | MfeI, EcoRI |
| M5_F |  | CG CCTGCAGG TTAAGG TTAATCATATGAAATTACCGATTTACCT | SbfI |  |
| M5B1_R |  | CG GGATCC **GCCTGCAGGTAT** TTAGTGATGTGACCATTCAA | BamHI | BfuAI |
| M5B2_R |  | CG GGATCC **GAATTC CAATTGAA**TTAGTGATGTGACCATTCAA | BamHI | MfeI, EcoRI |
| M6C_F |  | CCC AAGCTT TTAAGGTTAATCATATGTCAATCAGCTCGGTTAT | HindIII |  |
| M6C_R |  | CG CCTGCAGG **GAATTCTAGTACTGG** TTAATTGATGGCGGCGTC | SbfI | ScaI, EcoRI |
| M8D_F |  | CG CCTGCAGG TTAAGG TTAATCATATGAAATTACCGATTTACCT | SbfI |  |
| M8D_R |  | CG GGATCC **CGAAGAG GCAGGTGG** TTAGTGATGTGACCATTCAA | BamHI | BfuAI, Bst6I |
| M9E_F |  | CG CCTGCAGG TTAAGGTTAATCATATGCTACAAAACAACCCTG | SbfI |  |
| M9E_R |  | CG GGATCC TTC**GAAGAGCTCGCTCC**GGATCATACCAAAACCGCCTTAA | BamHI | BspEI, BstBI |
| M10E_F |  | CG CCTGCAGG TTAAGG TTAATCATTTGGCCCCTATATTCTCGAT | SbfI |  |
| M10E_R |  | CG GGATCC TTC**GAAGAGCTCGCTCC**GGATCACCGCATCACCATGCTCT | BamHI | BspEI, BstBI |
| M11E_F |  | CG CCTGCAGG TTAAGG TTAATCATATGAAATTACCGATTTACCT | SbfI |  |
| M11E_R |  | CG GGATCC TTC**GAAGAGCTCGCTCC**GGATTAGTGATGTGACCATTCAA | BamHI | BspEI, BstBI |
| M12C_F |  | CCC AAGCTT TTAAGGTTAATCATATGTCAATCAGCTCGGTTAT | HindIII |  |
| M12C_R |  | CG CCTGCAGG TTCGAATTCTAGTACTGGTTAATTGATGGCGGCGTCGG | SbfI |  |
| M13F_F |  | CCA ATGCAT TTAAGGTTAATCATATGAGTGAGGGGAAATTGC | NsiI |  |
| M13F_R |  | GA AGATCT TCG**CGAGCCAGTACTGCC**TCATACCTTCACCTCACCAA | BglII | ScaI, NruI |
| M15G_F |  | CG CCTGCAGG TTAAGGTTAATCATATGAAATTACCGATTTACC | SbfI |  |
| M15G_R |  | CG GGATCC AGTA**CTTCCT**TTAGTGATGTGACCATTC | BamHI | ScaI |
| M18J_F |  | CG CCTGCAGG TTAAGGTTAATCATATGTCTCCCCAGATTGAAGC | SbfI |  |
| M18J_R |  | CG GGATCC TT**CGAATCGCGCGATG**CTATACGAACATTTGCTG | BamHI | BstBI, BtgZI |
| M20M_F |  | CG CCTGCAGG TTAAGGTTAATCATATGAGTTCTAAGTTTCGGAA | SbfI |  |
| M20M_R |  | CG GGATCC **TTACAGTACTAG**TTAGAACTCATAACCCAGCC | BamHI | ScaI |

*Note*: ^a^ F: forward primer; R: reverse primer. ^b^ The restriction sites used for cloning are underlined in black; the restriction sites used for verification of the methylated base are double-underlined in red. The specific recognition motifs of MTases are highlighted with the blue bold characters. ^*^ RE: Restriction endonucleases.

### Table S10 Detailed information of the motifs and corresponding DNA MTases among the 14 *K. pneumoniae* strains

| **Strain No.** | **Sequence motif^a^** | **MTase** | **Modification** | **Type^b^** | **Locus**  **M gene** | **Locus**  **S gene** | **Comments^c^** |
| --- | --- | --- | --- | --- | --- | --- | --- |
| NTUH-K2044 | G**A**TC | M.KpnK2044I (Dam) | 6mA | orphan | peg364 | - | 100% aa identity to M.Kpn43816Dam |
|  | C**C**WGG | M.KpnK2044II (Dcm) | 5mC | orphan | peg1760 | - | 100% aa identity to M.Kpn62629II |
|  | GR**A**CRAC | M.KpnK2044III (KamC) | 6mA | RM-II | peg4432 | - | New |
|  | MT**C**GAK | M.KpnK2044IV (KcmA) | 5mC | RM-II | peg4434 | - | New |
| 11492 | G**A**TC | M.Kpn11492I (Dam) | 6mA | orphan | peg365 | - | 100% aa identity to M.Kpn43816Dam |
|  | C**C**WGG | M.Kpn11492II (Dcm) | 5mC | orphan | peg1825 | - | 100% aa identity to M.Kpn62629II |
|  | MT**C**GAK | M.Kpn11492III (KcmA) | 5mC | RM-II | peg4298 | - | New |
| 11420 | G**A**TC | M.Kpn11420I (Dam) | 6mA | orphan | peg366 | - | 100% aa identity to M.Kpn43816Dam |
|  | C**C**WGG | M.Kpn11420II (Dcm) | 5mC | orphan | peg1936 | - | 100% aa identity to M.Kpn62629II |
|  | RT**A**CN_5_GGC | M1.Kpn11420III  M2.Kpn11420III (KamA) | 6mA | RM-I | peg4701 peg4700 | peg4699 | New |
|  | TTC**A**N_7_TTC | M.Kpn11420IV (KamB) | 6mA | RM-I | peg5619 | peg5618 | New |
| 11454 | G**A**TC | M.Kpn11454I (Dam) | 6mA | orphan | peg356 | - | 100% aa identity to M.Kpn43816Dam |
|  | C**C**WGG | M.Kpn11454 (Dcm) | 5mC | orphan | peg1716 | - | 100% aa identity to M.Kpn62629II |
|  | **A**GCN_5_CTTC | M.Kpn11454III | 6mA | ? | peg1670 | peg1671 | 99.9% aa identity to M.KpnGH01II |
| 12208 | G**A**TC | M.Kpn11208I (Dam) | 6mA | orphan | peg379 | - | 100% aa identity to M.Kpn43816Dam |
|  | C**C**WGG | M.Kpn11208II (Dcm) | 5mC | orphan | peg1747 | - | 100% aa identity to M.Kpn62629II |
|  | AGGA**A**G | M.Kpn12208III (KamD) | 6mA | RM-II | peg4487 | - | New |
| 11311 | G**A**TC | M.Kpn11311I (Dam) | 6mA | orphan | peg358 | - | 100% aa identity to M.Kpn43816Dam |
|  | C**C**WGG | M.Kpn11311II (Dcm) | 5mC | orphan | peg1742 | - | 100% aa identity to M.Kpn62629II |
|  | CC**A**YN_7_TTYG | M.Kpn11311III (KamE) | 6mA | RM-I | peg4473 | peg4474 | New |
| 23 | G**A**TC | M.Kpn23I (Dam) | 6mA | orphan | peg361 | - | 100% aa identity to M.Kpn43816Dam |
|  | C**C**WGG | M.Kpn23II (Dcm) | 5mC | orphan | peg1802 | - | 100% aa identity to M.Kpn62629II |
|  | CC**A**YN_7_TTYG | M.Kpn23III (KamE) | 6mA | RM-I | peg4539 | peg4540 | New |
| 11305 | G**A**TC | M.Kpn11305I (Dam) | 6mA | orphan | peg365 | - | 100% aa identity to M.Kpn43816Dam |
|  | C**C**WGG | M.Kpn11305II (Dcm) | 5mC | orphan | peg1826 | - | 100% aa identity to M.Kpn62629II |
|  | CC**A**GN_7_RTTC | M.Kpn11305III | 6mA | RM-I | peg4529 | peg4530 | 100% aa identity to M.KpnAATI |
| N201205880 | G**A**TC | M.Kpn05880I (Dam) | 6mA | orphan | peg363 | - | 100% aa identity to M.Kpn43816Dam |
|  | C**C**WGG | M.Kpn05880II (Dcm) | 5mC | orphan | peg1726 | - | 100% aa identity to M.Kpn62629II |
|  | CC**A**N_7_TCAC | M.Kpn05880III (KamG) | 6mA | RM-I | peg5335 | peg5336 | New |
| 309074 | G**A**TC | M.Kpn309074I (Dam) | 6mA | orphan | peg363 | - | 100% aa identity to M.Kpn43816Dam |
|  | C**C**WGG | M.Kpn309074II (Dcm) | 5mC | orphan | peg1868 | - | 100% aa identity to M.Kpn62629II |
|  | C**A**TCN_6_TTYG | M.Kpn309074III | 6mA | RM-I | peg650 | peg649 | 100% aa identity to M.Kpn39795II |
|  | CT**A**N_5_GTAA | M.Kpn309074IV | 6mA | RM-I | peg4560 | peg4561 | 99.8% aa identity to M.Kpn35657I |
|  | C**A**GN_6_TCAA | M.Kpn309074V (KamH) | 6mA | RM-I | peg4841 | peg4840 | New |
| 13190 | G**A**TC | M.Kpn13190I (Dam) | 6mA | orphan | peg374 | - | 100% aa identity to M.Kpn43816Dam |
|  | C**C**WGG | M.Kpn13190II (Dcm) | 5mC | orphan | peg1805 | - | 100% aa identity to M.Kpn62629II |
|  | CC**A**GN_7_RTTC | M.Kpn13190III | 6mA | RM-I | peg4519 | peg4520 | 100% aa identity to M.KpnAATI |
|  | GGC**A**N_8_TCG | M.Kpn13190IV | 6mA | RM-I | peg4583 | peg4582 | 100% aa identity to M.KpnAATIV |
| 283747 | G**A**TC | M.Kpn283747I (Dam) | 6mA | orphan | peg368 | - | 100% aa identity to M.Kpn43816Dam |
|  | C**C**WGG | M.Kpn283747II (Dcm) | 5mC | orphan | peg1927 | - | 100% aa identity to M.Kpn62629II |
| 721005 | G**A**TC | M.Kpn721005I (Dam) | 6mA | orphan | peg375 | - | 100% aa identity to M.Kpn43816Dam |
|  | C**C**WGG | M.Kpn721005II (Dcm) | 5mC | orphan | peg1933 | - | 100% aa identity to M.Kpn62629II |
| 11021 | G**A**TC | M.Kpn11021I (Dam) | 6mA | orphan | peg372 | - | 100% aa identity to M.Kpn43816Dam |
|  | C**C**WGG | M.Kpn11021II (Dcm) | 5mC | orphan | peg1937 | - | 100% aa identity to M.Kpn62629II |

*Note*: ^a^ The methylated nucleotide in the motif is shown as bold letter. The underlined letter represents the guanine pairing with the methylated cytosine on the complementary strand. Degenerate bases used in our recognition sequences are listed in the following: R = G or A, Y = C or T, M = A or C, K = G or T, S = G or C, W = A or T, B = not A (C or G or T), D = not C (A or G or T), H = not G (A or C or T), V = not T (A or C or G), N = A or C or G or T; ^b^ The MTase prediction was based on the sequence alignment with REBASE database (http://rebase.neb.com/rebase/rebase.html); The predicted MTases were further classified as Type I, Type II or orphan MTases according to the annotation information. ^c^ Some predicted MTases showed ~ 99.8%-100% identities with the known MTases as previously reported. “New” indicates the eight newly identified methylation motifs and corresponding MTases in our study.

### Table S11 Fisher’s exact test of the unmethylated and methylated MTCGAK sites

| **Strain No.** | **Sequence** | **Methylated motif** | **Un-methylated motif** | **Fisher’s exact test** | **Total motif** |
| --- | --- | --- | --- | --- | --- |
| 11492 | GMT**C**GAK | 476 | 1092 | p < 0.001 | 1574 |
|  | HMT**C**GAK | 2636 | 709 |  | 3400 |
| NTUH-K2044 | GMT**C**GAK | 349 | 1107 | p < 0.001 | 1546 |
|  | HMT**C**GAK | 1840 | 711 |  | 2551 |

*Note*: Degenerate bases used in the recognition sequences are listed in the following: M = A or C, K = G or T, H = not G (A or C or T).

### Table S12 Distribution of GATC motifs in gene region (GR) and intergenic region (IGR) among 14 *K. pneumoniae* strains

| **Strains No.** | **Methylated sites (GATC)** | | | | **Hemi-methylated sites (GATC)** | | | | **Un-methylated sites (GATC)** | | | |
| --- | --- | --- | --- | --- | --- | --- | --- | --- | --- | --- | --- | --- |
|  | **Total** | **GR** | **IGR** | **IGR (%)** | **Total** | **GR** | **IGR** | **IGR (%)** | **Total** | **GR** | **IGR** | **IGR (%)** |
| 11492 | 30271 | 28622 | 1649 | 5.45 | 31 | 26 | 5 | 16.13 | 14 | 3 | 11 | 78.57 |
| 11420 | 26835 | 25440 | 1395 | 5.20 | 4669 | 4376 | 293 | 6.28 | 343 | 292 | 51 | 14.87 |
| NTUH-K2044 | 30151 | 28861 | 1290 | 4.28 | 551 | 507 | 44 | 7.99 | 25 | 9 | 16 | 64.00 |
| 11454 | 28046 | 26620 | 1426 | 5.08 | 2142 | 1989 | 153 | 7.14 | 90 | 61 | 29 | 32.22 |
| 12208 | 30695 | 28993 | 1702 | 5.54 | 790 | 716 | 74 | 9.37 | 27 | 10 | 17 | 62.96 |
| 11311 | 29520 | 27967 | 1553 | 5.26 | 1076 | 999 | 77 | 7.16 | 27 | 15 | 12 | 44.44 |
| 23 | 26523 | 25120 | 1403 | 5.29 | 4423 | 4181 | 242 | 5.47 | 329 | 279 | 50 | 15.20 |
| N201205880 | 28007 | 26322 | 1685 | 6.02 | 4852 | 4574 | 278 | 5.73 | 312 | 267 | 45 | 14.42 |
| 11305 | 29222 | 27668 | 1554 | 5.32 | 1122 | 1033 | 89 | 7.93 | 25 | 16 | 9 | 36.00 |
| 721005 | 31521 | 28855 | 1301 | 4.31 | 687 | 1046 | 80 | 7.10 | 53 | 24 | 29 | 54.72 |
| 11021 | 27907 | 26435 | 1472 | 5.27 | 3968 | 3744 | 224 | 5.65 | 205 | 168 | 37 | 18.05 |
| 309074 | 30384 | 28841 | 1543 | 5.08 | 782 | 722 | 60 | 7.67 | 42 | 17 | 25 | 59.52 |
| 13190 | 28766 | 27199 | 1567 | 5.45 | 2515 | 2310 | 205 | 8.15 | 138 | 99 | 39 | 28.26 |
| 283747 | 30156 | 30122 | 1399 | 4.44 | 1126 | 638 | 49 | 7.13 | 34 | 13 | 21 | 61.76 |

### Table S13 Distribution of CCWGG motifs in gene region (GR) and intergenic region (IGR) among 14 *K. pneumoniae* strains

| **Strains No.** | **Methylated sites (CCWGG)** | | | | **Hemi-methylated sites (CCWGG)** | | | | **Un-methylated sites (CCWGG)** | | | |
| --- | --- | --- | --- | --- | --- | --- | --- | --- | --- | --- | --- | --- |
|  | **Total** | **GR** | **IGR** | **IGR (%)** | **Total** | **GR** | **IGR** | **IGR (%)** | **Total** | **GR** | **IGR** | **IGR (%)** |
| 11492 | 19015 | 17652 | 1407 | 7.38 | 86 | 77 | 9 | 10.47 | 183 | 181 | 2 | 1.09 |
| 11420 | 18828 | 17782 | 1461 | 7.59 | 839 | 778 | 61 | 7.27 | 508 | 472 | 36 | 7.09 |
| NTUH-K2044 | 16102 | 14842 | 1260 | 7.83 | 1755 | 1660 | 95 | 5.41 | 1673 | 1580 | 93 | 5.56 |
| 11454 | 14983 | 14864 | 1278 | 7.92 | 2367 | 2227 | 140 | 5.91 | 1964 | 1842 | 122 | 6.21 |
| 12208 | 19874 | 18336 | 1538 | 7.74 | 35 | 32 | 3 | 8.57 | 199 | 189 | 10 | 5.03 |
| 11311 | 19126 | 17786 | 1491 | 7.73 | 299 | 267 | 32 | 10.70 | 243 | 236 | 7 | 2.88 |
| 23 | 18668 | 17217 | 1451 | 7.77 | 963 | 903 | 60 | 6.23 | 310 | 298 | 12 | 3.87 |
| N201205880 | 20863 | 19165 | 1698 | 8.14 | 38 | 34 | 4 | 10.53 | 210 | 194 | 16 | 7.62 |
| 11305 | 18964 | 17647 | 1440 | 7.54 | 261 | 246 | 15 | 5.75 | 235 | 219 | 16 | 6.81 |
| 721005 | 20472 | 18897 | 1575 | 7.69 | 143 | 105 | 38 | 26.57 | 217 | 198 | 19 | 8.76 |
| 11021 | 18352 | 16887 | 1465 | 7.98 | 1358 | 1275 | 83 | 6.11 | 1026 | 967 | 59 | 5.75 |
| 309074 | 17464 | 16150 | 1314 | 7.52 | 1460 | 1378 | 82 | 5.62 | 1042 | 980 | 62 | 5.95 |
| 13190 | 19149 | 17785 | 1364 | 7.12 | 694 | 494 | 200 | 28.82 | 217 | 190 | 27 | 12.44 |
| 283747 | 19140 | 17691 | 1449 | 7.57 | 574 | 532 | 42 | 7.32 | 442 | 417 | 25 | 5.66 |

### Table S14 Summary of the genes with upstream hemi/un-methylated GATC sites shared in the 14 *K. pneumoniae* strains

| **Gene annotation** | **Location^a^** | ***K. pneumoniae* strains^b^** | | | | | | | | | | | | | |
| --- | --- | --- | --- | --- | --- | --- | --- | --- | --- | --- | --- | --- | --- | --- | --- |
|  |  | **K2044** | **11492** | **11420** | **11454** | **12208** | **11311** | **23** | **05880** | **11305** | **21005** | **11021** | **09074** | **13190** | **283747** |
| Hydroxyaromatic non-oxidative decarboxylase protein B | -75bp | **-/-** | **-/-** | **-/-** | **-/-** | **-/-** | **-/-** | **-/-** | **-/-** | **-/+** | **-/+** | **-/-** | **-/-** | **-/-** | **-/-** |
| PTS system, glucitol /sorbitol-specific IIC component | -85bp | **-/-** | **-/-** | **-/-** | **-/-** | **-/-** | **-/-** | **-/-** | **-/-** | **-/-** | **-/-** | **-/-** | **-/-** | **-/-** | **-/-** |
| PTS system, mannitol -specific IIA/B/C component | -157bp | **-/+** | **-/+** | **-/+** | **-/+** | **-/+** | **-/+** | **-/+** | **-/-** | **-/+** | **-/-** | **-/-** | **-/-** | **-/+** | **-/-** |
| Transcriptional regulator of arabinitol utilization, DeoR family | -141bp | **-/-** | **-/-** | **-/-** | **-/-** | **-/-** | **-/-** | **-/-** | **-/-** | **-/-** | **-/-** | **-/-** | **-/-** | **-/-** | **-/-** |
|  | -168bp | **-/+** | **-/-** | **-/-** | **-/-** | **-/-** | **-/-** | **-/-** | **-/-** | **-/-** | **-/-** | **-/-** | **-/-** | **-/-** | **-/-** |
| D-arabinitol 4-dehydrogenase | -41bp | **-/+** | **-/-** | **-/-** | **-/-** | **-/-** | **-/-** | **-/-** | **-/-** | **-/-** | **-/-** | **-/-** | **-/-** | **-/-** | **-/-** |
|  | -68bp | **-/-** | **-/-** | **-/-** | **-/-** | **-/-** | **-/-** | **-/-** | **-/-** | **-/-** | **-/-** | **-/-** | **-/-** | **-/-** | **-/-** |
| Antiholin-like protein LrgA | -61bp | **-/+** | **-/-** | **-/-** | **-/-** | **-/-** | **-/+** | **-/-** | **-/+** | **-/+** | **-/-** | **-/+** | **-/+** | **-/-** | **-/+** |
| C4-type zinc finger protein, DksA/TraR family | -30bp | **-/+** | **-/+** | **-/+** | **-/+** | **-/+** | **-/+** | **-/+** | **-/+** | **-/+** | **-/+** | **-/+** | **-/+** | **-/+** | **-/+** |
| Alpha-galactosidase | -37bp | **-/-** | **-/-** | **-/+** | **-/-** | **-/-** | **-/-** | **-/-** | **-/-** | **-/-** | **-/-** | **-/-** | **-/+** | **-/-** | **-/-** |
| Putative outer membrane protein | -59bp | **-/+** | **-/+** | **-/-** | **-/-** | **-/-** | **-/+** | **-/-** | **-/-** | **-/+** | **-/+** | **-/+** | **-/-** | **-/+** | **-/+** |
| Nudix hydrolase family protein YffH | -63bp | **-/+** | **-/-** | **-/-** | **-/-** | **-/+** | **-/+** | **-/+** | **-/-** | **-/+** | **-/+** | **-/+** | **-/+** | **-/-** | **-/+** |
| FIG00732740: hypothetical protein | -125bp | **-/-** | **-/-** | **-/-** | **-/-** | **-/-** | **-/-** | **-/-** | **-/-** | **-/-** | **-/-** | **-/-** | **-/-** | **-/-** | **-/-** |

*Note:* ^a^ The distance from the hemi/Un-methylated site to the start codon of the downstream gene; ^b^ ‘-/-’ denotes the upstream Un-methylated motifs on both strands. ‘-/+’ indicates the upstream hemi-methylated motif.

### Table S15 Summery of the genes with upstream hemi/Un-methylated GATC motif clusters in the 14 *K. pneumoniae* strains^a^

| **Genes** | **Strains^b^** | **Hemi/Un-methylated motifs^a^ (No.)** |
| --- | --- | --- |
| PTS system, mannitol-specific IIC component | 6 | 3-4 |
| tRNA uridine 5-carboxymethylaminomethyl modification enzyme GidA | 5 | 4-5 |
| Glycerol uptake facilitator protein | 4 | 4-6 |
| Leucine-responsive regulatory protein, regulator for leucine (or lrp) regulon and high-affinity branched-chain amino acid transport system | 3 | 3-8 |
| Putative membrane protein precursor | 3 | 3-8 |
| Ferredoxin | 3 | 3-4 |
| Outer membrane protein W precursor | 2 | 3-4 |
| GMP synthase [glutamine-hydrolyzing] | 2 | 4-6 |
| Putative transport protein | 2 | 3-5 |
| Transaldolase | 2 | 5-6 |
| ParB | 2 | 5-6 |
| Glutathione S-transferase, theta | 2 | 3-4 |

*Note:* ^a^ “Upstream hemi/Un-methylated GATC motif cluster” means that an upstream intergenic region contains at least three consecutive hemi/Un-methylated motifs; ^b^ “Strains” represents the strains with the hemi/Un-methylated GATC motif clusters.

### Table S16 Slow re-methylated GATC motifs in the 5’ USR of genes enriched in the four COG categories (“Cell cycle control, cell division, chromosome partitioning”, “Carbohydrate transport and metabolism”, “Translation, ribosomal structure and biogenesis” and “Intracellular trafficking, secretion, and vesicular transport”)

| **COG categories** | **Gene Name** | **IGR_SlowMotifSiteNumber** | **Distance to AUG** | **NTUH_TM** | **11492_TM** | **Gene Annotation** |
| --- | --- | --- | --- | --- | --- | --- |
| [C] Energy production and conversion | *aceB* | 1 | 243 | 16.254 | 13.434 | Malate synthase (EC 2.3.3.9) |
|  | *dhaD* | 1 | 106 | 9.2106 | 7.6126 | Glycerol dehydrogenase (EC 1.1.1.6) |
|  | *gatD* | 1 | 49 | 10.2942 | 8.5082 | Galactitol-1-phosphate 5-dehydrogenase (EC 1.1.1.251) |
|  | *mdh_1* | 1 | 177 | 12.4614 | 10.2994 | Malate dehydrogenase (EC 1.1.1.37) |
| [D] Cell cycle control, cell division, chromosome partitioning | *GidA* | 7 | 370;232;215;197;183;157;115 | 7.5852;12.4614;28.7154;19.5048;21.672;7.0434;13.0032 | 7.6126;23.7334;24.1812;9.8516;8.5082;19.2554;24.629 | tRNA uridine 5-carboxymethylaminomethyl modification enzyme GidA |
|  | *mreB* | 2 | 306;232 | 14.0868;12.4614 | 10.7472;10.7472 | Rod shape-determining protein MreB |
| [E] Amino acid transport and metabolism | *lysC* | 2 | 80;53 | 16.254;16.254 | 13.434;13.8818 | Aspartokinase (EC 2.7.2.4) |
|  | *pepQ* | 2 | 24;84 | 17.3376;17.3376 | 14.3296;14.3296 | Xaa-Pro dipeptidase PepQ (EC 3.4.13.9) |
|  | *argI* | 1 | 80 | 10.836 | 8.956 | Ornithine carbamoyltransferase (EC 2.1.3.3) |
|  | *aroQ* | 1 | 205 | 13.0032 | 10.7472 | 3-dehydroquinate dehydratase II (EC 4.2.1.10) |
|  | *DtpB* | 1 | 167 | 17.3376 | 13.8818 | Di/tripeptide permease DtpB |
|  | *gabR* | 1 | 5 | 16.254 | 18.3598 | Transcriptional regulator GabR of GABA utilization (GntR family with aminotransferase-like domain) |
|  | *mtr* | 1 | 103 | 10.836 | 8.956 | Tryptophan-specific transport protein |
|  | *patA* | 1 | 202 | 9.2106 | 8.0604 | Putrescine aminotransferase (EC 2.6.1.82) |
|  | *PepA* | 1 | 258 | 12.4614 | 8.956 | Cytosol aminopeptidase PepA (EC 3.4.11.1) |
|  | *prlC* | 1 | 59 | 16.7958 | 13.8818 | Oligopeptidase A (EC 3.4.24.70) |
|  | *sstT* | 1 | 148 | 9.7524 | 8.0604 | Sodium:dicarboxylate symporter |
|  | *tyrB_3* | 1 | 144 | 15.7122 | 12.9862 | Biosynthetic Aromatic amino acid aminotransferase alpha (EC 2.6.1.57) |
| [F] Nucleotide transport and metabolism | *nrdD* | 2 | 91;3 | 11.3778;17.3376 | 9.4038;9.4038 | Ribonucleotide reductase of class III (anaerobic), large subunit (EC 1.17.4.2) |
|  | *nudF* | 1 | 87 | 23.8392 | 19.2554 | ADP-ribose pyrophosphatase (EC 3.6.1.13) |
| [G] Carbohydrate transport and metabolism | *lsrA* | 4 | 145;92;60;40 | 15.1704;14.6286;16.7958;15.7122 | 17.4642;18.3598;19.2554;20.151 | Autoinducer 2 (AI-2) ABC transport system, fused AI2 transporter subunits and ATP-binding component |
|  | *gatY* | 2 | 70;92 | 10.2942;11.9196 | 15.673;8.5082 | Tagatose 1,6-bisphosphate aldolase (EC 4.1.2.40) |
|  | *mtlA* | 2 | 200;317 | 43.344;32.508 | 15.673;15.673 | PTS system mannitol-specific IIA/B/C component |
|  | *pgi* | 2 | 685;712 | 16.254;16.254 | 13.434;13.8818 | Glucose-6-phosphate isomerase (EC 5.3.1.9) |
|  | *rafA* | 2 | 36;73 | 52.0128;31.9662 | 42.9888;16.5686 | Alpha-galactosidase (EC 3.2.1.22) |
|  | *RbsA* | 2 | 162;110 | 15.1704;14.0868 | 11.6428;11.6428 | ABC transport system, ATP-binding protein Z5691 |
|  | *dhaK_2* | 1 | 145 | 9.2106 | 7.6126 | Phosphoenolpyruvate-dihydroxyacetone phosphotransferase (EC 2.7.1.121), dihydroxyacetone binding subunit DhaK |
|  | *garD* | 1 | 111 | 10.2942 | 17.912 | D-galactarate dehydratase (EC 4.2.1.42) |
|  | *gatZ* | 1 | 76 | 10.2942 | 8.5082 | Tagatose-6-phosphate kinase GatZ (EC 2.7.1.144) |
|  | *gmhA* | 1 | 4 | 10.836 | 12.0906 | Phosphoheptose isomerase (EC 5.3.1.-) |
|  | *lamB_4* | 1 | 14 | 16.7958 | 12.9862 | Maltoporin (maltose/maltodextrin high-affinity receptor, phage lambda receptor protein) |
|  | *licH_2* | 1 | 45 | 12.4614 | 9.4038 | 6-phospho-beta-glucosidase (EC 3.2.1.86) |
|  | *malF_2* | 1 | 6 | 16.254 | 12.9862 | Maltose/maltodextrin ABC transporter, permease protein MalF |
|  | *msmF* | 1 | 245 | 8.127 | 22.39 | Multiple sugar ABC transporter, membrane-spanning permease protein MsmF |
|  | *srlA_1* | 1 | 84 | 28.1736 | 11.6428 | PTS system, glucitol/sorbitol-specific IIC component (EC 2.7.1.69) |
|  | *tagK* | 1 | 16 | 10.2942 | 8.956 | Tagatose-1-phosphate kinase TagK |
|  | *treB* | 1 | 108 | 14.6286 | 10.2994 | PTS system, trehalose-specific IIB component (EC 2.7.1.69) / PTS system, trehalose-specific IIC component (EC 2.7.1.69) |
|  | *ugpB* | 1 | 196 | 16.7958 | 17.912 | Glycerol-3-phosphate ABC transporter, periplasmic glycerol-3-phosphate-binding protein (TC 3.A.1.1.3) |
|  | *uxaC* | 1 | 110 | 9.7524 | 8.0604 | Uronate isomerase (EC 5.3.1.12) |
|  | *uxuA_1* | 1 | 88 | 11.9196 | 9.8516 | Mannonate dehydratase (EC 4.2.1.8) |
|  | *xynT* | 1 | 112 | 9.2106 | 17.0164 | Xyloside transporter XynT |
|  | *yjgK* | 1 | 145 | 10.836 | 8.956 | Protein YjgK, linked to biofilm formation |
| [H] Coenzyme transport and metabolism | *thiC* | 3 | 140;126;62 | 17.8794;16.7958;16.7958 | 13.8818;13.8818;13.8818 | Hydroxymethylpyrimidine phosphate synthase ThiC (EC 4.1.99.17) |
|  | *ispB* | 2 | 46;147 | 11.3778;11.3778 | 9.4038;9.4038 | Octaprenyl diphosphate synthase (EC 2.5.1.90) |
|  | *RibB* | 2 | 348;158 | 8.6688;8.6688 | 9.4038;7.1648 | 3,4-dihydroxy-2-butanone 4-phosphate synthase (EC 4.1.99.12) |
|  | *ecdB* | 1 | 74 | 25.4646 | 27.3158 | Hydroxyaromatic non-oxidative decarboxylase protein B (EC 4.1.1.-) |
|  | *ubiC* | 1 | 79 | 15.7122 | 12.9862 | Chorismate--pyruvate lyase (EC 4.1.3.40) |
| [I] Lipid transport and metabolism | *fadB* | 2 | 168;108 | 17.3376;17.3376 | 14.3296;14.3296 | Enoyl-CoA hydratase (EC 4.2.1.17) / Delta(3)-cis-delta(2)-trans-enoyl-CoA isomerase (EC 5.3.3.8) / 3-hydroxyacyl-CoA dehydrogenase (EC 1.1.1.35) / 3-hydroxybutyryl-CoA epimerase (EC 5.1.2.3) |
|  | *acs* | 1 | 94 | 14.6286 | 12.9862 | Acetyl-coenzyme A synthetase (EC 6.2.1.1) |
|  | *psd* | 1 | 161 | 13.0032 | 10.7472 | Phosphatidylserine decarboxylase (EC 4.1.1.65) |
| [J] Translation, ribosomal structure and biogenesis | *dusB* | 3 | 56;192;378 | 13.0032;13.0032;13.0032 | 10.7472;13.8818;10.7472 | tRNA dihydrouridine synthase B (EC 1.-.-.-) |
|  | *L22p* | 2 | 176;161 | 13.545;13.545 | 13.434;11.195 | LSU ribosomal protein L22p (L17e) |
|  | *rlmG* | 2 | 76;69 | 15.7122;9.7524 | 8.0604;9.8516 | 23S rRNA (guanine-N-2-) -methyltransferase rlmG (EC 2.1.1.-) |
|  | *rpsJ* | 2 | 367;299 | 13.545;13.545 | 15.673;11.195 | SSU ribosomal protein S10p (S20e) |
|  | *EF1A/EF-Tu* | 1 | 723 | 17.3376 | 14.3296 | Translation elongation factor Tu |
|  | *L13p* | 1 | 189 | 13.0032 | 12.0906 | LSU ribosomal protein L13p (L13Ae) |
|  | *L14p* | 1 | 15 | 13.545 | 11.195 | LSU ribosomal protein L14p (L23e) |
|  | *L17p* | 1 | 36 | 13.545 | 11.195 | LSU ribosomal protein L17p |
|  | *L28p* | 1 | 229 | 28.1736 | 23.2856 | LSU ribosomal protein L28p |
|  | *rplK* | 1 | 148 | 13.0032 | 14.3296 | LSU ribosomal protein L11p (L12e) |
|  | *rpsL* | 1 | 117 | 13.545 | 11.6428 | SSU ribosomal protein S12p (S23e) |
|  | *rpsO* | 1 | 40 | 10.836 | 8.956 | SSU ribosomal protein S15p (S13e) |
| [K] Transcription | *LsrR* | 4 | 26;79;111;131 | 15.1704;14.6286;16.7958;15.7122 | 17.4642;18.3598;19.2554;20.151 | LsrR, transcriptional repressor of lsr operon |
|  | *ExuR* | 2 | 103;126 | 9.7524;9.7524 | 8.0604;8.0604 | Hexuronate utilization operon transcriptional repressor ExuR |
|  | *argR* | 1 | 238 | 12.4614 | 10.2994 | Arginine pathway regulatory protein ArgR, repressor of arg regulon |
|  | *ccdA* | 1 | 167 | 18.4212 | 12.5384 | Transcriptional regulator of arabinitol utilization, DeoR family |
|  | *QseA* | 1 | 47 | 12.4614 | 10.2994 | LysR family transcriptional regulator QseA |
|  | *rpoD* | 1 | 122 | 8.6688 | 7.1648 | RNA polymerase sigma factor RpoD |
| [L] Replication, recombination and repair | *dnaA* | 2 | 423;434 | 21.1302;17.8794 | 20.151;8.0604 | Chromosomal replication initiator protein DnaA |
|  | *parC* | 2 | 96;11 | 8.127;8.127 | 8.0604;9.4038 | Topoisomerase IV subunit A (EC 5.99.1.-) |
|  | *dnaB_2* | 1 | 18 | 15.7122 | 12.9862 | Replicative DNA helicase (EC 3.6.1.-) |
|  | *RsmD* | 1 | 30 | 16.254 | 16.1208 | 16S rRNA (guanine(966)-N(2))-methyltransferase (EC 2.1.1.171) |
|  | *smc_3* | 1 | 351 | 13.545 | 12.5384 | putative exonuclease |
|  | *TDG* | 1 | 584 | 8.6688 | 7.1648 | G:T/U mismatch-specific uracil/thymine DNA-glycosylase |
| [M] Cell wall/membrane/envelope biogenesis | *murB* | 7 | 1748;2948;3721;4140;4314;4965;5389 | 17.3376;17.3376;17.3376;17.3376;17.3376;27.09;17.3376 | 14.3296;16.5686;14.3296;14.3296;14.3296;15.2252;14.3296 | UDP-N-acetylenolpyruvoylglucosamine reductase (EC 1.1.1.158) |
|  | *blc_2* | 1 | 114 | 13.0032 | 10.7472 | Outer membrane lipoprotein Blc |
|  | *mltB* | 1 | 193 | 28.1736 | 11.6428 | Membrane-bound lytic murein transglycosylase B precursor (EC 3.2.1.-) |
|  | *murA* | 1 | 43 | 11.3778 | 9.4038 | UDP-N-acetylglucosamine 1-carboxyvinyltransferase (EC 2.5.1.7) |
| [O] Posttranslational modification, protein turnover, chaperones | *fkpA* | 1 | 203 | 14.0868 | 11.6428 | FKBP-type peptidyl-prolyl cis-trans isomerase FkpA precursor (EC 5.2.1.8) |
|  | *nrdG* | 1 | 240 | 11.3778 | 9.4038 | Ribonucleotide reductase of class III (anaerobic), activating protein (EC 1.97.1.4) |
|  | *ppiA* | 1 | 120 | 14.0868 | 19.7032 | Peptidyl-prolyl cis-trans isomerase PpiA precursor (EC 5.2.1.8) |
| [P] Inorganic ion transport and metabolism | *MgtE* | 2 | 63;233 | 16.7958;16.7958 | 13.8818;15.673 | Mg/Co/Ni transporter MgtE / CBS domain |
|  | *PhnA* | 2 | 339;271 | 14.0868;14.0868 | 11.6428;11.6428 | Alkylphosphonate utilization operon protein PhnA |
|  | *mgtA* | 1 | 188 | 10.836 | 8.956 | Mg(2+) transport ATPase, P-type (EC 3.6.3.2) |
|  | *TerC* | 1 | 177 | 9.7524 | 8.0604 | Integral membrane protein TerC |
| [Q] Secondary metabolites biosynthesis, transport and catabolism | *dhaB* | 1 | 210 | 14.6286 | 20.151 | Glycerol dehydratase large subunit (EC 4.2.1.30) |
| [R] General function prediction only | *ghxP* | 2 | 73;376 | 14.6286;14.6286 | 12.0906;12.0906 | Guanine-hypoxanthine permease |
|  | *dcuB* | 1 | 27 | 16.7958 | 13.8818 | C4-dicarboxylate transporter DcuB |
|  | *FxsA* | 1 | 58 | 13.545 | 11.195 | FxsA protein |
|  | *qorA_2* | 1 | 152 | 15.7122 | 12.9862 | Quinone oxidoreductase (EC 1.6.5.5) |
|  | *yffH* | 1 | 62 | 21.672 | 18.3598 | Nudix hydrolase family protein YffH |
|  | *yhhW_1* | 1 | 55 | 15.7122 | 12.9862 | Qercetin 2,3-dioxygenase |
|  | *ytfJ* | 1 | 32 | 15.1704 | 9.8516 | Protein ytfJ precursor |
| [S] Function unknown | *iolC* | 3 | 197;158;46 | 10.2942;10.2942;10.2942 | 8.5082;8.5082;8.5082 | 5-keto-2-deoxygluconokinase (EC 2.7.1.92) / uncharacterized domain |
|  | *psiE* | 3 | 31;53;59 | 15.7122;15.7122;15.7122 | 12.9862;12.9862;12.9862 | PsiE protein |
|  | *NusA* | 2 | 147;135 | 11.3778;11.3778 | 8.956;11.195 | FIG000325: clustered with transcription termination protein NusA |
|  | *rraB* | 1 | 84 | 10.836 | 8.956 | Ribonuclease E inhibitor RraB |
|  | *slyX* | 1 | 20 | 14.0868 | 11.6428 | Protein SlyX |
|  | *YqcC* | 1 | 517 | 15.7122 | 17.912 | Hypothetical protein YqcC (clustered with tRNA pseudouridine synthase C) |
|  | *yqjF* | 1 | 133 | 10.2942 | 8.5082 | Inner membrane protein YqjF |
| [T] Signal transduction mechanisms | *arcB* | 1 | 135 | 11.9196 | 9.8516 | Aerobic respiration control sensor protein arcB (EC 2.7.3.-arcB |
|  | *ompR* | 1 | 37 | 14.6286 | 12.0906 | Response regulator protein Z5684 |
| [U] Intracellular trafficking, secretion, and vesicular transport | *pilD* | 2 | 17;85 | 13.545;13.545 | 15.673;11.195 | Leader peptidase (Prepilin peptidase) (EC 3.4.23.43) / N-methyltransferase (EC 2.1.1.-) |
|  | *pilQ* | 2 | 298;234 | 14.6286;16.254 | 12.0906;14.3296 | Type IV pilus biogenesis protein PilQ |
|  | *ExbD/TolR* | 1 | 7 | 7.5852 | 8.5082 | Biopolymer transport protein ExbD/TolR |
|  | *ftsY* | 1 | 141 | 16.254 | 16.1208 | Signal recognition particle receptor protein FtsY (alpha subunit) (TC 3.A.5.1.1) |
|  | *secE* | 1 | 56 | 17.3376 | 14.3296 | Preprotein translocase subunit SecE (TC 3.A.5.1.1) |
|  | *secG* | 1 | 180 | 17.3376 | 21.0466 | Preprotein translocase subunit SecG (TC 3.A.5.1.1) |

*Note:* Genes in red fonts indicate the genes with hemi/un-methylated sites in 5’ USRs shared in 14 *K. pneumoniae* strains. Genes with yellow background color indicate the genes with hemi/un-methylated clusters in 5’ USRs.

### Table S17 Distribution of newly reported motifs in GRs and IGRs among seven *K. pneumoniae* strains

| **Strains No.** | **Motif** | **Methylated sites** | | | | **Hemi-methylated sites** | | | | **Un-methylated sites** | | | |
| --- | --- | --- | --- | --- | --- | --- | --- | --- | --- | --- | --- | --- | --- |
|  |  | **Total** | **GR** | **IGR** | **IGR (%)** | **Total** | **GR** | **IGR** | **IGR (%)** | **Total** | **GR** | **IGR** | **IGR (%)** |
| NTUH-K2044 | GRACRAC^*^ | 2,060 | 1,846 | 214 | 10.39 | / | / | / | / | 44 | 41 | 3 | 6.82 |
| 11420 | RTACN_5_GGC^*^ | 1,156 | 1,027 | 129 | 11.16 | / | / | / | / | 119 | 106 | 13 | 6.82 |
|  | TTCAN_7_TTC^*^ | 598 | 524 | 74 | 12.37 | 231 | 199 | 32 | 13.85 | 24 | 19 | 5 | 20.83 |
| 12208 | AGGAAG^*^ | 2,848 | 2,495 | 353 | 12.39 | / | / | / | / | 16 | 13 | 3 | 18.75 |
| 11311 | CCAYN_7_TTYG^*^ | 608 | 557 | 51 | 8.39 | 58 | 44 | 14 | 24.13 | 1 | 1 | 0 | 0 |
| 23 | CCAYN_7_TTYG^*^ | 522 | 478 | 44 | 8.42 | 156 | 131 | 25 | 16.03 | 9 | 9 | 0 | 0 |
| N201205880 | CCAN_7_TCAC^*^ | 433 | 383 | 50 | 11.55 | 102 | 83 | 19 | 18.63 | 8 | 5 | 3 | 37.5 |
| 309074 | CAGN_6_TCAA^*^ | 413 | 349 | 64 | 15.50 | 26 | 17 | 9 | 34.62 | / | / | / | / |
| NTUH-K2044 | MT**C**GAK^*^ | 2,340 | 2,257 | 83 | 3.55 | 268 | 249 | 19 | 7.09 | 2,803 | 2,591 | 212 | 7.56 |
| 11492 | MT**C**GAK^*^ | 2,368 | 2,268 | 100 | 4.22 | 258 | 231 | 27 | 10.47 | 2,760 | 2,568 | 192 | 6.96 |

### Table S18 Summary of the genes with upstream hemi/un-methylated CCAYN_7_TTYG sites shared in the 23 and 11311 *K. pneumoniae* strains.

| **Gene annotation** | **Location^a^** | ***K. pneumoniae* strains^b^** | |
| --- | --- | --- | --- |
|  |  | **23** | **11311** |
| 6-phospho-beta-glucosidase (EC 3.2.1.86) | -151bp | **-/+** | **-/+** |
| Deoxyribose-phosphate aldolase (EC 4.1.2.4) | -350bp | **-/+** | **-/+** |
| Iron(III) dicitrate transport protein FecA | -10bp | **-/+** | **-/+** |

*Note:* ^a^ The distance from the hemi/Un-methylated site to the start codon of the downstream gene; ^b^ ‘-/-’ denotes the upstream Un-methylated motifs on both strands. ‘-/+’ indicates the upstream hemi-methylated motif.

### Table S19 Summary of the genes with upstream hemi/un-methylated MTCGAK sites shared in the NUTH-K2044 and 11492 *K. pneumoniae* strains.

| **Gene annotation** | **Location^a^** | ***K. pneumoniae* strains^b^** | |
| --- | --- | --- | --- |
|  |  | NUTH-2044 | 11492 |
| hypothetical protein | -238bp | -/- | -/- |
| Purine nucleotide synthesis repressor | -40bp | -/- | -/- |
| LSU ribosomal protein L10p (P0) | -310bp | -/- | -/- |
| 6-phosphofructokinase (EC 2.7.1.11) | -214bp | -/- | -/- |
| Type I secretion outer membrane protein%2C TolC precursor | -66bp | -/- | -/- |
| Translation elongation factor Tu | -1bp | -/- | -/- |
| hypothetical protein | -13bp | -/- | -/+ |
| putative membrane permeability altering protein | -213bp | -/- | -/- |
| probable fimbrial protein staA | -42bp | -/- | -/- |
| tRNA-Leu-CAG | -9bp | -/- | -/- |
| LSU ribosomal protein L28p | -65bp | -/+ | -/- |
| hypothetical protein | -1bp | -/- | -/- |
| Cystathionine beta-synthase (EC 4.2.1.22) | -146bp | -/- | -/- |
| Phosphotransferase system%2C phosphocarrier protein HPr | -208bp | -/- | -/- |
| HTH-type transcriptional regulator YidP | -75bp | -/- | -/- |
| NAD-dependent protein deacetylase of SIR2 family | -77bp | -/- | -/- |
| Aspartate-semialdehyde dehydrogenase (EC 1.2.1.11) | -93bp | -/- | -/- |
| Ribitol operon repressor | -57bp | -/- | -/- |
| Putative sulfate permease | -149bp | -/- | -/- |
| Respiratory nitrate reductase alpha chain (EC 1.7.99.4) | -141bp | -/- | -/- |
| Pca regulon regulatory protein PcaR | -24bp | -/- | -/- |
| Multimodular transpeptidase-transglycosylase (EC 2.4.1.129) (EC 3.4.-.-) | -67bp | -/- | -/- |
| Protein rarD | -37bp | -/- | -/- |
| FIG00622692: hypothetical protein | -43bp | -/+ | -/+ |
| Thiamin biosynthesis lipoprotein ApbE | -6bp | -/- | -/- |
| hypothetical protein | -513bp | -/- | -/- |
| Thioredoxin 2 (EC 1.8.1.8) | -69bp | -/- | -/- |
| 2-oxoglutarate dehydrogenase E1 component (EC 1.2.4.2) | -250bp | -/- | -/+ |
| Aerobic glycerol-3-phosphate dehydrogenase (EC 1.1.5.3) | -42bp | -/- | -/- |
| Para-aminobenzoate synthase%2C aminase component (EC 2.6.1.85) | -7bp | -/- | -/- |
| Membrane fusion protein of RND family multidrug efflux pump | -12bp | -/- | -/- |
| hypothetical protein | -138bp | -/- | -/- |
| Glucans biosynthesis protein G precursor | -35bp | -/- | -/- |
| putative alpha helix protein | -15bp | -/- | -/- |
| tRNA dihydrouridine synthase B (EC 1.-.-.-) | -301bp | -/- | -/- |
| Na+/H+ antiporter NhaA type | -30bp | -/- | -/- |
| Cellulose synthase (UDP-forming) (EC 2.4.1.12) | -51bp | -/- | -/- |
| Ureidoglycolate dehydrogenase (EC 1.1.1.154) | -149bp | -/- | -/- |
| Deoxyribose-phosphate aldolase (EC 4.1.2.4) | -244bp | -/- | -/- |
| 2-acylglycerophosphoethanolamine acyltransferase (EC 2.3.1.40) / Acyl-[acyl-carrier-protein] synthetase (EC 6.2.1.20) | -180bp | -/+ | -/+ |
| Deoxyguanosinetriphosphate triphosphohydrolase (EC 3.1.5.1) | -178bp | -/- | -/- |
| HTH-type transcriptional regulator YidP | -119bp | -/- | -/- |
| FIG00732100: hypothetical protein | -314bp | -/- | -/- |
| 5-methylthioribose kinase (EC 2.7.1.100) | -7bp | -/- | -/- |
| Negative regulator of allantoin and glyoxylate utilization operons | -5bp | -/- | -/- |
| Mg/Co/Ni transporter MgtE / CBS domain | -2bp | -/- | -/- |
| Conidiation-specific protein 10 | -72bp | -/- | -/- |
| Inosose dehydratase (EC 4.2.1.44) | -137bp | -/- | -/- |
| hypothetical protein | -886bp | -/- | -/- |
| Thermostable carboxypeptidase 1 (EC 3.4.17.19) | -184bp | -/- | -/- |
| Cobalamin synthase (EC 2.7.8.26) | -83bp | -/- | -/- |
| Ribonucleotide reductase of class Ia (aerobic)%2C alpha subunit (EC 1.17.4.1) | -267bp | -/- | -/- |
| putative exonuclease | -28bp | -/- | -/- |
| Transcriptional regulator%2C LysR family | -20bp | -/- | -/- |
| Transcriptional regulator YbiH%2C TetR family | -26bp | -/- | -/- |
| hypothetical protein | -44bp | -/- | -/- |
| putative plasmid-related protein | -44bp | -/- | -/- |
| FIG00731333: hypothetical protein | -24bp | -/- | -/- |
| FIG00732152: hypothetical protein | -150bp | -/- | -/- |
| 6-phospho-beta-glucosidase (EC 3.2.1.86) | -32bp | -/- | -/- |
| Putative cation-transporting P-type ATPase | -12bp | -/- | -/- |
| Mgl repressor and galactose ultrainduction factor GalS%2C HTH-type transcriptional regulator | -149bp | -/- | -/- |
| Flavodoxin 2 | -14bp | -/- | -/- |
| Uncharacterized oxidoreductase ydgJ (EC 1.-.-.-) | -150bp | -/- | -/- |
| tRNA-Met-CAT | -85bp | -/- | -/- |
| Regulatory protein SoxS | -23bp | -/- | -/+ |
| hypothetical protein | -8bp | -/- | -/+ |
| Coenzyme PQQ synthesis protein B | -102bp | -/- | -/- |
| Regulatory protein RecX | -10bp | -/+ | -/+ |
| hypothetical protein | -151bp | -/- | -/- |
| Ribonuclease HI (EC 3.1.26.4) | -29bp | -/- | -/- |
| FIG074102: hypothetical protein | -40bp | -/- | -/- |
| tRNA-Gln-CTG | -23bp | -/- | -/+ |
| ABC-type Fe3+-siderophore transport system%2C permease component | -44bp | -/- | -/- |
| Ribonuclease E (EC 3.1.26.12) | -273bp | -/- | -/- |
| HTH-type transcriptional regulator YidP | -13bp | -/- | -/+ |
| hypothetical protein | -17bp | -/- | -/- |
| Transcriptional regulator%2C GntR family | -25bp | -/- | -/- |
| probable membrane protein yjeI | -16bp | -/- | -/- |
| hypothetical protein | -112bp | -/- | -/- |
| putative exported protein | -239bp | -/- | -/- |
| Diguanylate cyclase/phosphodiesterase domain 2 (EAL) | -355bp | -/- | -/- |
| Transcriptional regulator%2C LysR family | -1bp | -/- | -/- |
| Nucleoside permease NupG | -92bp | -/- | -/- |
| FIG00731430: hypothetical protein | -128bp | -/- | -/- |
| C4-type zinc finger protein%2C DksA/TraR family | -27bp | -/- | -/- |
| Iron binding protein SufA for iron-sulfur cluster assembly | -2bp | -/+ | -/+ |
| tRNA-Leu-CAG | -93bp | -/- | -/- |
| Putative regulator protein | -83bp | -/- | -/- |
| Carbonic anhydrase (EC 4.2.1.1) | -26bp | -/- | -/- |
| PTS system%2C mannose-specific IIA component (EC 2.7.1.69) / PTS system%2C mannose-specific IIB component (EC 2.7.1.69) | -73bp | -/- | -/- |
| FIG00731878: hypothetical protein | -171bp | -/- | -/- |
| Quinate/shikimate dehydrogenase [Pyrroloquinoline-quinone] (EC 1.1.99.25) | -146bp | -/- | -/- |
| DNA helicase IV | -8bp | -/- | -/- |
| Shikimate transporter | -141bp | -/- | -/- |
| Small Subunit Ribosomal RNA%3B ssuRNA%3B SSU rRNA | -355bp | -/- | -/- |
| Sulfate and thiosulfate binding protein CysP | -342bp | -/- | -/- |
| Transcriptional regulator GabR of GABA utilization (GntR family with aminotransferase-like domain) | -32bp | -/- | -/- |
| LysR-family transcriptional regulator STM3020 | -18bp | -/- | -/- |
| Leader peptidase (Prepilin peptidase) (EC 3.4.23.43) / N-methyltransferase (EC 2.1.1.-) | -19bp | -/- | -/- |
| Protein ImpG/VasA | -173bp | -/- | -/- |
| Dipeptide-binding ABC transporter%2C periplasmic substrate-binding component (TC 3.A.1.5.2) | -83bp | -/+ | -/+ |
| hypothetical protein | -312bp | -/- | -/- |
| Sensory histidine kinase in two-component regulatory system with RstA | -192bp | -/- | -/- |
| Succinate dehydrogenase flavoprotein subunit (EC 1.3.99.1) | -126bp | -/- | -/- |
| Nucleoside permease NupG | -61bp | -/- | -/- |
| tmRNA-binding protein SmpB | -37bp | -/- | -/- |
| Ribitol/Xylitol/Arabitol transporter%2C MFS superfamily | -9bp | -/+ | -/+ |
| Argininosuccinate synthase (EC 6.3.4.5) | -17bp | -/- | -/+ |
| Glutathione S-transferase%2C theta (EC 2.5.1.18) | -116bp | -/- | -/- |
| Transcriptional regulator%2C IclR family | -30bp | -/- | -/- |
| PTS system%2C diacetylchitobiose-specific IIA component (EC 2.7.1.69) | -2bp | -/- | -/+ |
| UDP-N-acetylglucosamine 2-epimerase (EC 5.1.3.14) | -48bp | -/- | -/- |
| tRNA-Tyr-GTA | -123bp | -/- | -/- |
| hypothetical protein | -91bp | -/- | -/- |
| DNA polymerase I (EC 2.7.7.7) | -49bp | -/- | -/- |
| Ribosome hibernation protein YfiA | -258bp | -/- | -/- |
| RNA polymerase sigma factor RpoD | -124bp | -/- | -/- |
| tRNA-Arg-ACG | -86bp | -/- | -/- |
| C-terminal domain of CinA paralog%2C YdeJ | -66bp | -/- | -/- |
| Nitrate/nitrite transporter | -46bp | -/- | -/- |
| hypothetical protein | -26bp | -/- | -/- |
| 5S RNA | -35bp | -/- | -/- |
| Calcium/proton antiporter | -106bp | -/- | -/- |
| Response regulator protein Z5684 | -39bp | -/- | -/- |
| tRNA-Phe-GAA | -63bp | -/- | -/- |
| FIG00732313: hypothetical protein | -132bp | -/- | -/- |
| Ribitol operon repressor | -85bp | -/- | -/- |
| hypothetical protein | -32bp | -/- | -/- |
| LysR family transcriptional regulator YeiE | -59bp | -/- | -/- |
| Beta-lactamase-like | -81bp | -/- | -/- |
| PTS system%2C fructose-specific IIA component (EC 2.7.1.69) | -39bp | -/- | -/- |
| Ribosomal-protein-S5p-alanine acetyltransferase | -86bp | -/- | -/- |
| Nucleoside permease NupC | -128bp | -/- | -/- |
| hypothetical protein | -74bp | -/- | -/- |
| O-antigen export system permease protein RfbD | -90bp | -/- | -/- |
| putative exonuclease | -218bp | -/- | -/- |
| Probable secreted protein | -13bp | -/- | -/+ |
| FIG00731568: hypothetical protein | -69bp | -/- | -/- |
| Intracellular septation protein IspA | -6bp | -/- | -/- |
| Cold shock protein CspC | -240bp | -/- | -/- |
| FIG074102: hypothetical protein | -99bp | -/- | -/- |
| UPF0246 protein YaaA | -59bp | -/- | -/- |
| ABC transport system%2C permease component YbhR | -80bp | -/- | -/+ |
| Putative two-domain glycosyltransferase | -60bp | -/- | -/- |
| Zinc ABC transporter%2C ATP-binding protein ZnuC | -61bp | -/- | -/- |

*Note:* ^a^ The distance from the hemi/Un-methylated site to the start codon of the downstream gene; ^b^ ‘-/-’ denotes the upstream Un-methylated motifs on both strands. ‘-/+’ indicates the upstream hemi-methylated motif.
